# Systematic clustering alignment and feature characterization for single-cell omics using ACE-OF-Clust

**DOI:** 10.64898/2026.03.09.710668

**Authors:** Xiran Liu, Ritambhara Singh, Sohini Ramachandran

## Abstract

Clustering is widely used to identify cell types in cellular-resolution transcriptomic data, including single-cell RNA sequencing (scRNA-seq) and spatial transcriptomics (ST). Mixed-membership clustering assigns fractional memberships across clusters and captures continuous variation beyond hard clustering, but integrating and interpreting results from either approach is complicated by the “clustering alignment problem,” which arises from label switching, multi-modality, and differences in model settings (including differing numbers of clusters). We introduce `ACE-OF-Clust`, enabling a four-step workflow for single-cell clustering: multiple clustering, clustering alignment, model comparison, and identification of informative features. `ACE-OF-Clust` introduces direct comparison of clustering solutions, assesses consistency against annotations, and leverages feature-level clustering profiles to prioritize genes discriminating among cell types, as well as to evaluate how gene sets contribute to clustering patterns. We demonstrate its utility on PBMC scRNA-seq and breast cancer ST data, and on multi-omic single-cell data. `ACE-OF-Clust` quantifies cross-omic clustering variability and suggests putative cross-omic regulatory links. Overall, `ACE-OF-Clust` increases the interpretability, flexibility, and robustness of single-cell clustering, providing a scalable tool for studying cellular heterogeneity and gene expression dynamics.

## 1 Introduction

Computational cell type identification in scRNA-seq groups individual cells by their transcriptional profiles to reveal functional and molecular heterogeneity (Wagner et al., 2016; Kiselev et al., 2019), enabling resolution of neuronal (Macosko et al., 2015; Zeisel et al., 2015), immune (Villani et al., 2017; G. X. Zheng et al., 2017), and tumor subtypes (Puram et al., 2017; F. Wu et al., 2021). This quantitative task supports characterization of cellular diversity, detection of rare populations, and investigation of how cell types contribute to development, disease, and tissue homeostasis, as well as the determinants and variability of cell identity (Wagner et al., 2016; Zeng, 2022). Spatial transcriptomics (ST) extends these analyses by measuring gene expression while preserving spatial organization (Crosetto et al., 2015; Cheng et al., 2023): each “spot” provides a transcriptome and tissue coordinates and is commonly treated as a proxy “cell” despite often capturing multiple cells (Cheng et al., 2023). The spatial context enables identification of spatially structured subpopulations, characterization of tissue transcriptional architecture, and study of subcellular RNA localization with functional implications (Crosetto et al., 2015; Cheng et al., 2023; J. Du et al., 2023).

Hard clustering approaches which partition individual cells into discrete groups based on gene expression are central to cell type identification in scRNA-seq and ST analyses. *Hard clustering* assigns each cell to a single cluster and is widely used to resolve transcriptionally distinct subpopulations (Kiselev et al., 2019; Petegrosso et al., 2020; Qi et al., 2020; S. Zhang et al., 2023; Sun et al., 2024). Popular pipelines in `Scanpy` (Wolf et al., 2018) and `Seurat` (Satija et al., 2015; Butler et al., 2018) typically apply dimensionality reduction (e.g., PCA) followed by clustering (e.g., k-means or community detection) (Kiselev et al., 2019; Petegrosso et al., 2020; Qi et al., 2020), and have supported cell-type discovery and downstream validation in many scRNA-seq studies (Macosko et al., 2015; Zeisel et al., 2015; Puram et al., 2017; Villani et al., 2017; G. X. Zheng et al., 2017; F. Wu et al., 2021). Similar models are often used in ST, sometimes alongside probabilistic approaches such as Gaussian mixture modeling (Long et al., 2023), and they also serve as benchmarks for spatial clustering methods that incorporate spatial context before applying nonspatial clustering (Hu et al., 2021; E. Zhao et al., 2021; Long et al., 2023; Xu et al., 2024; Yuan et al., 2024). However, because hard clustering enforces mutually exclusive assignments, it cannot capture continuous variation such as transient states or spatial gradients (Wagner et al., 2016; Dey et al., 2017; Kiselev et al., 2019), motivating the use of *mixed-membership clustering* to better reflect the dynamic, continuous nature of cell types in functional genomics.

In contrast to hard clustering, *mixed-membership clustering* (also known as soft or fuzzy clustering) allows each data point to have fractional memberships across, typically interpreted as probabilities or proportions of cluster membership. These memberships are often represented by a matrix *Q*, with entries *q*_*ik*_ denoting probabilistic membership assignments that satisfy 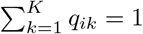 (see Methods). Mixed-membership clustering has been applied in scRNA-seq (Dey et al., 2017; Bravo González-Blas et al., 2019; H.-J. Kim et al., 2020; Carbonetto, Sarkar, et al., 2021; Pancheva et al., 2022; Carbonetto, K. Luo, et al., 2023; Malagoli et al., 2024) and can capture continuous variation, including hybrid or intermediate states. Many such clustering models are based on topic modeling or non-negative matrix factorization and additionally infer a feature-level matrix *P* (see Methods), often interpreted as relative gene expression per cluster, which enables identification of genes driving clustering heterogeneity (Dey et al., 2017; Pancheva et al., 2022; Carbonetto, K. Luo, et al., 2023). The availability of *P* provides an additional advantage: gene-level signals are inferred jointly with clusters, mitigating the double-dipping (double use of data) issue that can arise in post hoc differential testing (Lähnemann et al., 2020). Moreover, clustering is not limited to RNA-seq gene expression; it is also a core step for analyzing other omic data types, including chromatin accessibility (ATAC-seq; e.g., Cusanovich et al., 2018; Lareau et al., 2019), joint RNA–protein profiling (CITE-seq; e.g., Peterson et al., 2017; Stoeckius et al., 2017), and multi-omic analyses (e.g., Y. Hao et al., 2021). However, as in scRNA-seq and ST, mixed-membership approaches remain largely unexplored in other omics settings.

A key challenge in both hard and mixed-membership clustering is aligning results across runs. Stochasticity (e.g., random initialization), local optima, and parameter choices can yield markedly different clustering outcomes on the same data, leading to the well-known “clustering alignment problem” encountered when analyzing multiple unsupervised clustering runs. This problem arises from (i) label switching, where cluster labels are permuted across different clustering runs; (ii) multi-modality, where runs exhibit distinct clustering patterns (i.e., “modes”) that are not reconcilable by label permutation; and (iii) difference in number of clusters *K*. Yet many scRNA-seq and ST studies run clustering only once (or a few times) and report a single solution, potentially overlooking alternative clustering solutions and consequently the informative patterns clustering reveals about cellular heterogeneity. Even when stochasticity in scRNA-seq clustering results is recognized, comparisons often lack explicit alignment of cluster labels (e.g., Fig. 3 in Sant et al., 2025; Figs. 1–2 in H. Kim et al., 2025). Likewise, in ST benchmarking, clustering results are rarely aligned despite clear spatial structure (e.g., Fig. 3 in Hu et al., 2021; Figs. 2–4 in Long et al., 2023; Figs. 2 and 6 in Yuan et al., 2024; Fig. 5 in Xu et al., 2024), so comparisons rely on summary metrics such as normalized mutual information (NMI) and cannot directly explain how models differ in grouping spots (e.g., fine-scale regions versus broader differentiation).

**Figure 1:**
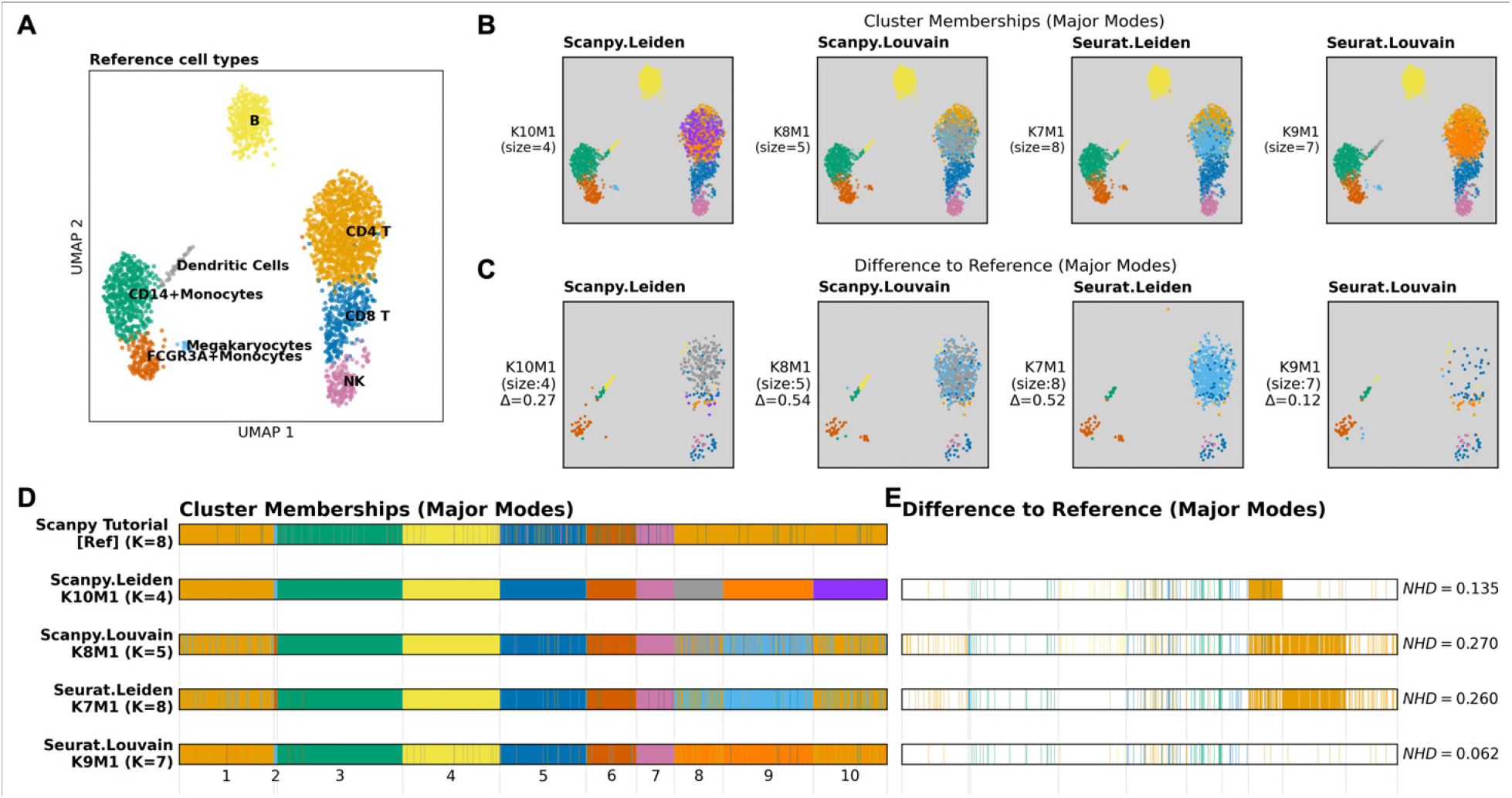
Aligned hard-clustering results for PBMC3k scRNA-seq differ substantially across models. We aligned clustering results from 10 runs of each of four models—`Seurat` Leiden, `Seurat` Louvain, `Scanpy` Leiden, and `Scanpy` Louvain—together with the reference clustering reproduced from the `Scanpy` tutorial (Leiden), using `Clumppling`. We show only the major mode (largest number of runs) for each model; all modes are provided in Supplementary Fig. S5. **(A)** Reference clustering from the `Scanpy` tutorial (Scanpy development team, 2025), shown on a 2D UMAP and colored by the eight annotated cell-type clusters. **(B)** Major modes from the four models, shown on the same UMAP as in (A). Each mode is labeled by the number of clusters *K* and mode size (in parentheses). **(C)** Cells whose labels differ from the reference in (A), colored by their mode-specific clusters. Most discrepancies occur in the following reference cell types: CD4 T, CD8 T, Dendritic Cells, FCGR3A+Monocytes, or Megakaryocytes. Each mode is annotated with the average total membership difference (Δ) which equals 2× the normalized Hamming distance (NHD) for hard clustering. **(D)** Structure plots for the reference and selected modes. Columns denote cells and stacked bars show memberships across *K* clusters, summing to 1. When annotations are available, cells are often grouped or reordered to improve visual; because PBMC3k lacks such annotations, we instead order cells by their cluster labels in the major mode at the largest *K* (“`Scanpy` Leiden K10M1”), labeled on the x-axis. **E)** Differences from the reference in structure plots, with only mismatched cells colored; NHD is reported for each mode.

**Figure 2:**
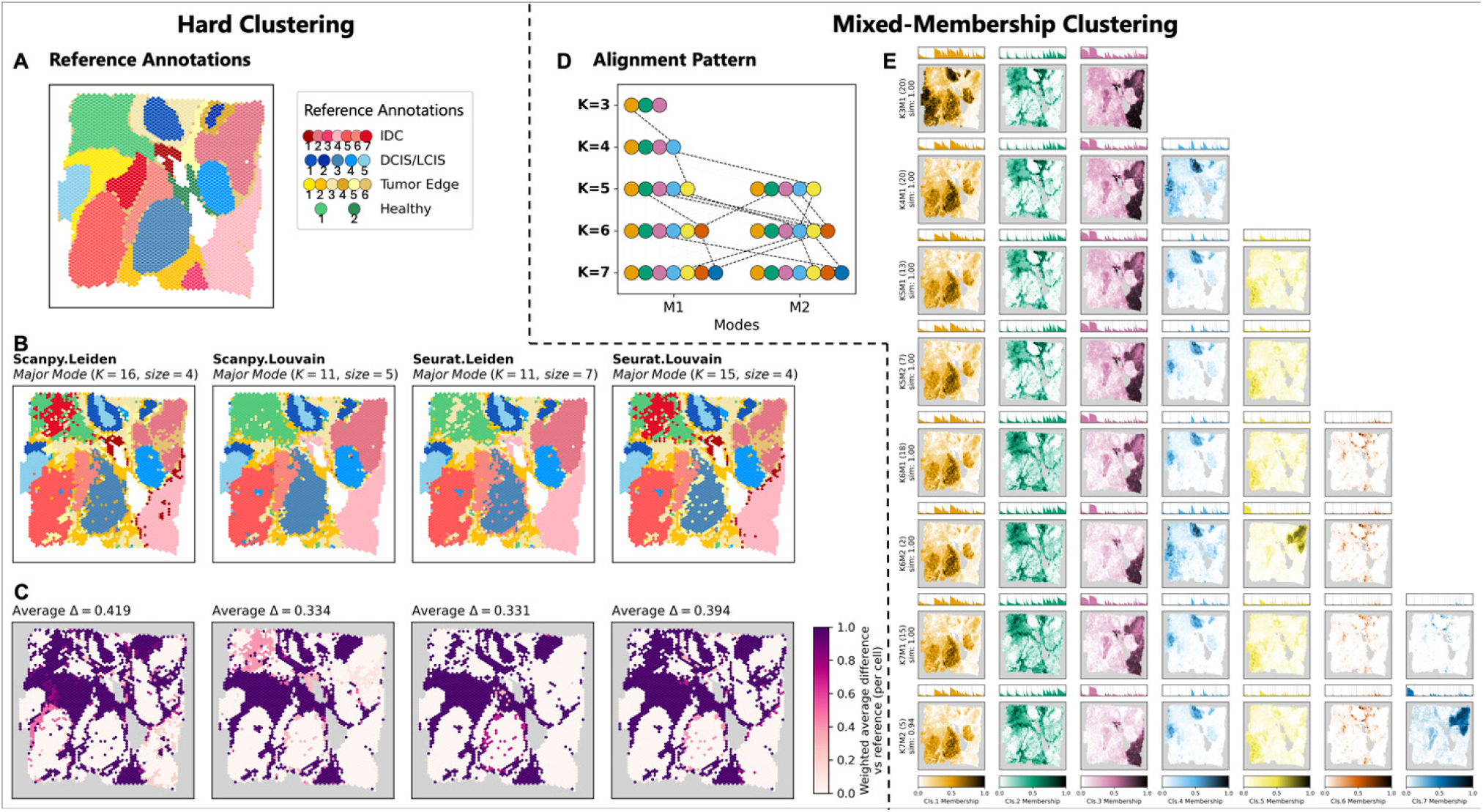
Aligned hard- and mixed-membership clustering of human breast cancer ST data shows large variability in some morphotype regions. Hard-clustering results (left) include 10 runs of each of four models—`Seurat` Leiden, `Seurat` Louvain, `Scanpy` Leiden, and `Scanpy` Louvain—together with the ground-truth annotations, all aligned using `Clumppling` (X. Liu et al., 2024). Mixed-membership results (right) include 20 runs of `FastTopics` (Carbonetto, Sarkar, et al., 2021; Carbonetto, K. Luo, et al., 2023) for each *K* from 3 to 7, aligned across modes (e.g., K3M1 denotes the first/major mode at *K* = 3). **(A)** Ground-truth segmentation into 20 regions across four morphotypes: 7 IDC, 5 DCIS/LCIS, 6 Tumor Edge, and 2 Healthy (indexed within morphotype). **(B)** Major mode from each hard-clustering model, shown in tissue coordinates and colored by cluster; each mode is labeled by *K* and the size of the mode (how many runs are in this mode). Different models produce clustering results with different *K*. **(C)** Per-spot disagreement with the ground truth, computed as the mode-size–weighted fraction of modes in which a spot’s label differs (0/1 per mode); darker colors indicate larger disagreement. For example, for a major mode of size 7 and a minor mode of size 3, if a spot differs in the major mode but not in the minor mode, this spot has a fraction value of 0.7. The average total membership difference of a mode from the ground truth is computed as the sum of absolute membership differences across all spots divided by the number of spots. The mode-size–weighted mean total membership difference (“average Δ”) is reported at the top of the plot for each model. Clustering alignment reveals the variability in clustering assignment consistency of spots in different morphotypes of the tissue. Spots with clustering labels agreeing with the ground truth occur mostly in IDC and DCIS/LCIS regions. **(D)** Cluster-alignment graph across mixed-membership modes, showing how newly emerged clusters align as *K* increases. Each node in the alignment graph denotes a cluster, and aligned clusters across modes (if differently colored) are connected. **(E)** Spatial maps of cluster memberships for each mode, shown separately by cluster. Spots in DCIS/LCIS regions tend to have large memberships in the same cluster, while tumor edge regions are less clearly isolated.

**Figure 3:**
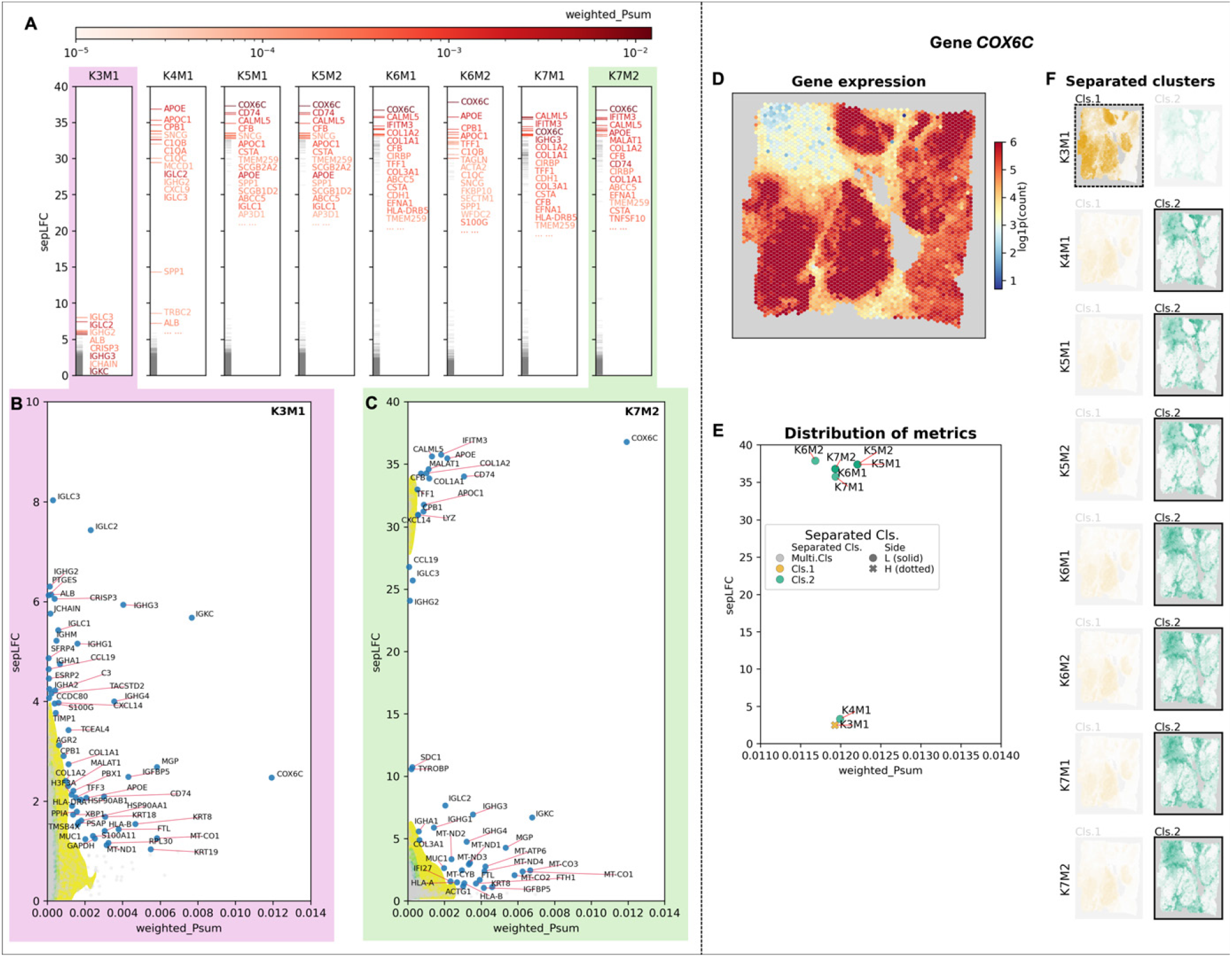
Joint feature-metric distributions from mixed-membership clustering highlight candidate clustering-informative genes in human breast cancer ST data. Feature-level metrics computed from the modes in Fig. 2 identify “outlier” genes that are clustering-informative candidates. **(A)** Top genes by largest LFC gap (*sepLF C* in Eq. 8)in each mode (modes ordered left to right). Grey ticks show the full *sepLF C* distribution; top genes are labeled in red, with color intensity proportional to total relative expression (weighted *P* sum; 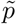 in Eq. 9). **(B–C)** Joint distributions of *sepLF C* versus weighted *P* sum for all genes in modes K3M1 (B) and K7M2 (C), shown as contour densities with selected outliers labeled. Genes such as *COX6C* and *IGKC* exhibit high values in both metrics and recur across modes. **(D)** Expression counts for *COX6C*, a candidate clustering-informative gene previously linked to breast cancer (Table 1). **(E)** *sepLF C* versus weighted *P* sum for *COX6C* across modes (one point per mode). Marker color/shape indicates the cluster(s) on the smaller side of the *sepLF C* gap. If that side contains a single cluster, the marker is colored accordingly; otherwise, the marker is shown in grey. If the fewer-cluster side has lower relative expression (L), the marker is a solid circle and the corresponding memberships are boxed with solid lines in panel F; if higher (H), the marker is a cross and memberships are boxed with dotted lines. **(F)** Membership maps for the clusters separated by *COX6C* in each mode (as in Fig. 2E), with separated clusters highlighted (dotted = higher, solid = lower relative expression). *COX6C* shows consistently high *sepLF C* for *K* ≥ 5 and repeatedly isolates cluster 2 (green), corresponding to regions where it is under-expressed (D).

Clustering alignment is important for consistent interpretation when multiple clustering solutions are generated from the same dataset. `Clumppling` (X. Liu et al., 2024), originally developed by the first author of this study for population genetic studies, uses optimization strategies to detect and align modes across clustering runs and across *K* values. Here, we extend the alignment approach of `Clumppling` to clustering outcomes in scRNA-seq and ST analyses. We introduce `ACE-OF-Clust` (Alignment, Comparison, and Evaluation of Omics Features in Clustering), a framework that extends `Clumppling` to align clustering results in single-cell transcriptomic data. We refer to a specific clustering algorithm—implemented in a given tool and applied to a particular data type—as a *clustering model*. For hard clustering, our new framework enables automated comparison of clusters across multiple runs of the same model and across different models, highlighting cells or spots with persistent inconsistencies and uncertainty. For mixed-membership clustering, we further leverage feature-level signals to identify clustering-informative features (e.g., genes in scRNA-seq and spatial transcriptomics) that drive the inferred clustering hierarchy. We also evaluate sets of features (e.g., gene sets) to assess whether they show enriched signals for cluster separation and whether features within each set contribute differently to the clustering. Taken together, `ACE-OF-Clust` provides a systematic char-acterization of clusters and the features underlying observed cellular heterogeneity. In multi-omic settings, `ACE-OF-Clust` compares clustering results within and across modalities, and proposes regulatory links (e.g., distal gene-regulatory element relationships) by identifying concordant clustering-informative features across data types (e.g., genes and accessible chromatin regions).

## 2 Results

We first present the workflow and key steps of `ACE-OF-Clust` (also see Methods and Supplementary Fig. S1). We then evaluate `ACE-OF-Clust` on three datasets: a 3k scRNA-seq dataset of Peripheral Blood Mononuclear Cells (denoted here as “PBMC3k”), an ST dataset of a human breast cancer (“HBC”) sample, and a 10k Peripheral Blood Mononuclear Cells dataset with both RNA-seq and Assay for Transposase-Accessible Chromatin using Sequencing (ATAC-seq) data (“PBMC10k”). Data descriptions and processing steps are provided in the Supplementary Materials. Specifically, we show (i) how clustering results vary across state-of-the-art models and relative to ground-truth annotations; (ii) how `ACE-OF-Clust` enables interpretable and quantitative comparisons across clustering outputs; (iii) how mixed-membership clustering profiles quantitatively assess the roles of genes in differentiating cells; and (iv) how `ACE-OF-Clust` supports multi-omic clustering comparison and examination of cross-omic links.

### 2.1 Workflow of ACE-OF-Clust

`ACE-OF-Clust` comprises four key steps: (0) generating multiple clustering results from expression data, potentially across models and parameter settings; (1) importing, organizing, and aligning clustering results using `Clumppling;` (2) interpreting and comparing aligned clustering outputs, including quantifying differences across models and relative to reference annotations; and (3) when feature-level signals are available, assessing feature contributions and identifying clustering-informative features. Step (0) is not directly implemented in our framework, as we leave users the flexibility to apply `ACE-OF-Clust` to clustering results produced by any unsupervised method for single-cell data.

`ACE-OF-Clust` takes multiple clustering results as input. Given *N* cells and *G* features (genes), a clustering method maps the data matrix *X* ∈ R^*N ×G*^ to a membership matrix *Q* ∈ R^*N ×K*^, where *K* is the number of clusters and 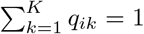 for each cell *i*. For hard clustering, *Q* is one-hot encoded. For mixed-membership clustering, we additionally use a feature-level matrix *P* ∈ R^*G×K*^ (see Methods). We refer to a single clustering result as a *clustering run*, comprising one *Q* matrix and, when available, one *P* matrix. Clustering alignment is performed by applying `Clumppling` to the set of *Q* matrices for each model. For each *K*, `Clumppling` groups runs into a small number of representative modes (distinct clustering solutions), with the one containing the most runs being the “major mode”, then aligns them across adjacent *K* values, yielding an alignment quality measure for each aligned mode pair. Columns of each *Q* matrix, which encode the memberships of all cells to a given cluster, are permuted to conform to the optimal alignment pattern. Across *K*, multiple clusters at higher *K* may align to a single cluster at lower *K*, capturing cluster splitting and merging patterns as *K* varies. For multi-model comparison, we additionally align modes across models via a two-level alignment procedure.

To quantify differences in aligned modes, we compute the normalized Hamming distance (NHD), defined as the ratio of cells with differing labels to the total cell count, between aligned cluster labels for hard clustering. This can also be interpreted as the fraction of different cell-cluster assignments between two clustering results. More generally, `ACE-OF-Clust` measures the difference between two modes with membership matrices *Q* and *Q*^*′*^ using the average total membership difference 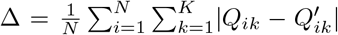. When the modes have different *K* values, we first merge clusters in the higher-*K* mode according to the alignment so that *Q* and *Q*^*′*^ have matching dimensions. For hard clustering, the per-cell sum equals 0 or 2, so Δ = 2× NHD. If annotation labels are available, we additionally compute a per-group difference for cells within each annotated group.

When *P* is available from mixed-membership clustering, we construct a *clustering profile* for each feature *j* comprising two vectors: a log fold change (LFC) vector *L*^*j*^, and an index vector 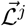. After sorting the corresponding *P* entries in increasing order as 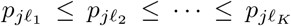, where the index vector 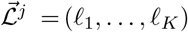 is a permutation of {1, …, *K*}, we compute 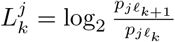 (see Supplementary Fig. S3 for an example). By construction, 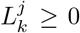, and larger values indicate a greater fold difference between adjacent relative feature levels. The clustering profile thus provides a direct measure of cluster separation on a (log) expression scale, offering an interpretable indicator of how informative a feature is for distinguishing clusters. We then derive metrics to quantify the clustering informativeness of each feature. Specifically, we focus on two metrics: the total relative expression 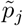 (the “weighted *P* sum”), defined as 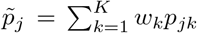 with 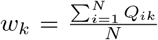, which summarizes a feature’s overall contribution to clustering; and the largest separation gap (“*sepLF C*”), defined as 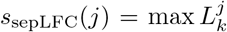, which quantifies a feature’s strongest separating signal between two sets of clusters (see Methods). If a feature is broadly informative for clustering (high weighted *P* sum) and substantially contributes to cluster separation (high *sepLFC*), then we consider it a clustering-informative feature. Downstream analyses can then be performed to investigate or validate the functional roles of these features, or to evaluate known feature sets with shared biological programs and assess their joint effects or distinct behaviors (see Supplementary Materials S1.12).

### 2.2 Revealing variability across clustering models on benchmark PBMC3k data

The PBMC3k scRNA-seq dataset (Universal 3’ Gene Expression Dataset by Cell Ranger v1.1.0, 2016) is a popular benchmark and is frequently used as an illustrative example for single-cell computational tools, most notably in the tutorials for `Seurat` (Satija, 2023) and `Scanpy` (Scanpy development team, 2025). We selected this dataset to demonstrate that `ACE-OF-Clust` enables quantitative model comparison in single-cell clustering workflows, which is often overlooked or performed only qualitatively.

We examined four hard clustering models—`Scanpy` (Wolf et al., 2018) and `Seurat` (Satija et al., 2015; Butler et al., 2018), each paired with the Louvain (Blondel et al., 2008) and Leiden (Traag et al., 2019) community detection algorithms. Although we focus on these widely used models, `ACE-OF-Clust` is compatible with any clustering model. To enable a direct comparison, we used the same clustering settings across the two software packages (see Supplementary Materials). Even under identical algorithmic settings, intrinsic differences and algorithmic randomization still produced substantial variation in clustering results between `Seurat` (Satija et al., 2015; Butler et al., 2018) and `Scanpy` (Wolf et al., 2018), as well as among the runs produced by the same software with different random seeds (Fig. 1). We used the cell-type annotations from the `Scanpy` tutorial as reference labels. In the tutorial, cells are partitioned into eight clusters, marker genes are ranked for each cluster, and cell-type labels are assigned based on these markers (Scanpy development team, 2025). We show the reference labels in Fig. 1A on a 2D UMAP generated following the tutorial (Scanpy development team, 2025).

We visualize clustering alignment using *structure plots* (Fig. 1D), i.e., stacked bar charts commonly used for mixed-membership results in population genetics (Rosenberg et al., 2002; Rosenberg, 2004), and by projecting cells onto the UMAP embedding (Fig. 1B). We show only the major modes among the 10 runs per model; all modes are provided in Supplementary Fig. S5. Even under identical algorithmic settings, the inferred *K* values varies widely across runs, and aligned solutions differ substantially (Figs. 1C,E). Thus, running clustering only once, which is common practice, risks producing a minor mode not representative of the cell grouping patterns identified across clustering runs. The *normalized Hamming distance* (NHD) between each model’s clustering and the reference cell-type labels ranges from 0.062 to 0.270 across the selected modes, further quantifying the degree of disagreement and indicating that a substantial fraction of cells are assigned to different clusters (Fig. 1D). The cell group most prone to disagreement across models is the CD4 T cells, accounting for the majority of mismatches relative to the reference; in contrast, the CD14+Monocytes are clustered most consistently across all four models.

We also aligned mixed-membership results from `FastTopics` (Carbonetto, K. Luo, et al., 2023) on PBMC3k. We find that cells predominantly assigned to particular cell types progressively subdivide as the number of clusters increases (e.g., separating CD8 T from NK cells as *K* increases from 2 to 3), and that some clustering-informative genes contribute weakly at smaller *K* but become substantially more informative at higher *K* (see Supplementary Materials S1.8). Our pipeline reveals non-negligible variability both within and across clustering models on this benchmark dataset, underscoring the value of performing multiple runs followed by clustering alignment.

### 2.3 Examining intratumoral heterogeneity in breast cancer

We applied `ACE-OF-Clust` to 10X Genomics Visium ST data from a human breast cancer (HBC) sample (Spatial Gene Expression Dataset by Space Ranger v1.1.0, 2020), previously studied by Xu et al. (2024), as this dataset includes human-annotated morphotypes that serve as spot labels. Mixed-membership clustering was performed on all 24,923 genes with variation in read count, whereas hard clustering is performed on a subset of 3,000 highly variable genes. Manual pathology labeling based on H&E staining is provided with the data, and we used these labels as reference annotations. There are four main labeled morphotypes across 20 regions: five ductal carcinoma in situ/lobular carcinoma in situ regions (“DCIS/LCIS”), two healthy regions (“Healthy”), seven invasive ductal carcinoma regions (“IDC”), and six tumor surrounding regions with low features of malignancy (“Tumor Edge”). We show them in different colors in Fig. 2A, e.g., red for IDC, with shades of a given color denoting different regions in the same morphotype. Xu et al. (2024) applied multiple clustering approaches to this dataset (see their Fig. 5) using *K* = 20, but did not align or quantitatively compare the resulting clusters either against one another or against the annotations.

#### 2.3.1 Consistency of clustering results varies across pathological annotations

We aligned modes from 10 runs of the same four hard-clustering models applied to PBMC3k: `Seurat` Leiden, `Seurat` Louvain, `Scanpy` Leiden, and `Scanpy` Louvain (Blondel et al., 2008; Satija et al., 2015; Wolf et al., 2018; Traag et al., 2019). We then aligned modes across all models, together with the *K* = 20 reference annotations. Fig. 2B shows the major modes identified for each model; all modes are provided in Supplementary Figs. S10 and S11. As in our PBMC3k hard-clustering analysis, HBC results vary both across models and across repeated runs, and the inferred number of clusters *K* sometimes varies substantially across random initializations. For example, across 10 runs, `Seurat` Louvain yields three modes: *K* = 13 (3 runs), *K* = 14 (3 runs), and *K* = 15 (4 runs, the major mode shown in Fig. 2B). The aligned clustering results broadly agree with the annotated morphotypes, although moderate discrepancies remain. Such variability is expected when using simple nonspatial hard clustering; here, our goal is to showcase how `ACE-OF-Clust` can quantify and evaluate these discrepancies.

We computed the normalized Hamming distance (NHD) between each mode and the ground truth and report mean NHD across modes for each clustering model (Fig. 2C). For each spot, we also computed a disagreement fraction as the fraction of modes whose labels disagree with the ground truth and visualized these values as a heatmap, weighted by their mode sizes. For example, under `Scanpy` Leiden, a spot mislabeled in all modes has value 1, whereas a spot mislabeled only in the major mode (4 of 10 runs) has value 0.4. Using both NHD and the per-spot disagreement fractions, we find that all hard-clustering models correctly distinguish most spots in the large IDC and DCIS/LCIS regions (IDC 2, 4, and 5; DCIS/LCIS 3, 4, and 5). In contrast, tumor-edge regions and smaller morphotype regions remain difficult to resolve, as reflected by the dark-colored spots in Fig. 2C. Among major modes, `Seurat` Louvain and `Scanpy` Leiden perform worse in the large healthy region (Healthy 1) while `Scanpy` Louvain performs the best. When aggregating across all modes, however, `Seurat` Leiden performs best overall at separating this healthy region from adjacent DCIS/LCIS and tumor-edge regions (Fig. 2C).

Beyond discrete separation of regions, many spots exhibit continuous variation that is captured by mixed-membership clustering. We summarize the aligned mixed-membership clustering results in Figs. 2D,E (see also Supplementary Materials S1.9), where the emergence and splitting of clusters as *K* increases reveals spatial transcriptomic structure across pathological morphotypes. Spots in IDC 5 show moderate membership in Cluster 1 (orange), which dominates DCIS/LCIS 3, but they also exhibit appreciable membership in Cluster 4 (blue), the dominant cluster in DCIS/LCIS 1 and 5, and in Cluster 5 (yellow), a noisier cluster with memberships dispersed across many tissue regions. Together, these patterns suggest that IDC 5 spots are resolved as intermediate between multiple DCIS/LCIS regions, showing a spatial mixing or gradual transitions in cell-state composition. These non-discrete heterogeneity patterns are difficult to resolve with hard clustering, underscoring mixed-membership clustering as a valuable foundation for future single-cell and spatial clustering analyses.

#### 2.3.2 Clustering profiles identify candidate genes that are informative for clustering

In single-cell analyses, many pipelines, including `Seurat` (Satija et al., 2015; Butler et al., 2018) and `Scanpy` (Wolf et al., 2018), identify “marker genes”—genes most characteristic of each cluster—via differential expression (DE) and use them to label clusters. However, each gene’s contribution to the clustering itself and to cluster separation has rarely been studied. To identify genes driving mixed-membership clustering results, we computed each gene’s total relative expression (weighted *P* sum; Eq. 9) and largest separation gap (*sepLFC;* Eq. 8) across modes. Several genes with large *sepLFC* and non-negligible weighted *P* sums recur across modes (Fig. 3A), including *COX6C, APOE*, and *COL1A1*, which have been associated to breast cancer in prior studies (Table 1). We further visualized the joint distribution of these metrics for each mode using contour plots with overlaid scatter points (Fig. 3B,C) to highlight outlier genes, which we treat as candidate clustering-informative genes for downstream investigation.

**Table 1.** We list several clustering-informative genes from mixed-membership clustering of the HBC dataset (Fig. 3A–C); they have been previously associated with breast cancer.

|  | Gene | Figure of clustering profile (if included) | Findings from previous publications |
| --- | --- | --- | --- |
| 1 | Cytochrome c oxidase subunit 6C (*COX6C*) | Fig. 3D-F | Yin et al., 2020 identifies *COX6C* as one of the hub genes with significantly elevated mRNA and protein expression correlated with breast cancer development. See also West et al., 2001; Emerson et al., 2009. |
| 2 | Apolipoprotein E (*APOE*) | Supplementary Fig. S13 | Harrison et al., 2021 shows that genetic polymorphisms in gene *APOE* is associated with neuroconnectivity in breast cancer patients. See also Y. Zhou et al., 2020; Darwish et al., 2023. |
| 3 | Immunoglobulin kappa constant (*IGKC*) | Supplementary Fig. S13 | Schmidt et al., 2009; Marcus Schmidt, Hellwig, et al., 2012; Marcus Schmidt, Edlund, et al., 2021 find prognostic significance of *IGKC* in node-negative breast cancer patients. |
| 4 | Collagen, type I, alpha 1 (*COL1A1*) | N/A | Wang et al., 2018 identifies *COL1A1* as a cancer stromal key gene associated with breast cancer invasion and metastasis. See also Jing Liu et al., 2018; Xue Li et al., 2022. |
| 5 | Matrix Gla protein (*MGP*) | N/A | T. Du et al., 2022 and S. H. Yang et al., 2025 both find *MGP* to be an immunohistochemical marker that supports breast origin, upregulated in breast carcinoma. See also Caiado et al., 2023; Y. Wu et al., 2024. |

Examining these metric values for a gene across modes clarifies the role of the gene in the inferred clustering results. To illustrate this, we show results for *COX6C*, an outlier in Figs. 3B-C, in Figs. 3D-F. Two additional genes (*APOE* and *IGKC*) are shown in Supplementary Fig. S13. *COX6C* contributes strongly to the emergence of cluster 2 (green), which has high membership in several healthy and tumor-edge regions. It has large *sepLFC* at large *K* (*K >* 4), indicating that it is most informative for differentiating finer-scale clusters. More generally, `ACE-OF-Clust` enables gene-level investigation of clustering results, revealing that even genes not highlighted as traditional markers can be informative for how clusters are separated.

Notably, restricting to 3,000 highly variable genes (HVGs) selected by `Scanpy` excludes some top-ranked clustering-informative genes, including *COX6C*, despite their high expression in certain regions corresponding to the identified clusters. We further assessed how selecting a subset of genes—typically highly variable genes (HVGs)—affects mixed-membership clustering outcomes. We compared the results obtained using 3,000 HVGs selected by the standard `Scanpy` procedure against gene subsets defined by our clustering-informative metrics. HVGs and non-HVGs showed highly similar distributions of weighted *P* sum and *sepLFC* (Supplementary Fig. S14), indicating that restricting to HVGs can discard non-HVG genes that are equally informative for clustering. In contrast, filtering out genes with low weighted *P* sum or *sepLFC* preserved the dominant clustering patterns, whereas imposing more stringent *sepLFC* thresholds altered results. This suggests that clustering is largely insensitive to non-informative genes but is influenced by genes with non-negligible separation signal. See Supplementary Materials S1.10 for details. Complementary to clustering alignment, `ACE-OF-Clust` offers a post hoc framework for evaluating—and potentially guiding—feature selection in clustering.

#### 2.3.3 Gene set analysis reveals cluster enrichment and heterogeneous gene contributions within the gene set

In addition to measuring the influence of individual genes on clustering, it is often informative to examine groups of genes together, particularly those thought to participate in the same biological pathways, processes, or cellular states. To demonstrate how `ACE-OF-Clust` supports gene set analysis, we investigated hallmark gene sets in the HBC clustering results. Hallmark gene sets are curated collections of genes that summarize well-defined biological processes (Liberzon et al., 2015). The results for gene set *Hallmark E2F Targets* (a set of cell cycle-related targets of E2F transcription factors) are shown in Supplementary Figs. S19-S23; the methods are described in Supplementary Materials S1.12. Briefly, for each clustering mode, we used the gene-set average relative expression across clusters to test whether the gene set showed enriched cluster-specific expression or an unusually large *sepLFC* relative to random gene sets. We then decomposed the gene-set-level separating bipartition to identify individual genes and cluster pairs contributing most strongly to the observed separation. These analyses can be applied to any gene set of interest, including known marker gene sets or subsets of clustering-informative candidate genes identified in earlier steps.

A gene set can be enriched either in specific clusters or in the separation between specific sets of clusters across clustering results with different numbers of clusters. For example, the *Hallmark E2F Targets* set shows elevated mean relative expression in Cluster 3 (pink; Fig. 2E), which largely corresponds to IDC regions, across all clustering modes. This pattern suggests that these IDC spots are enriched for actively cycling, highly proliferative tumor cells. In clustering modes with fewer clusters, the *sepLFC* of this gene set is significantly different from the null distribution of random gene sets of the same size (*p <* 0.05), separating Cluster 2 (green; Fig. 2E; healthy and tumor-edge spots) from the remaining clusters. This enrichment is not observed in finer clustering results with more clusters, where the gene set instead separates Cluster 6 (dark orange; Fig. 2E; a subset of tumor-edge spots split from the earlier-emerging Cluster 2) from the rest (Supplementary Figs. S19B-C, S20C-D, and S22C-D). Individual genes within the same set contribute heterogeneously to these clustering patterns.

Genes with the highest relative expression in a given cluster are not necessarily the same as those with *sepLFC* across the clusters separated by the gene set as a whole; in many cases we find that they are not (Supplementary Figs. S20E-F and S22E-F). By examining the top genes contributing to the cluster separation, we find that *BRCA1* has the largest gene-specific *sepLFC* between Clusters 2 and 6 in mode *K*7*M* 2 (Supplementary Fig. S22F), followed by *SPC25*. Notably, the gene set has its lowest mean relative expression in Cluster 6 and its second-lowest mean relative expression in Cluster 2 (Supplementary Fig. S22A). This suggests that the separation between these two tumor-edge-associated clusters may reflect context-dependent variation within tumor-edge regions rather than a uniformly elevated E2F program. One possible explanation is that tumor-edge spots are heterogeneous: some, such as those in Cluster 2, may retain stronger epithelial or proliferative signals, whereas others, such as those in Cluster 6, may be more stromal and contain fewer epithelial or cycling cells and more fibroblasts or immune cells. Genes outside the focal gene set with comparably high *sepLFC* values between the separated clusters are also shown alongside the top contributing genes within the set (Supplementary Figs. S21C-D and S23C-D). Together, these results suggest that the separation between these clusters is not driven solely by the biological program defined by the hallmark gene set. Instead, the additional genes may reflect parallel or overlapping biological programs that distinguish the same spots, or indicate that the cluster differentiation involves multiple biological axes.

Together, enriched relative expression, enrichment of the largest separation gap, and the corresponding top contributing genes provide complementary perspectives for assessing how a gene set contributes to clustering and for linking biological programs to cluster differentiation. These results demonstrate that `ACE-OF-Clust` gene set analysis can reveal both cluster-level enrichment of coherent biological programs and substantial heterogeneity in how individual genes within the same set contribute to the observed clustering patterns.

### 2.4 Comparing clustering results within and across PBMC10k multi-omic data

`ACE-OF-Clust` generalizes beyond standalone scRNA-seq and spatial transcriptomics (ST) analyses: it can integrate and compare multi-omic clustering results whenever the underlying entities (cells) are shared across modalities. Using the PBMC10k single-cell multiome dataset (Epi Multiome ATAC + Gene Expression dataset analyzed using Cell Ranger ARC 1.0.0, 2020), previously used to showcase multi-omic tools such as `muon` (Bredikhin et al., 2022), we demonstrate how to 1) align clustering results within and across omics, 2) quantify membership variation within annotated cell groups, and 3) examine relationships among clustering-informative features across modalities. `ACE-OF-Clust` reveals multi-omic differences that reflect sources of variability not apparent in single-omic analyses (e.g., PBMC3k), informing downstream tasks such as gene regulatory network inference and providing insight into cross-omic regulatory relationships.

We generated 2D UMAP embeddings following the `muon` tutorial (Bredikhin, 2025) and show the corresponding reference annotations in Supplementary Fig. S16C. We performed hard clustering on both scRNA-seq and scATAC-seq using the same four models, with 50 runs each, and compared aligned results across all eight model-omic combinations. For downstream analyses by cell type (e.g., gene regulatory network inference (Aibar et al., 2017)), it is important to identify cell groups that are consistently clustered; if an annotated group is not stable, using a single clustering solution as the reference is misleading. `ACE-OF-Clust` is designed to quantify and highlight such variability that can affect downstream analyses. As expected, clustering variability was larger across omic data types than within. Stratifying differences by annotated groups (Supplementary Fig. S16D) further shows that variability is cell-type dependent. See Supplementary Materials S1.11 and Supplementary Fig. S16 for more details.

#### 2.4.1 Alignment of ATAC-seq and RNA-seq clustering modes validates regulatory relationships between peaks and genes

Aligning and comparing cross-omic clustering results enables broader analyses, as we illustrate with the best-aligned RNA-seq/ATAC-seq mode pair (Supplementary Fig. S17). For this pair, we quantified within–cell-type membership variability (using the annotations in Suppelementary Fig. S16C) for both omics data types via the bootstrapped FStruct ratio (Morrison et al., 2022), where larger values in Fig. 4A indicate higher variability in cluster memberships. ATAC-seq shows substantially higher FStruct ratio values than RNA-seq in cell types CD4+ memory T, CD4+ naive T, and CD8+ naive T, suggesting these subtypes are resolved differently by RNA- and ATAC-based clustering, with ATAC-seq exhibiting greater variability.

**Figure 4:**
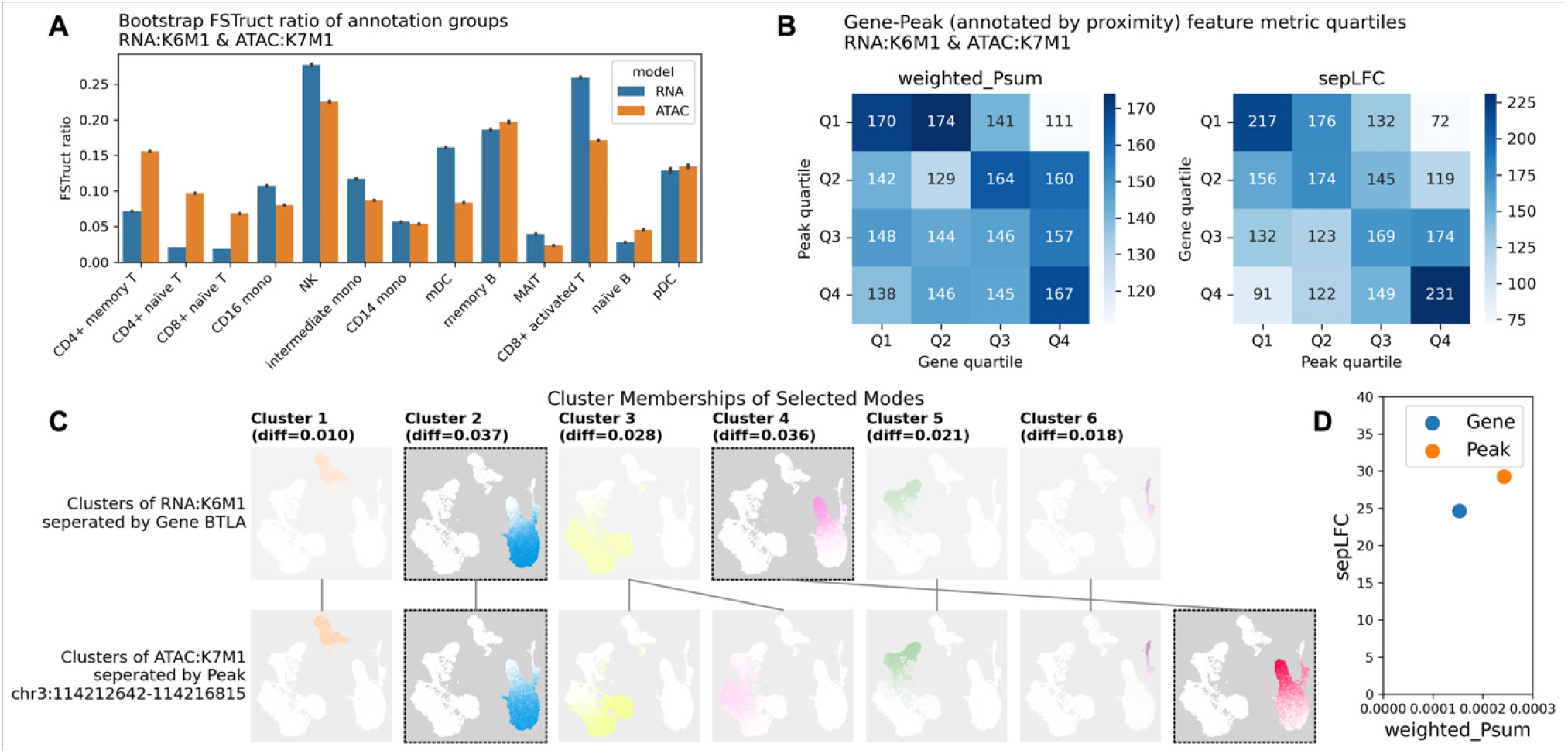
Comparing scRNA-seq and scATAC-seq mixed-membership clustering reveals celltype–dependent membership variation and highlights putative gene–peak links. We focus on RNA-seq mode K6M1 and ATAC-seq mode K7M1, the aligned pair with the smallest average total membership difference (Supplementary Fig. S17A). **(A)** Membership variation within each annotated cell group (Fig. S16C), quantified by the FStruct ratio (Morrison et al., 2022). **(B)** Heatmaps of proximal gene–peak pairs binned by joint quartiles of feature metrics (RNA-seq on x-axis; ATAC-seq on y-axis): total relative gene expression (weighted *P*-sum; Eq. 9) and largest separation gap (*sepLFC*; Eq. 8). **(C)** Memberships for the two modes, with clusters separated by the example features—*BTLA* and *chr3:114212642-114216815* — highlighted (others dimmed). Aligned clusters are connected by lines, and mean per-cluster differences are shown at the top; undimmed memberships are in Supplementary Fig. S17B. **(D)** *sepLFC* versus weighted *P*-sum for the selected gene and peak; both are high, consistent with clustering-informative candidates.

In PBMC10k, each ATAC peak is annotated with one or more nearby genes. For the selected cross-omic mode pair, we compared feature-level metrics (weighted *P* sum and *sepLFC*) by tabulating proximal gene–peak pairs across joint quartiles in the two modalities (Fig. 4B). Although many pairs fall on the diagonal (both high or both low), a substantial fraction lie in off-diagonal quartiles (∼ 74.3% for weighted *P* sum and ∼66.8% for *sepLFC*), indicating that proximity does not necessarily imply similar clustering informativeness. This motivates examining non-proximal gene–peak links as candidate regulatory relationships, particularly among pairs with high metrics in both modalities.

By looking at features with hight *sepLFC* and weighted *P* sum, the same procedure as for PBMC3k and HBC, we identified candidate clustering-informative genes and peaks in PBMC10k. We then focused on gene–peak pairs with large *sepLFC* that separate the same aligned clusters in the selected RNA/ATAC mode pair. In Figs. 4C-D, we highlight one such pair—*BTLA* and *chr3:114212642-114216815* —and show the clusters they separate and their feature-level metrics; both exhibit high values for feature metrics and are thus clustering-informative candidates. This non-proximal pair is reported in the Registry of human candidate cis-Regulatory Elements (cCREs) (Moore et al., 2020), providing independent evidence consistent with a regulatory link. We investigate gene–peak pairs (3,026 genes; 14,896 peaks) that separate matched clusters in this selected mode pair. Because RNA- and ATAC-based clusterings can still differ appreciably after alignment, and because separated clusters can be sensitive to the estimated *P* matrices, peaks and nearby genes may not share the same separation patterns. Among 1,017 annotated gene-peak pairs supported by either cCREs or proximity (maximum distance 199,548 bp), about half are supported by cCREs but not proximity. Among the 222 pairs with *sepLF C >* 7.5 for both the gene and the peak, about onethird are supported only by cCREs (Supplementary Fig. S18). These results suggest putative regulatory relationships between genes and peaks that contribute similarly to cluster separation despite a lack of genomic proximity. Overall, our clustering profiles provide a new approach to prioritize cross-omic links based on shared contributions to differentiating and grouping cells.

## 3 Discussion

Clustering is central for identifying cell types in scRNA-seq and ST studies. Many analyses follow a standard pipeline (e.g., the `Scanpy` tutorial (Wolf et al., 2018; Scanpy development team, 2025)) that performs a single run of hard clustering and assigns cell types based on the resulting clusters; however, this approach has caveats that can compromise robustness. To refine and extend this pipeline, we develop `ACE-OF-Clust`, a clustering alignment and analysis framework for single-cell clustering based on `Clumppling` (X. Liu et al., 2024). Our framework summarizes repeated runs of clustering models across varying *K* into representative modes, then aligns modes across models—and, when available, to reference annotations—for direct quantitative comparison and tracking of cluster emergence. When mixed-membership feature-level outputs are available, `ACE-OF-Clust` constructs clustering profiles and derives metrics to prioritize clustering-informative features. We provide four practical recommendations to improve the reliability and interpretability of singlecell clustering and to guide downstream computational and molecular follow-up.

First, we recommend running clustering multiple times, even when *K* is fixed. In all our data demonstrations, widely used tools produced markedly different clustering solutions that are difficult to justify or interpret without repeated runs and alignment (Supplementary Figs. S5,S6,S10,S11). Therefore, we view aligning multiple clustering solutions in single-cell clustering analyses as both an opportunity and a necessity.

Second, examining multiple models jointly—especially for hard clustering—helps assess the consistency of cluster assignments and identify cells with uncertain labels (Figs. 1,2). Alignment provides both a global view of clustering patterns and cell-level label comparisons across models, and it naturally extends to different settings of the same algorithm. Although we do not aim to optimize hard clustering, `ACE-OF-Clust` streamlines alignment of divergent runs and enables direct comparisons that can reflect cell-specific or algorithmic-based sources of clustering variability.

Third, for mixed-membership clustering we recommend using all genes, rather than restricting to highly variable genes (HVGs), whenever feasible. In HBC, many non-HVG (“less variable”) genes still contribute meaningfully to the clustering, indicating that HVG filtering can omit clustering-informative signal (Supplementary Fig. S14). Although restricting to HVGs can reduce computational costs, it risks discarding genes that drive the inferred clusters.

Last but not least, we recommend using `ACE-OF-Clust` to integrate and compare multi-omic clustering results. By quantifying within- and cross-omic disagreement, it highlights cell groups that are consistently clustered versus those whose memberships shift across data types (Supplementary Fig. S16). In addition, concordance of clustering-informative features can prioritize putative cross-omic links beyond genomic proximity for follow-up with independent regulatory evidence (Fig. 4).

`ACE-OF-Clust` is designed to align *unsupervised* clustering results. If the clustering procedure includes supervised components—for example, if some groups are pre-defined (e.g., morphotype labels are provided to the algorithm before clustering)—we do not recommend using it, as it ignores available label information and treats clusters as unknown. Although the framework can still be applied and evaluated against the supervised labels, it is likely less efficient.

Another limitation of `ACE-OF-Clust` is that alignment assumes the same set of cells across clustering results, making it unsuitable for pseudotime analyses that integrate multiple static snapshots to reconstruct temporal dynamics (Ding et al., 2022). Because snapshots contain distinct cells, clustering results cannot be aligned across time points. However, `ACE-OF-Clust` remains applicable to the intermediate clustering step within a snapshot used to initialize trajectories; repeating and aligning these clustering results can help derive a more reliable consensus structure for downstream pseudotime inference.

Additionally, assessing clustering-informative genes in mixed-membership clustering depends on the quality and stability of the feature-level relative expression estimates (the *P* matrices). Although `ACE-OF-Clust` aligns *P* matrices within each mode via the aligned *Q* matrices, per-cluster gene weights can still vary slightly across runs. Because these relative expression values are small (summing to one across genes within each cluster), even minor fluctuations can produce noticeable changes in *sepLFC* value, potentially affecting interpretation of a gene’s contribution to cluster separation.

As a general tool for aligning unsupervised clustering results, `ACE-OF-Clust` extends beyond functional genomics. It supports any application that requires comparing, summarizing, and interpreting multiple clustering solutions, including feature-level assessment of the inferred structure. Continued methodological extensions and new domains of application will further expand its utility and enable additional discoveries. Future work may augment our framework via multiple new directions. Alignment algorithms could incorporate cell-specific weights to emphasize higher-quality profiles or biologically prioritized regions. Additional feature-level metrics and alternative representations of clustering profiles may offer new insights into regulatory structure and transcriptomic variation. Extending alignment to partially overlapping (or similar) cell sets would broaden applicability, including to pseudotime settings. Finally, integrating clustering-derived signals with other molecular layers (e.g., genetic variation, chromatin accessibility, and epigenomic marks) could help prioritize regulatory relationships and generate mechanistic hypotheses about cell-type–specific function and gene-expression dynamics.

## 4 Methods

We detail each step of the `ACE-OF-Clust` workflow, summarized in Supplementary Fig. S1, in this section, including generation of clustering results. Our framework is implemented as an open-source Python package `ace-of-clust` (see Code Availability).

### 4.1 Clustering of single-cell gene expression data

Single-cell RNA sequencing (scRNA-seq) offers high-resolution information about gene expression at the individual cell level. Spatial transcriptomics (ST) data with single-cell resolution combines the gene expression profiles of individual spots with their spatial location within a tissue; ST technologies are rapidly approaching single-cell resolution, with spots in many present-day datasets each representing 1-10 cells. To unify the notations in our methodological framework, we refer to both “barcodes” in sequencing protocols and “spots” in spatial transcriptomics as “cells,” regardless of whether they represent actual single cells.

In the clustering context, cells are the individual data points, and genes are the features. The information input to clustering algorithms for *N* cells and *G* genes are recorded in a gene expression count matrix *X* of size *N* × *G*. Note that `ACE-OF-Clust` does not explicitly incorporate spatial information, unlike spatial clustering methods (Hu et al., 2021; E. Zhao et al., 2021; Long et al., 2023; Yuan et al., 2024). We note that many spatial clustering methods couple a spatially encoded representation of gene expression with classic nonspatial clustering (Hu et al., 2021; E. Zhao et al., 2021; Long et al., 2023; Yuan et al., 2024), meaning that our alignment framework for unsupervised clustering is still applicable to spatially aware clustering outputs. Our methodological framework is thus broadly applicable to any single-cell clustering analysis, while still allowing spatially aware methods—provided that they produce outputs in a compatible format, as detailed below—to fit seamlessly within it.

Clustering is applied to the input matrix *X*. Suppose there are *K* clusters. In graph-based hard clustering (e.g., Louvain, see Blondel et al. (2008); and Leiden, see Traag et al. (2019)), *K* is inferred by the clustering algorithm. In some other clustering models, like k-means clustering or topic-modeling-based mixed-membership clustering (Dey et al., 2017; Carbonetto, Sarkar, et al., 2021), *K* is specified by the user as an input. Results from a single clustering run are encoded in a membership matrix *Q* of size *N × K*, where each entry *q*_*ik*_ is the inferred membership coefficient of individual *i* in cluster *k*, with 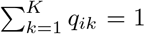. In hard clustering, *q*_*ik*_ ∈ {0, 1}; in mixed-membership clustering, 0 ≤ *q*_*ik*_ ≤ 1. See Supplementary Fig. S2 for a schematic overview of the input and output data matrices.

Additionally, many mixed-membership clustering algorithms, especially those derived from topic-modeling approaches (Dey et al., 2017; Carbonetto, Sarkar, et al., 2021), also output a feature-level matrix, *P*, reflecting gene signals across clusters. *P* is of size *G × K*, with 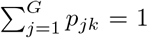 for all *k* (Supplementary Fig. S2). Information contained in *P* matrices have been used to identify distinctive genes for the detected clusters (Dey et al., 2017; Carbonetto, K. Luo, et al., 2023). We use the *Q* matrices from multiple clustering solutions as inputs to clustering alignment, and use the *P* matrices—combined with the *Q* matrices—to construct gene profiles that quantify their contributions to clustering.

For hard clustering analyses, we benchmark the two popular tools, `Scanpy` (Wolf et al., 2018) (implemented in Python) and `Seurat` (Satija et al., 2015; Butler et al., 2018) (implemented in R); we apply two community detection clustering algorithms implemented in these tools, the Louvain algorithm (Blondel et al., 2008) and the Leiden algorithm (Traag et al., 2019). For hard clustering, we follow standard practice by selecting a subset of highly variable genes as the input—those most likely to drive cell-to-cell variation within a homogeneous population (Yip et al., 2019), since community detection algorithms become highly inefficient when applied to all genes. Meanwhile, for mixed-membership clustering, we keep all genes (other than those with no variation) in the analysis. We use `FastTopics` (Carbonetto, K. Luo, et al., 2023) to perform mixed-membership clustering on transcriptomic data, which uses Poisson non-negative matrix factorization for topic-modeling fitting. `FastTopics` remains efficient and is robust to the existence of low-variance genes (Carbonetto, K. Luo, et al., 2023).

### 4.2 Clustering alignment and model comparison

As stated before, let *K* be the number of clusters in a clustering output run. For each *K* value, suppose there are a total of *R*_*K*_ runs of the clustering solutions, denoted as 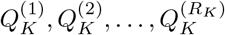. If the number of clusters is pre-specified by the user, *R*_*K*_ may be the same for all *K*; if the number of clusters is inferred from the model, they may vary for different *K*. `ACE-OF-Clust` performs clustering alignment using `Clumppling` (X. Liu et al., 2024), which was previously developed for population-genetic applications. `Clumppling` first aligns clustering runs with same *K* values and then finds *m*_*K*_ modes among the *R*_*K*_ runs of *K*. The consensus memberships of each mode can either be the average of all runs in the mode or the memberships of a representative run. Here we choose to use the latter for interpretability in empirical analyses, particularly for feature-level signals we study later in this work.

We denote the alignment between a pair of runs (or modes) in terms of their membership matrices as follows. Given two membership matrices 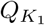 and 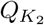 with *K*_1_ ≤ *K*_2_, the alignment pattern is represented by a mapping *α* such that *α*(*j*) = *i* for *i* ∈ [*K*_1_] and *j* ∈ [*K*_2_]. Each cluster *j* maps to exactly one cluster *i*, and each *i* is the image of at least one *j*. If *K*_1_ = *K*_2_ = *K*, then *α* is a permutation of [*K*] = {1, 2, …, *K*}; if *K*_1_ *< K*_2_, then *α* is a many-to-one mapping from [*K*_2_] to [*K*_1_].

For example, suppose *K*_2_ = *K*_1_ + 1 and *α* maps both *j*_1_, *j*_2_ ∈ [*K*_2_] of 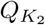 to some *i*^*^ ∈ [*K*_1_] of 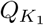, that is, *α*(*j*_1_) = *α*(*j*_2_) = *i*^*^. Let [·]_*i*_denote the *i*-th column of a matrix, then 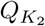 can be aligned to 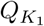 as a new matrix 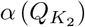 of the same size with columns

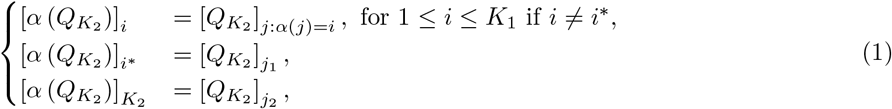

with the last column being the newly emerged cluster. This is essentially a relabeling of clusters in 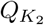. There are usually a range of *K* values, ordered increasingly as *K*_1_ ≤ *K*_2_ ≤ · · ·. Any modes between adjacent *K* values can be aligned in the same manner.

To align hard clustering results, we first convert the cluster labels from each run into an *N* × *K* binary membership matrix via one-hot encoding, and then perform clustering alignment on these matrices. Denote the optimal alignment by *α*^*^. Next, to quantitatively compare the aligned results, we relabel the clusters from the run with more clusters (*K*_2_ in this example) according to *α*^*^. Because *α*^*^ is many-to-one when *K*_2_ *> K*_1_, multiple *K*_2_ clusters may receive the same label, so that the number of unique labels matches that of the other run (*K*_1_). We then quantify disagreement in cell memberships using the normalized Hamming distance (NHD) between the (re-labeled) cluster assignments:

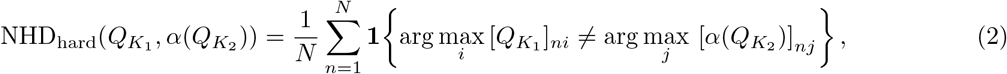

To compare clustering outputs from multiple hard clustering models, `ACE-OF-Clust` uses a two-level alignment scheme. In the first level (within-model alignment), we run each clustering model multiple times (e.g., 10 runs) and align the resulting outputs with `Clumppling`. This yields a small set of representative modes—potentially with different *K* values—that summarize each model’s clustering behavior. In the second level (across-model alignment), we pool the modes from all models and realign them, bringing all models’ outcomes into a common reference frame for direct comparison.

In mixed-membership clustering via `FastTopics` (Carbonetto, K. Luo, et al., 2023), the number of clusters are pre-specified by the user. We therefore fit models over a consecutive range of *K* values to systematically trace the emergence of new clusters as *K* increases. We then relabel clusters in the membership matrices according to the optimal alignment from `Clumppling`, both to examine the clustering hierarchy of cells and to consistently reorder clusters in the feature-level output matrices for comparing relative gene expression across clusters.

### 4.3 Identification of distinctive genes using feature-level matrices

In order to facilitate a biological understanding of clustering results, the mixed-membership clustering feature-level output matrix *P* has been used to identify the most distinctively differentially expressed genes in each cluster (Dey et al., 2017; Carbonetto, K. Luo, et al., 2023). A gene (feature) is considered “distinctive” if its expression in one cluster is significantly different from that in all other clusters. Dey et al. (2017) compute a distinctiveness score for each gene *j* based on the Kullback–Leibler (KL) divergence of Poisson distribution with parameter *p*_*jk*_:

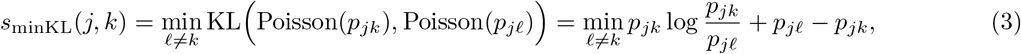

and select genes with the highest scores to interpret the gene ontologies driving each cluster. A later study from the same group, Carbonetto, K. Luo, et al. (2023), adopts a similar approach and identifies distinctive genes by estimating a least extreme log fold change (leLFC) score while accounting for uncertainty in the log fold change (LFC) estimates. The leLFC score is calculated as

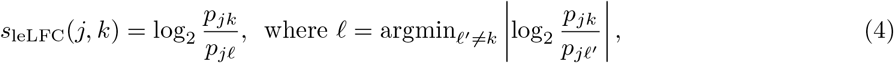

where a higher score indicates more distinctiveness for gene *j* in cluster *k*.

However, as noted by Carbonetto, K. Luo, et al. (2023), focusing on distinctiveness risks ignoring informative genes that exhibit common expression profiles among multiple clusters. To comprehensively characterize relationships among expression levels in matrix *P*—capturing not only gene distinctiveness in a single cluster but also the overall contribution of a gene to the entire clustering output—we propose constructing a novel “clustering profile” for each gene, from which we derive various metrics to summarize the gene’s contributing role, as detailed in the next section.

### 4.4 The clustering profile: a comprehensive snapshot of feature contributions to clustering output

From a feature-level matrix *P*, we construct a profile for each gene to summarize its contribution to the clustering, and we refer to this as its *clustering profile*. Gene *j*’s clustering profile comprises two vectors: a log fold change (LFC) vector *L*^*j*^ of length *K* − 1, and an index vector 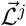 of length *K*.

For each gene *j*, we sort its corresponding *P* matrix entries {*p*_*j*1_, …, *p*_*jK*_} in increasing order as 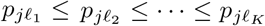, where the index vector 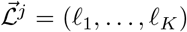 is a permutation of {1, …, *K*}. The entry 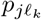 is the *k*-th smallest entry in row *j* of the *P* matrix. The LFC vector *L*^*j*^ contains LFC values computed as follows:

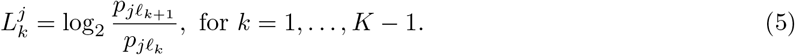

Since 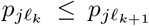, all 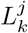 values are non-negative. The LFC vector and the index vector 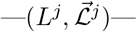 together form the clustering profile of gene *j*, which characterizes how gene *j* is relatively expressed across all *K* clusters by enabling two types of comparisons: a direct comparison of each cluster to “neighboring” clusters with the closest feature values for gene *j* (either higher or lower), and an indirect comparison of each cluster to all remaining clusters. Supplementary Fig. S3 illustrates the clustering profile constructed for a gene with varying relative expression in 5 clusters.

Metrics from the gene clustering profile 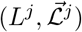 quantify different aspects of the gene’s contribution to the clustering, including its *distinctiveness score* (*s*_leLFC_ in Eq. 4; Carbonetto, K. Luo, et al. (2023)), as we detail below.

#### The distinctiveness of gene *j* in an individual cluster (denoted *leLFC*, the least extreme LFC)

The leLFC distinctiveness score *s*_leLFC_(*j, k*) for gene *j* in cluster *k* can be computed using the clustering profile vectors as

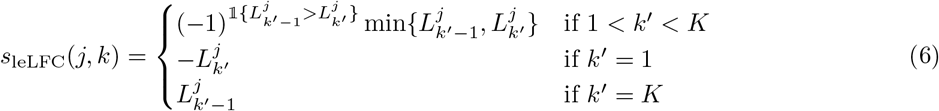

where 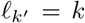 and 𝕝{·} is the indicator function. *s*_leLFC_(*j, k*) scores for all *k* can be extracted from the clustering profile following this process.

#### Maximum pairwise differentiation for gene *j* (denoted *sumLFC*)

Beyond distinctiveness in individual clusters, the clustering profile can also be used to evaluate a gene’s overall contribution to the clustering output. The maximum LFC observed between any pair of clusters for gene *j* is summarized as

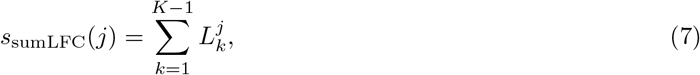

quantifying the divergence in relative feature levels—i.e., relative expression differences—across all clusters, with a larger value suggesting a stronger role in differentiating clusters, in comparison to other genes.

#### Largest separation gap in the relative expression profile for gene *j* (denoted *sepLFC*)

Another interesting metric is the maximum separation gap between adjacent clusters, *largest separation gap*, which we define as

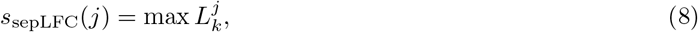

representing the largest “gap”, or difference, between adjacent values of sorted relative feature levels for gene *j*. The larger this gap is, the clearer the gene’s role is in distinguishing subsets of clusters. Determining which clusters are separated by the largest gap provides insight into the gene’s role in the differentiation of cell groups. For instance, *sepLFC* indicates whether the gene contributes to isolating a single cluster or distinguishing multiple clusters from the rest of the clusters. While the former replicates the idea behind distinctiveness scores (Section 4.3), the latter introduces the concept of “multi-cluster separation”, e.g., separating multiple clusters from others or distinguishing subgroups within broader groupings—a dimension not captured by distinctiveness measures.

### 4.5 Pipeline for identifying clustering-informative genes

`ACE-OF-Clust` identifies clustering-informative genes by examining two dimensions of the gene’s relative expression profile that are relevant for clustering: the total relative expression (explained below), and the largest separation gap (Eq. 8). Genes with high relative expression encoded in the matrix *P* are those contributing non-negligibly to the overall clustering. Genes with large separation gaps of log fold change, on the other hand, are those contributing to the differentiation of specific clusters. These two dimensions of the *P* matrix together inform us on a gene’s contributing role in clustering.

#### Genes with high relative expression

Recall that in mixed-membership clustering, we use all genes— excluding those with no variation in their expression profile. This is in contrast to the common practice of subsetting to highly variable genes (HVGs, for scRNA-seq) or spatially variable genes (SVGs, for ST) before downstream hard clustering. Selection of variable genes has been a focus in methodological development, yet benchmarking studies have raised concerns about the appropriate use of such selection; it remains under debate whether any single method provides a clear advantage, and what optimal usage conditions are (Yip et al., 2019; P. Yang et al., 2021; Adhikari et al., 2024; R. Zhao et al., 2024). The fact that most variable genes are not always most informative for clustering (Zihao Chen et al., 2024)—and that the optimal number of genes to use depends both on the data and on the algorithm—further underscores the complication of excluding genes from clustering.

Carbonetto, K. Luo, et al. (2023) proposed the mixed-membership clustering method `FastTopics` which can be applied to all sampled genes. By including all genes in clustering, the feature-level matrix *P* provides the option to filter out non-informative genes post-hoc after clustering. If a gene has low relative expression in all clusters, then it is likely not strongly influencing the clustering. We compute the total relative expression as a weighted sum of *P* entries for each gene *j*, denoted as 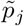, where the weight of each cluster, *w*_*k*_, is proportional to its total membership and *p*_*jk*_ is the *P* entry inferred for gene *j* in cluster *k*:

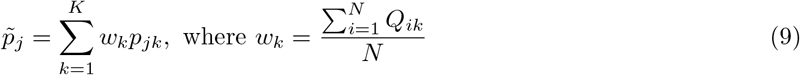

In applications in this study, we filtered genes by their 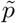 values—e.g., retaining the top 10%—to focus our subsequent analyses on the most promising candidate clustering-informative genes. This selection step is not guided by a gene’s impact on the statistical variation in expression data, but solely by its contribution to the clustering task. Another advantage of thresholding genes using feature-level matrix versus selecting HVGs pre-clustering is that the feature-level matrix is inferred simultaneously with the membership matrix, requiring less effort.

#### Genes that distinguish clusters

Genes with large output values for the summary metrics of the clustering profile (see Section 4.4), are key contributors to various aspects of the clustering output—e.g., they may be discriminating between distinct cell groups, or consistently contributing to the hierarchical branching of clusters. These genes offer insights into the biological relevance of clusters, such as the cell types, states, or functions that the clusters may represent; for instance, cell types are sometimes labeled using ontologies of top genes in them (e.g., T cells, B cells). Meanwhile, auxiliary annotations of cells in clusters (e.g., by tissue type and/or morphotype) generate hypotheses regarding biological activity of clustering-informative genes. The latter is particularly valuable in ST analysis, where the spatial information enables direct exploration of gene-cluster relationships within their tissue context.

We focus primarily on using the largest separation gap of log fold change (*sepLFC* in Eq. 8) to assess each gene’s contribution to clustering, since it inherently encompasses the scenario captured by the distinctiveness score in Eq. 4. To evaluate how different metrics in Section 4.4 identify clustering-informative genes and demonstrate the limitation of distinctiveness scores for highlighting genes that influence multi-cluster separation, we conduct a simulation analysis to show the advantages of *sepLFC* (Supplementary Methods). *sepLFC* provides insight from three perspectives: (1) all genes’ contribution in a single mode; (2) genes driving the emergence of new clusters; and (3) a specific gene’s role in the clustering hierarchy across multiple modes.

Within a single mode, by looking at the distribution of *sepLFC* of all genes (after thresholding by weighted *P* sum in Eq. 9), we group genes based on whether they are separating clusters, i.e., having a large *sepLFC* value, and how they separate clusters, i.e., the indices of clusters separated by the gap (Supplementary Fig. S12B). Genes that separate the same cluster(s) and exhibit similarly high *sepLFC* values likely play a shared and important role in shaping the clustering pattern of a given mode. These genes serve as markers for the corresponding cluster separation, and can be used to characterize the resulting clusters. Downstream analyses, such as gene set enrichment analysis, may also be performed on them.

For a specific gene of interest, we track how its *sepLFC*, along with the clusters separated by this gap, varies across the clustering hierarchy of modes across all *K* values. A gene’s role may shift throughout the clustering hierarchy, reflecting its varying importance in distinguishing cells at different scales. If the values are small in small-*K* modes, but become larger as *K* increases, this gene likely contributes more to the differentiation of fine-scale clusters that can be identified from the clustering profile index vectors.

## Data availability

Example data are available in our GitHub repository at https://github.com/xr-cc/ace-of-clust. The raw datasets used for clustering are publicly available from 10x Genomics (see Universal 3’ Gene Expression Dataset by Cell Ranger v1.1.0 (2016), Epi Multiome ATAC + Gene Expression dataset analyzed using Cell Ranger ARC 1.0.0 (2020), and Spatial Gene Expression Dataset by Space Ranger v1.1.0 (2020)).

## Code availability

The method is implemented as the Python package `ace-of-clust`. Source code: https://github.com/xr-cc/ace-of-clust. Documentation: https://xr-cc.github.io/ace-of-clust.

## Acknowledgments

This work was supported by the U.S. National Institutes of Health (NIH R35 GM139628 to SR and NIH R35 HG011939-01 to RS) and by the Data Science Institute at Brown University.

## Notes

https://github.com/xr-cc/ace-of-clust

### Competing Interest Statement

The authors have declared no competing interest.

### Summary of Updates

Section 2.3.3 added to present the gene set analysis; Supplementary files updated to include the corresponding methods and results.

