## Supplementary Materials for "Systematic clustering alignment and feature characterization for single-cell omics using ACE-OF-Clust"

### S1 Supplementary Materials

#### S1.1 The clustering alignment challenges and solutions

There are three challenges in the clustering alignment problem: (i) label switching, (ii) multi-modality, and (iii) difference in number of clusters.

Label switching arises when clusters with high similarity are assigned permuted labels, generating ambiguity across $K!$ possible permutations for $K$ clusters. Clustering results with labels switched are essentially equivalent. Multi-modality, on the other hand, occurs when output has fundamentally distinct clustering patterns, usually representing multiple global or local optima, where no label permutation can reconcile their differences. A third challenge arises in the comparison of results across varying model settings, most importantly the number of clusters $K$. Many clustering models require a pre-specified $K$ as input. However, identifying the optimal $K$ remains a long-standing challenge, as it is a model-dependent parameter without a well-defined "best" value, and researchers often iteratively generate clustering results for the same data using different $K$ values to select a suitable value based on some quantitative or qualitative assessment (Vayansky et al., 2020; Zhang et al., 2023). On the other hand, clustering models capable of inferring the number of clusters—such as community detection algorithms (e.g., Louvain (Blondel et al., 2008) and Leiden (Traag et al., 2019))—can also yield different $K$ when applied to the same dataset, which complicates the comparison and interpretation of clustering outputs across different models.

By resolving problems of label switching, multi-modality, and varying values of $K$, clustering alignment enables robust integration and comparison of clustering outcomes. Clustering alignment has already been studied in population genetics, where tools like `Structure` (Pritchard et al., 2000) and `Admixture` (Alexander et al., 2009), which apply model-based mixed-membership clustering to genotype data, are widely used in clustering analyses of genetic ancestry and population relationships. Multiple approaches have been developed to address clustering alignment, including `Clumpp` (Jakobsson et al., 2007), `Clumpak` (Kopelman et al., 2015), `Pong` (Behr et al., 2016), and `Clumppling` (X. Liu et al., 2024). The most recent method advancement, `Clumppling` (X. Liu et al., 2024), uses optimization and network strategies to efficiently detect and align all modes across different $K$ values.

#### S1.2 Clustering alignment objectives

To align two hard clustering runs with membership matrices $Q_{K_1}$ and $Q_{K_2}$, we adopt our "direct" approach implemented in `Clumppling` (X. Liu et al., 2024) which finds, for each cluster, the closest matching cluster in another run by minimizing the objective function

$$\min_{\alpha} D\Big(Q_{K_1}, \alpha(Q'_{K_2})\Big) = \min_{\alpha} \frac{1}{2N} \sum_{\ell=1}^{N} \sum_{k=1}^{K_2} \Big(q_{\ell k} - q'_{\ell \alpha(k)}\Big)^2, \tag{10}$$

where $N$ is the number of individuals (cells), $\alpha$ denotes the mapping of cluster indices from $[K_2]$ to $[K_1]$, and $q_{\ell k}$ is the membership of individual (cell) $\ell$ in cluster $k$, as described in the main text. The choice of the "direct" approach is because the $K$ values inferred by the community detection clustering algorithms are not guaranteed to be consecutive—meaning that adjacent $K$ values can differ by more than one, thereby disabling the "merge" option in `Clumppling`.

For the alignment of mixed-membership clustering runs with number of clusters $K_1$ and $K_2 = K_1 + 1$, we adopt the "merge" approach implemented in `Clumppling`; this approach minimizes the following dissimilarity between matrix $Q_{K_1}$ and some matrix $Q'_{K_2 \to K_1}$ that merges a pair of clusters from $Q_{K_2}$ so that two matrices have the same size:

$$\min_{\alpha, K_2 \to K_1} D\Big(Q_{K_1}, \alpha(Q'_{K_2 \to K_1})\Big) = \min_{\alpha, K_2 \to K_1} \frac{1}{2N} \sum_{\ell=1}^{N} \sum_{k=1}^{K_1} \Big(q_{\ell k} - q'_{\ell \alpha(k)}\Big)^2. \tag{11}$$

#### S1.3 3k Peripheral Blood Mononuclear Cells (PBMC3k) scRNA-seq data

##### S1.3.1 Overview of PBMC3k data

The single-cell RNA-seq benchmark dataset of Peripheral Blood Mononuclear Cells (PBMC) (Universal 3' Gene Expression Dataset by Cell Ranger v1.1.0, 2016) contains 2,700 single cells sequenced on the Illumina NextSeq 500. There are a total of 2,638 cells after basic processing (see next section for the quality control settings).

Out of the four implementations (the Leiden algorithm (Traag et al., 2019) and the Louvain algorithm (Blondel et al., 2008) in both `Seurat` (Satija et al., 2015; Butler et al., 2018) and `Scanpy`), `Seurat` uses the Louvain algorithm as default, and the tutorial of `Scanpy` uses Leiden. Following the pipeline taken by both software tutorials (Satija, 2023; Scanpy development team, 2025), we selected 995 highly variable genes (out of 32,738) and use the normalized, log-transformed, and scaled expression counts as input.

For mixed-membership clustering with `FastTopics` (Carbonetto, Luo, et al., 2023), expression counts of 13,714 genes with variation after initial preprocessing are used as input.

##### S1.3.2 Quality control, data processing and clustering settings for PBMC3k data

For hard clustering, the PBMC3k data was filtered following the `Seurat` tutorial (Satija, 2023). The raw count data with at least 3 cells and at least 200 genes has size 2,700 cells $\times$ 13,714 genes. The percentage of reads that map to the mitochondrial genome was calculated. Cells with unique feature counts over 2,500 or less than 200 and cells having 5% mitochondrial counts were then filtered, leaving a total of 2,638 cells. We then normalized the feature expression measurements for each cell by total expression and log-transformed the data. Next, feature selection was performed. To explicitly match the highly variable feature selection in `Seurat` (in R) and `Scanpy` (in Python), we chose the following filtering parameters for the functions:

```
# in Seurat
pbmc <- FindVariableFeatures(pbmc, selection.method = "mean.var.plot", num.bin = 20, mean.
    cutoff = c(0.1, 8), dispersion.cutoff = c(1, Inf))
// in Scanpy
scanpy.pp.highly_variable_genes(pbmc, flavor='seurat', n_bins=20, min_mean=0.1, max_mean=8,
    min_disp=1, max_disp=np.inf)
```

After feature selection, 995 variable features (genes) remain.

The transformed and filtered count data was then scaled to zero mean and unit variance with a max value of 10 for performing principal component analysis (PCA). A k-nearest neighbors (KNN) graph with $k = 15$ neighbors was constructed based on the euclidean distance in PCA space using the first 10 principal components (PCs). Based on this KNN graph, modularity optimization (a technique of community detection) was used to perform hard clustering, via either the Louvain algorithm (Blondel et al., 2008) or the Leiden algorithm (Traag et al., 2019). The resolution parameter that sets the "granularity" of the downstream clustering in the iterative clustering process of modularity optimization is set to 1.0 for both algorithms. For each algorithm in each software package, the clustering was performed 10 times with different random seeds.

For mixed-membership clustering using `FastTopics`, the pre-processing criteria are the same as for hard clustering: 13,714 genes with variation were included; cells with unique feature counts over 2,500 or less than 200 and cells having 5% mitochondrial counts were excluded. Genes are not filtered. The count matrix of size 2,638 cells $\times$ 13,714 genes was then used as input to the `FastTopics` function `fit_topic_model` with different specified number of clusters and parameters `numiter.main = 100, numiter.refine = 100, method.main = "em", method.refine = "scd"`. For each number of clusters $K$ from 2 to 13, clustering was performed 20 times with different random seeds.

**UMAP generation.** Note that the neighborhood construction used for UMAP is not the same setting as that used for clustering, therefore none of the clustering results (Fig. 1B, Supplementary Fig. S5A) clearly distinguish points in this UMAP, unlike in the reference result from `Scanpy`'s tutorial (Scanpy development team, 2025) (Fig. 1A). The UMAP was performed following `Scanpy`'s tutorial (Scanpy development team, 2025), where the KNN graph was constructed with 10 neighbors and top 40 PCs.

##### S1.3.3 Cell types in PBMC3k

The eight annotated cell types from the `Scanpy` tutorial (Scanpy development team, 2025) discussed in the main text (Fig. 1A) are: CD4 T cells (Cluster of Differentiation 4 T lymphocytes), CD8 T cells (cytotoxic T lymphocytes), B cells (B lymphocytes), NK cells (natural killer cells), FCGR3A+ monocytes (Fc gamma receptor IIIa-positive monocytes), CD14+ monocytes (Cluster of Differentiation 14-positive monocytes), dendritic cells, and megakaryocytes.

#### S1.4 Human Breast Cancer (HBC) ST data

##### S1.4.1 Overview of HBC data

The HBC gene expression data includes a total of 36,601 genes over 3,798 spots for one sampled slide. Mixed-membership clustering was performed on all 24,923 genes with variation in read count (11,678 genes show zero variation in read count and were discarded before analysis). For hard clustering analysis, we used a subset of 3,000 highly variable genes (HVGs) selected via `Scanpy`'s variable selection. This subset of HVGs is also used to examine whether highly variable genes are more likely to be clustering-informative.

##### S1.4.2 Quality control, data processing and clustering settings for HBC data

For hard clustering, we used the subset of 3,000 highly variable genes (selected using `Scanpy`'s function `scanpy.pp.highly_variable_genes`), resulting in the input data matrix of size 3,798 spots $\times$ 3,000 genes. For both `Seurat` and `Scanpy`, the data was normalized, log-transformed, and scaled to have zero mean and unit variance, following the same preprocessing steps used for the PBMC3k data. For hard clustering, a KNN graph was constructed using 20 neighbors and the top 10 PCs. Both the Louvain and Leiden algorithms (Blondel et al., 2008; Traag et al., 2019) were run with a resolution parameter of 1.0, each for a total of 10 runs with different random seeds for each run.

For mixed-membership clustering, the count matrix consisting of 3,798 spots and all 24,916 genes with variation in gene count across spots was used as input to `FastTopics`. The parameters for function `fit_topic_model` are the same as those used for PBMC3k data: `numiter.main = 100, numiter.refine = 100, method.main = "em", method.refine = "scd"`. For each number of clusters $K$ from 3 to 9, clustering was performed 20 times with different random seeds.

#### S1.5 10k Peripheral Blood Mononuclear Cells (PBMC10k) scRNA-seq and ATAC-seq data

##### S1.5.1 Overview of PBMC10k data

The PBMC10k multiome dataset (Epi Multiome ATAC + Gene Expression dataset analyzed using Cell Ranger ARC 1.0.0, 2020) contains paired scRNA-seq and scATAC-seq profiles from cryopreserved PBMCs of a healthy 25-year-old female donor, generated by 10x Genomics (AllCells). It includes 11,909 cells, 36,601 genes, and 108,377 peaks (chromatin accessible regions). We processed and subset each modality following the `muon` tutorial (Bredikhin et al., 2022; Bredikhin, 2025) to obtain matched cells. For both hard and mixed-membership clustering, we followed the tutorial's highly variable feature selection procedure (genes for RNA-seq and peaks for ATAC-seq) to reduce computational cost—particularly for ATAC-seq—despite our earlier observation that this filtering step can miss features that are informative for clustering.

##### S1.5.2 Quality control, data processing, and clustering for PBMC10k data

For RNA, we retained genes detected in $\geq 3$ cells and cells with 200–5,000 detected genes, total counts $\leq 15{,}000$, and mitochondrial fraction $\leq 20\%$, leaving 26,349 genes. We then selected 3,026 highly variable genes (HVGs) using `sc.pp.highly_variable_genes(rna, min_mean=0.02, max_mean=4, min_disp=0.5)`, and used these HVGs for both hard and mixed-membership clustering. UMAP and clustering follow the `muon` tutorial, using:

```
sc.pp.neighbors(rna, n_neighbors=10, n_pcs=20)
sc.tl.leiden(rna, resolution=0.5, flavor="igraph", directed=False, n_iterations=2)
```

```
sc.tl.umap(rna, spread=1., min_dist=.5, random_state=11)
```

After manually matching clusters to the tutorial's annotated groups and removing noisy clusters as recommended, we obtained 10,908 cells across 13 cell types. We also ran mixed-membership clustering on RNA-seq output using the full set of 26,349 genes and observed the same HVG vs. non-HVG patterns in feature-level metrics (Fig. S14).

For ATAC, we retained peaks detected in $\geq 10$ cells and cells with 2,000–15,000 detected peaks and total counts between 4,000 and 40,000. This yields 10,069 cells and 106,086 peaks, from which 14,896 highly variable peaks were selected via `sc.pp.highly_variable_genes(atac, min_mean=0.05, max_mean=1.5, min_disp=.5)` and used for both hard and mixed-membership clustering. Clustering mirrors the RNA pipeline (PCA on log-normalized counts, neighborhood graph construction, and Leiden clustering). After annotation and removal of noisy clusters as in the tutorial, 9,815 ATAC cells remained. Intersecting modalities yields 9,554 shared cells and 132,435 total features (26,349 genes + 106,086 peaks).

##### S1.5.3 Cell types in PBMC10k

The 13 annotated cell types from the `muon` tutorial (Bredikhin, 2025) discussed in the main text (Fig. S16A) are: CD4+ memory T (Cluster of Differentiation 4–positive memory T lymphocytes), CD4+ naive T (Cluster of Differentiation 4–positive naive T lymphocytes), CD8+ naive T (Cluster of Differentiation 8–positive naive T lymphocytes), CD16 mono (Cluster of Differentiation 16–positive monocytes; non-classical monocytes), NK (natural killer cells), intermediate mono (intermediate monocytes), CD14 mono (Cluster of Differentiation 14–positive monocytes; classical monocytes), mDC (myeloid dendritic cells; conventional dendritic cells), memory B (memory B lymphocytes), MAIT (mucosal-associated invariant T cells), CD8+ activated T (activated Cluster of Differentiation 8–positive cytotoxic T lymphocytes), naive B (naive B lymphocytes), and pDC (plasmacytoid dendritic cells).

#### S1.6 Simulation study: genes with different contributing roles in clustering

**Overview.** To evaluate how different clustering profile metrics in Section 4.4 identify genes contributing to clustering in different ways—particularly genes that are distinctive for a single cluster versus those that influence the differentiation of multiple clusters—we conducted a simulation analysis.

We generate a synthetic expression matrix $X$ by multiplying a count-adjusted membership matrix $H$ and a feature-level matrix $W$, separately simulated, following the matrix factorization formulation of the mixed-membership model (as in Carbonetto, Sarkar, et al. (2021)). We choose $N = 1000$ cells, $G = 3000$ genes, and $K = 4$ clusters. To generate genes with different roles in clustering, we explicitly simulate genes to have feature-level values according to three scenarios: (1) similar in all clusters, (2) different in one cluster from the other three, or (3) different in cluster 1 and 2 from those in cluster 3 and 4. This creates coarse differentiation in simulated gene expression profiles between clusters 1,2 and clusters 3,4, as well as fine-scale differentiation among the individual clusters. Simulated genes belonging to scenario 1 are genes that are not informative for the clustering. Those belonging to scenario 2 are distinctive genes. Those belonging to scenario 3 represent genes that contribute to the differentiation of clusters, but not distinct to a single cluster.

**Simulation parameterization** The count-adjusted membership matrix entries in $H$ are simulated as $h_{ik} = c_{ik}t_i$, where $t_i$ is the total count for each synthetic cell $i$, and $c_{ik}$ values encode the membership fraction of each cell $i$ in cluster $k$. We use $\mathcal{N}(\mu, \sigma^2)$ to denote the normal distribution with mean $\mu$ and standard deviation $\sigma$. The total count is sampled as $t_i = \exp(u_i)$ with $u_i \sim \mathcal{N}(9.7, 0.8^2)$. This sampling distribution was selected based on resemblance to the distribution of total gene count for spots in real HBC data. Each cell is simulated to have non-zero membership in $K'$ of $K$ clusters, where this number is drawn from $K' \in \{1, \cdots, K\}$ with probability proportional to $2^{-K'}$. These $K'$ clusters are uniformly sampled from $\{1, \cdots, K\}$. If $K' = 1$, $c_{ik} = 1$ for the selected cluster and $c_{ik} = 0$ for other clusters; if $K' > 1$, $c_{ik} \sim \mathrm{Dir}(\alpha_{K'})$ for the $K'$ selected clusters and $c_{ik} = 0$ otherwise. $\mathrm{Dir}(\alpha)$ denotes the Dirichlet distribution parameterized by $\alpha$, and here we simply set it as a vector of length $K'$ with all ones, corresponding to a flat Dirichlet distribution. Note that $\sum_{k=1}^{K} c_{ik} = 1$ for all $i$.

The feature-level matrix $P$ is simulated in a slightly different manner. By examining the $P$ matrices generated on the real HBC spatial transcriptomics data (also used in Section 2.3), we observe that log base-10 values in $P$ columns, that is, the log of feature-level values in each cluster, roughly follow a bimodal distribution (see also Supplementary Fig. S12). Based on this observation, for each synthetic gene $j$, we draw log feature values from two normal distributions: $a_j \sim \mathcal{N}(a_0, \sigma^2_{a_0})$, where $a_0 \sim U[-13, 12.5]$ and $\sigma_{a_0} = (14 + a_0)/3$, and $b_j \sim \mathcal{N}(b_0, \sigma^2_{b_0})$, where $b_0 \sim U[-5, -3.5]$ and $\sigma_{b_0} \sim U[0.8, 1.2]$. Let $g_j \in \{0, 1, 2, 3, 4\}$ with probability (0, 0.1, 0.4, 0.3, 0.2). For each gene $j$, its role in the clustering is determined by $g_j$: (1) $g_j = 0$ or 4 (a total probability of 0.2) corresponds to the first simulation scenario where the gene has similar feature-level relative expression values in all clusters; (2) $g_j = 1$ or 3 (a total probability of 0.4) corresponds to the second simulation scenario where the gene has different feature-level relative expression values in one cluster from the rest; (3) $g_j = 2$ (a probability of 0.4) corresponds to the third simulation scenario where the relative expression values in clusters 1,2 are different from those in clusters 3,4. For each gene $j$, out of the $K$ clusters, we randomly choose $g_j$ to have feature value $w_{jk} = 10^{b_j}$, and the rest have $w_{jk} = 10^{a_j}$. After setting the $w_{jk}$ values, we re-normalize each column by $w_{jk} \leftarrow w_{jk} / \sum_{j=1}^{M} w_{jk}$ to make sure each column of $W$ sums to one.

A simulated expression matrix $X = HW^T$ is then generated, on which the clustering methods are applied. Using the clustering results, in Section S1.7, we next evaluated how different scores, or summary values of the gene clustering profile, can be used to highlight genes with different roles in clustering.

#### S1.7 Simulation results: feature-level metrics capture different aspects of a gene's contribution to clustering

To assess the utility of gene distinctiveness scores described in Section 4.3, we conducted a simulation analysis comparing their performance against the more comprehensive set of summary values we derived from the clustering profile. Our analysis highlights the limitations in using distinctiveness scores to identify clustering-informative genes.

Our simulation procedure for an expression matrix is detailed previously in Section S1.6. We generated 20 simulated expression matrices with 3000 genes, and for each matrix, `FastTopics` (Carbonetto, Luo, et al., 2023) was run 5 times for $K$ from 2 to 4, giving a total of 100 clustering results for each $K$. For each clustering result, we compute the distinctiveness scores (Eqs. 3 and 4) as well as the summary values $sumLFC$ (Eq. 7) and $sepLFC$ (Eq. 8) after constructing the clustering profiles for the genes.

In our simulations for each gene, we chose $g_j$ clusters to have higher feature values than the rest, with $g_j = 1, 2, 3, 4$ (note that we set the case $g_j = 0$ to zero probability). We grouped genes based on their simulated type characterized by how many clusters ($a$) having different feature values than the rest ($b$), encoded in a "$a - b$" notation, with $a \leq b$. The above four choices of $g_j$ correspond to the simulated gene type of "1-3", "2-2", "1-3", and "0-4", respectively.

In the clustering results for each simulation, there are two types of scores for each gene: the scores $s_{\text{minKL}}(j,k)$ and $s_{\text{leLFC}}(j,k)$ with one value per cluster, and the scores $s_{\text{sumLFC}}(j)$ and $s_{\text{sepLFC}}(j)$ with a single value for all clusters. To simplify the comparison of scores, for $s_{\text{minKL}}(j,k)$ and $s_{\text{leLFC}}(j,k)$ we got a single value (the maximum value) from the $K$ scores across all clusters: the maximum $s_{\text{minKL}}$, and the maximum absolute $s_{\text{leLFC}}$ (the absolute was taken to avoid negative values). For $s_{\text{sepLFC}}$, we further split the genes based on the separation pattern—the number of clusters separated by the largest LFC gap. For $K$ clusters, the possible separation patterns are $(1, K-1)$ up to $(\lceil K/2 \rceil, K - \lceil K/2 \rceil)$.

The distributions of these scores across all tested $K$ values are visualized in Fig. S4), providing comparative analyses of the scores. In the grouped box plot for each score, there are a total of $3000 \times 100$ points, except for the plot for $s_{\text{sepLFC}}$ which is further split into $\lceil K/2 \rceil$ subplots (last panel(s) in each row). Not surprisingly, at $K = 4$ where the number of clusters matches the simulated ground truth, two distinctiveness scores $s_{\text{minKL}}$ and $s_{\text{leLFC}}$ are high for genes simulated to have a single cluster different from the other three (1-3), and lower for other genes. These patterns can also be obtained using the $s_{\text{sepLFC}}$ values separating a single cluster ("$sepLFC(1-3)$"). However, both genes with different expression in two clusters versus the other two clusters (2-2) and genes with no difference among the clusters (0-4) are missed by the distinctive scores for $K = 4$ (Fig. S4 last row, first two columns). While the (0-4) genes are not necessarily informative, the (2-2) genes remain valuable for defining cluster structure and should still be prioritized for analysis. $s_{\text{sepLFC}}$ values separating two clusters ("$sepLFC(2-2)$") exactly capture this missing piece.

At $K = 2$, all LFC-based scores are the same, since there is a single LFC value between two clusters for each gene. All LFC-based scores as well as $s_{\text{minKL}}$ are highest for genes under simulated scenario 2-2. Genes with a single distinctly expressed cluster (1-3) exhibit higher scores compared to those with no difference in the relative expressions of all clusters (0-4). Similar patterns exist at $K = 3$, except that scores for genes with simulated type 1-3 are further elevated, specifically for the LFC-based scores. When $K$ used in clustering is smaller than the true number of clusters used to generate the data, many genes of simulated type 2-2 serve as genes that help isolate a single cluster (as a combination of two simulated clusters) from the other, thereby having large distinctive scores.

#### S1.8 Additional results on PBMC3k feature analysis

We ran `FastTopics` (Carbonetto, Luo, et al., 2023) 20 times for each of $K$ from 2 to 13 on the PBMC3k data, and aligned all the 240 clustering runs. The structure plots of detected modes alongside the alignment patterns—i.e., how clusters from a $K$-mode are matched to those in a $(K+1)$-mode—are shown in Supplementary Fig. S6, the these aligned mixed-membership results allow us to observe how new clusters emerge. For example, when $K$ increases from 2 to 3, cells that predominantly correspond to CD8 T and NK cells in the reference shift a large portion of their membership to the newly formed cluster. Similarly, when $K$ increases from 3 to 4, the cluster matching B cells in the reference splits away from the cells mostly labeled as CD4 T.

We then identified genes that are informative for the clustering by examining both the total relative expression (weighted $P$ sum in Eq. 9) and the largest separation gap (*sepLFC* in Eq. 8). Feature analyses for PBMC3k are shown in Supplementary Figs. S7 and S8. We ranked genes in each mode by the largest separation gap—which reflects their power in separating sets of clusters—and highlighted the top genes that also exhibit substantial total relative expression (having a weighted $P$ sum value among the top 10%). We also examined the joint distribution of genes with respect to the two metrics (weighted $P$ sum and *sepLFC*) in each mode, and focused on outliers that contribute most to the clustering. For a specific gene of interest, using its clustering profile metrics, we tracked how its clustering role varies across all analyzed modes (Supplementary Fig. S8). For instance, the gene *MALAT1* shows a consistently moderately high weighted $P$ sum value and has limited contribution to separating clusters when $K$ is small, but exhibits large *sepLFC* values as $K$ increases, primarily distinguishing multiple clusters (Supplementary Fig. S8).

#### S1.9 Cluster emergence in mixed-membership clustering of HBC ST data

On HBC data, we performed mixed-membership clustering using `FastTopics` (Carbonetto, Luo, et al., 2023) for $K = 3 - 9$, and show the aligned mixed-membership clustering results for $K = 3 - 7$ in Figs. 2D,E. See Supplementary Fig. S9 for selection of this $K$ range. Each mode is represented by the clustering memberships from a representative run. Across these five $K$ values, we identified eight modes. For example, "K6M2" denotes the second mode from clustering runs with $K = 6$. Spatial visualizations of spot-level memberships are shown in Fig. 2E, with the corresponding alignment patterns in Fig. 2D. Viewing membership coefficients in their spatial context allows us to track how spot expression profiles differentiate as $K$ increases. In two annotated IDC regions (IDC 2 and 4; 2A), most spots have high membership in the same cluster (pink) at small $K$. Only at larger $K$ (e.g., K6M2 and K7M2, Fig. 2E) do the dominant memberships in these regions separate into distinct clusters (pink vs. yellow in K6M2; pink vs. dark blue in K7M2). Spots in the healthy morphotype consistently show similar memberships to those in the tumor-edge morphotype (cluster 2, green), consistent with the difficulty of separating these regions by hard clustering (Fig. 2B). In contrast, tumor edge 3 exhibits distinct signals: it forms a standalone cluster at $K = 6$ (cluster 6, brown) and further splits into two clusters in K7M1 (clusters 6 and 7), suggesting unique transcriptomic features in this annotated region (Fig. 2E). The same cluster emergence may occur at a different $K$. For example, in K6M2, clusters 5 (yellow) and 3 (purple) split from cluster 3 (purple) in K5M1; a similar split reappears in K7M2, where clusters 5 (dark blue) and 3 (purple) both align to cluster 3 in K6M1 (purple).

Many spots also exhibit continuous variation in memberships. In IDC 5, spots begin to show substantial membership in a new cluster (cluster 5, yellow; except in K6M2) at $K = 5$, but their memberships remain split between this cluster and an existing cluster (cluster 1, orange; prevalent in DCIS/LCIS 3). In addition, the blue cluster shares membership with the orange and yellow clusters in some spots (particularly in

DCIS/LCIS 4), providing further evidence for a non-discrete spectrum of transcriptomic heterogeneity.

#### S1.10 Assessing the influence of gene selection on clustering output

We performed mixed-membership clustering on the HBC dataset using the full set of genes (with variation in read count). To assess how restricting analysis to a subset of genes affects clustering analyses outcomes, we examined clustering results on the same HBC dataset using two gene subsets: (1) highly variable genes selected through a popular implementation, and (2) clustering-informative genes identified based on our proposed metrics.

In Supplementary Fig. S14, we compare the distribution of total relative expression (weighted $P$ sum in Eq. 9) and largest separation gap (*sepLFC* in Eq. 8) for 3,000 highly variable genes (HVGs) against the complement of non-highly variable ones (non-HVGs)—out of the 24,916 genes assayed containing variation in read count. The highly variable gene selection is performed via the implemented function in `Scanpy` (Wolf et al., 2018), which reproduces the R-implementation of `Seurat` (Satija et al., 2015). Genes are placed into several bins based on their average expression across all cells, and within each bin top genes are selected based on a dispersion measure. We show the distributions of weighted $P$ sum and *sepLFC* for modes K3M1 and K7M2 in Supplementary Fig. S14 (both modes included in Fig. 2E). Although the quartiles of weighted $P$ sum and *sepLFC* are slightly lower for HVGs than for non-HVGs in both modes, this difference is minimal. Notably, both groups exhibit a bimodal distribution for their *sepLFC* values, with a small subset of genes showing substantially higher values, indicating that these genes are much more strongly clustering-informative than the others. Thus, discarding non-HVGs in mixed-membership clustering overlooks genes that are equally informative for clustering as the HVGs.

We also compared clustering results using all genes against those obtained using a subset of genes selected based on the two clustering-informative metrics we proposed in Section 4.5. We re-performed clustering with $K = 4$ and $K = 5$ using only genes with top 10%, 20%, and 30% of total relative expression (weighted $P$ sum in Eq. 9). The clustering results, shown in Supplementary Fig.S15A, align well to modes K4M1 and K5M1 from clustering using all genes (Fig. 2E); these two modes are highlighted in Supplementary Fig. S15A. When genes were further thresholded based on their largest separation gap (*sepLFC* in Eq. 8), clustering results remain unchanged when leaving out genes with small *sepLFC* value (e.g. $sepLFC \leq 2$ for K4M1 and $sepLFC \leq 1$ for K5M1), as shown in Supplementary Fig. S15B, and a higher *sepLFC* threshold usually yields different results from clustering using all genes. In that sense, clustering is not sensitive to genes deemed non-clustering-informative (based on the metrics weighted $P$ sum and *sepLFC*), whereas genes with non-negligible *sepLFC* values have considerable impact on clustering output. Excluding non-clustering-informative genes does not alter the overall clustering structure.

#### S1.11 Variability in hard clustering across omic data types

We performed hard clustering on both scRNA-seq and scATAC-seq data using the same four models, with 50 runs each. The aligned major modes are shown in Supplementary Fig. S16A, with cells embedded in a 2D UMAP generated following the `muon` tutorial (Bredikhin, 2025), analogous to the PBMC3k scRNA-seq example in Supplementary Fig. 1B. Reference cell-type annotations from the `muon` tutorial (Bredikhin, 2025) are shown in Supplementary Fig. S16C; because we could not fully reproduce the tutorial clustering, we assigned labels by manual inspection and closest matching to the tutorial. Rather than comparing modes to the tutorial clusters, Supplementary Fig. S16B compares each major mode to an internal reference: K16M1 from scRNA-seq `Seurat` Leiden, which has the largest mode size (27 out of 50 runs). For each non-reference mode, only cells that differ from the reference are colored, and the total difference is reported in the label. As in the PBMC3k scRNA-seq and HBC ST examples, cluster memberships vary across clustering models, and the variability is, as expected, greater across different omics than within the same omic.

Stratifying differences by annotated groups (Supplementary Fig. S16D) shows that variability is cell-type dependent. Many cells labeled intermediate mono, as well as some labeled as CD4+ naive T, CD8+ naive T, or CD14 mono, differ substantially between ATAC-seq and the RNA-seq reference. Several smaller annotation groups also exhibit large differences in a subset of ATAC-seq modes, consistent with reduced stability for rare populations. These results underscore the importance of reliable cell-type identification, particularly for downstream analyses that depend on these groupings.

#### S1.12 Gene set analysis

Recall that matrix $P : G \times K$ is the relative gene expression matrix for a given clustering mode with $K$ clusters and $G$ genes, where $P_{jk}$ encodes the relative expression of gene $j$ in cluster $k$. Let $sepCls = (sep\mathcal{L}, sep\mathcal{H})$ be the separating bipartition (i.e., two sets of clusters) identified via *sepLFC* for this mode. Let $\mathcal{S} \subseteq 1, \ldots, G$ denote a gene set (abbreviated as "GS") of size $|\mathcal{S}|$. The submatrix $P_{\mathcal{S}} : |\mathcal{S}| \times K$, which contains the rows of $P$ corresponding to this gene set, is the focal point of the gene set analysis.

##### S1.12.1 Gene set enrichment analysis

Gene set enrichment analyses are performed on the *gene-set average relative expression vector*

$$\bar{p}^{\mathcal{S}} = \frac{1}{|\mathcal{S}|} \sum_{j \in \mathcal{S}} P_{j\cdot} \ \in \ \mathbb{R}^K, \tag{12}$$

which summarizes the collective relative expression of all genes in the set across clusters.

**Null distribution.** A permutation null is generated by drawing $B = 10{,}000$ random gene sets of the same size $|\mathcal{S}|$ (without replacement from the $G$ genes), each producing a null gene-set average relative expression vector $\bar{p}^{(b)} \in \mathbb{R}^K$. Hypothesis tests compare observed statistics to this shared null, yielding empirical $p$-values (with a small correction) of the form

$$p_{\text{emp}} = \frac{1 + \#\{b : \text{null} \geq \text{obs}\}}{B + 1}. \tag{13}$$

**Relative expression ($P$) enrichment.** For each cluster $k$, the observed $\bar{p}_k^{\mathcal{S}}$ is compared to the null distribution $\{\bar{p}_k^{(b)}\}_{b=1}^{B}$. The empirical one-sided $p$-value (Eq. 13) is reported, quantifying the evidence that the relative expression of that gene set is greater than the null value.

**Largest separation gap (*sepLFC*) enrichment.** Clusters are sorted by $\bar{p}_k^{\mathcal{S}}$ in ascending order, and recall that *sepLFC* is the largest log-fold-change gap in this sorted sequence. Clusters below the gap form the low group $sep\mathcal{L}$ and clusters above form the high group $sep\mathcal{H}$. Statistical significance is assessed against two null distributions:

1. *sepLFC of the null*: the largest separation gap computed independently for each null vector $\bar{p}^{(b)}$, testing whether the largest separation for the gene set exceeds what any random set of the same size achieves.

2. *LFC across sepCls of the gene set*: the LFC between the fixed bipartition of the gene set, $(sep\mathcal{L}, sep\mathcal{H})$, evaluated on each null set,

$$\delta^{(b)} = \log_2 \min_{k \in sep\mathcal{H}} \bar{p}_k^{(b)} - \log_2 \max_{k \in sep\mathcal{L}} \bar{p}_k^{(b)}, \tag{14}$$

testing whether the gene set's separation at the same cluster partition exceeds chance. This $\delta^{(b)}$ value can be negative if the cluster in $sep\mathcal{H}$ with the lowest relative expression has a value lower than that of the cluster in $sep\mathcal{L}$ with the highest relative expression.

Empirical $p$-values (Eq. 13) are computed for both comparisons.

**Enrichment in a single mode and across multiple modes.** The enrichment analyses are performed independently for each clustering mode, and the results can be visualized for a single mode or compared across modes. For relative-expression enrichment (Supplementary Fig. S19A), the $P$-enrichment $p$-values for each cluster across all modes are displayed as bar charts. For largest-separation-gap enrichment (Supplementary Figs. S19B-C), the null distribution for each mode is shown as one row in a heatmap, with modes stacked vertically; darker cells indicate higher density. The observed *sepLFC* for the gene set is marked by a red dot, and its empirical $p$-value is annotated alongside it, with significance indicated by an asterisk when the $p$-value falls below the chosen threshold (0.05 by default).

##### S1.12.2 Per-gene decomposition of the *sepLFC* and the associated *sepCls*

**Individual LFC contribution of each gene to the *sepLFC* of the gene set.** Within a given gene set, we are also interested in the behavior of individual genes in terms of relative expression and cluster-separation patterns, especially the genes with the highest relative expression and genes with the largest individual *sepLFC* for the gene set's *sepCls*. Per-gene analyses operate directly on the rows of $P_{\mathcal{S}}$.

It is not straightforward to decompose the contribution of each gene in the gene set to the *sepCls* pair $(sep\mathcal{L}, sep\mathcal{H})$ identified by the *sepLFC*, because log fold change is not linearly related to the inclusion or exclusion of each relative-expression value $P_{jk}$, and the clusters separated by each gene's *sepLFC* can differ from the gene set's *sepCls*. For each gene $j \in \mathcal{S}$, two measures of the individual LFC contribution are computed with respect to the gene-set bipartition $(sep\mathcal{L}, sep\mathcal{H})$ (Supplementary Figs. S21A-B and S23A-B). The *mean* measure averages relative expression within each group,

$$\ell_j^{\text{mean}} = \log_2\left(\frac{1}{|sep\mathcal{H}|}\sum_{k\in sep\mathcal{H}} P_{jk}\right) - \log_2\left(\frac{1}{|sep\mathcal{L}|}\sum_{k\in sep\mathcal{L}} P_{jk}\right), \tag{15}$$

while the *extreme* measure uses the most conservative pair of values:

$$\ell_j^{\text{extreme}} = \log_2 \min_{k\in sep\mathcal{H}} P_{jk} - \log_2 \max_{k\in sep\mathcal{L}} P_{jk}. \tag{16}$$

Many genes in the set need not have large $\ell_j^{\text{mean}}$ or $\ell_j^{\text{extreme}}$ values. Neither $\ell_j^{\text{mean}}$ nor $\ell_j^{\text{extreme}}$ is required to be positive; some genes may even oppose the gene-set-level separation. Nevertheless, some genes must collectively contribute substantially to the formation of the set's *sepCls*.

To place the gene set in the context of all genes examined in the analysis, genes outside the gene set that share the same (unordered) bipartition according to their own *sepLFC* are also ranked by their *sepLFC* values (Supplementary Figs. S21D and S23D). Here, "unordered" means that the bipartition is considered the same as long as the same clusters remain in $sep\mathcal{L}$ and $sep\mathcal{H}$; the specific ordering of clusters within either side does not matter. The highest-ranked non-gene-set genes provide complementary information about the factors contributing to the major clustering-separation pattern beyond the gene set of interest, and may therefore be worth further investigation for their relationships with genes in the set.

**Separation of each pair of clusters in *sepCls*.** As in the gene-level analysis, we also compute a per-gene *sepLFC* and the corresponding optimal separating bipartition independently for each gene, again denoted by $sepCls = (sep\mathcal{L}, sep\mathcal{H})$. Among the genes $j \in \mathcal{S}$ whose individual optimal bipartition matches $(sep\mathcal{L}, sep\mathcal{H})$ (up to permutation within each set), we identify, for each pair of clusters across $sep\mathcal{L}$ and $sep\mathcal{H}$, the top $n$ genes ranked by individual *sepLFC*. These are the genes whose largest separation gap occurs between a cluster in $sep\mathcal{L}$ with lower relative expression and a cluster in $sep\mathcal{H}$ with higher relative expression. These genes are not necessarily—and often are not—the genes with the highest relative expression in each cluster. This suggests that genes with the highest overall expression are not always the ones that differ most strongly across clusters and therefore drive cluster separation.

The separating bipartition is visualized as a bipartite graph (Supplementary Figs. S20F and S22F), with clusters in $sep\mathcal{H}$ shown as top nodes and clusters in $sep\mathcal{L}$ as bottom nodes. For each pair $(k \in sep\mathcal{H}, k' \in sep\mathcal{L})$, an edge connects cluster nodes $k$ and $k'$, with width proportional to $\sum_{j\in\mathcal{S}} \ell_j^{(kk')}$, where $\ell_j^{(kk')}$ denotes the gene-specific *sepLFC* for that cluster pair. Each edge is further divided into gene-specific segments, with segment lengths proportional to each gene's contribution relative to the total across all top-$n$ genes and segment color reflecting the gene-specific *sepLFC* value.

### S2 Supplementary Figures

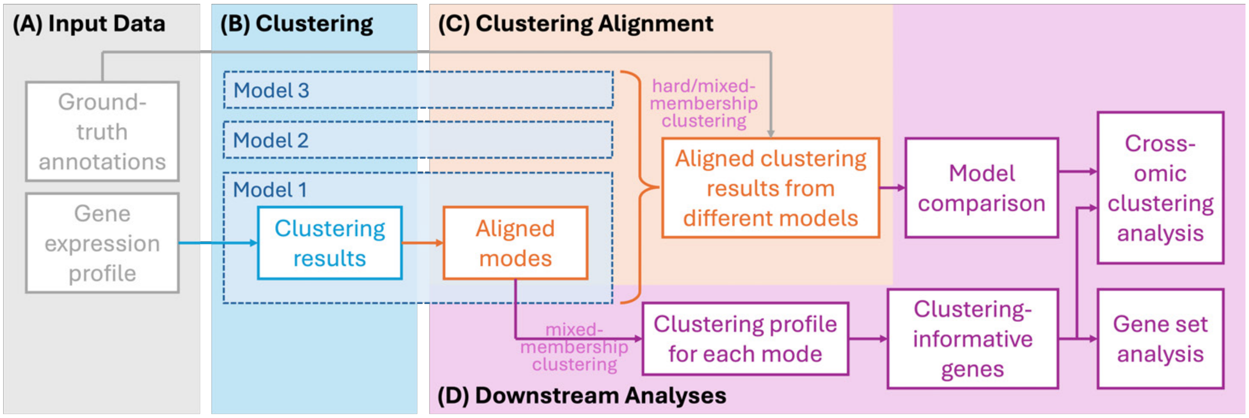


Figure S1: **Schematic overview of the `ACE-OF-Clust` workflow.** **(A)** Input consists of scRNA-seq or ST gene expression profiles (or other omic features); when available, ground-truth annotations are reserved for downstream model comparison. **(B)** Hard or mixed-membership clustering is run repeatedly, producing multiple membership matrices and potentially varying $K$. **(C)** `Clumppling` (X. Liu et al., 2024) aligns clustering outputs, identifies modes for each $K$, and represents each mode by a representative run. **(D)** The aligned results enable qualitative and quantitative model comparison. For mixed-membership clustering, feature-level metrics summarizing gene-specific clustering profiles prioritize clustering-informative genes. These feature metrics further support gene set enrichment analysis and cross-omic feature-pair analysis.

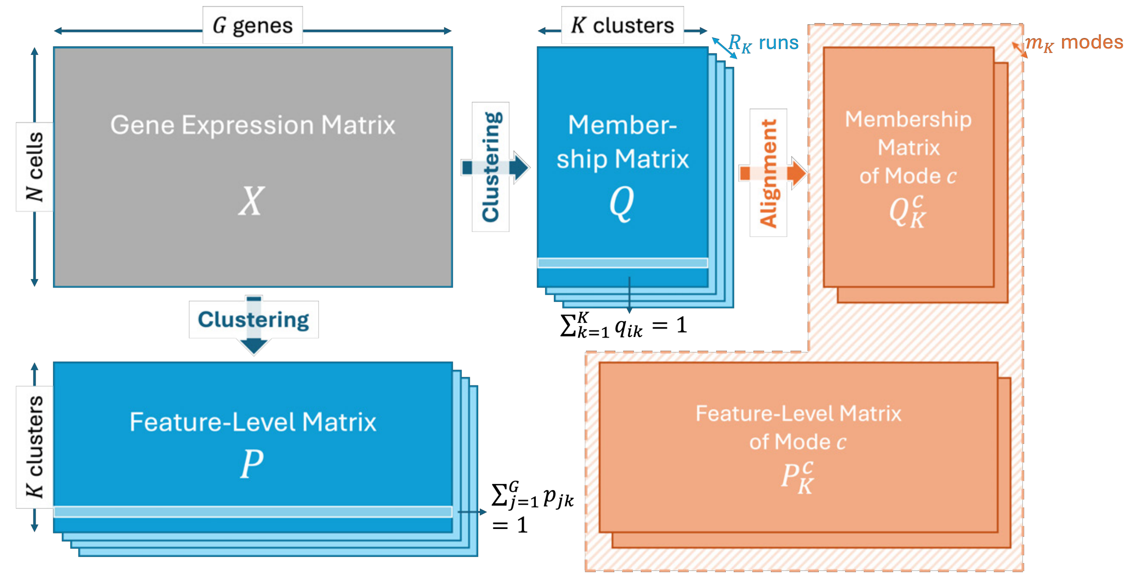


Figure S2: **A schematic overview of the input and output data matrices of clustering and clustering alignment.** Suppose there are $N$ cells, $G$ genes, and $R_K$ runs of clustering results with $K$ clusters. The input data matrix to clustering is $X$ (colored in grey). The output data matrices of clustering are $Q$ and $P$ matrices (colored in blue). Clustering alignment takes in the $Q$ matrices, and outputs the membership matrices of aligned modes; corresponding feature-level matrices of the aligned modes are obtained simultaneously using the alignment derived from the $Q$ matrices (colored in orange).

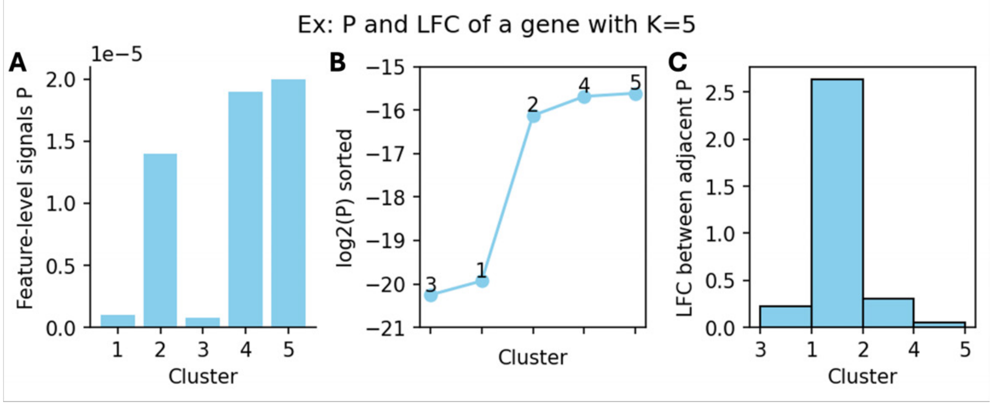


Figure S3: **An illustrative example of how a single gene's clustering profile, $(L^j, \vec{\mathcal{L}}^j)$, captures signals in $P$ for the calculation of feature-level metrics.** **(A)** Relative feature levels of a (hypothetical) gene, corresponding to a row in matrix $P$ with $K = 5$ clusters. The gene has relatively high expression in clusters 2, 4, and 5, and low expression in clusters 1 and 3. **(B)** Base-2 logarithm of sorted entries in this row of $j$, labeled by their indices $\ell_k$. The index vector in clustering profile is $\vec{\mathcal{L}}^j = (3, 1, 2, 4, 5)$. Accordingly, $s_{\text{leLFC}}(j, 1) = (-1)^0 \min\{L_1^j, L_2^j\} = L_1^j$, $s_{\text{leLFC}}(j, 4) = (-1)^1 \min\{L_3^j, L_4^j\} = -L_4^j$, and $s_{\text{sumLFC}}(j) = L_1^j + L_2^j + L_3^j + L_4^j$. **(C)** A bar chart of the LFC vector in clustering profile, $L^j = (L_1^j, L_2^j, L_3^j, L_4^j)$, where left and right edges of each bar $k$ are labeled by $\ell_k$ and $\ell_{k+1}$. The highest bar represents the largest separation gap of log fold change, $L_2^j$, which occurs between clusters 1 and 2 in (B), so $s_{\text{sepLFC}}(j) = L_2^j$ and gene $j$ is separating clusters {3,1} from clusters {2,4,5}—a multi-cluster separation. The smallest separation gap of LFC, $L_4^j$, occurs between clusters 4 and 5.

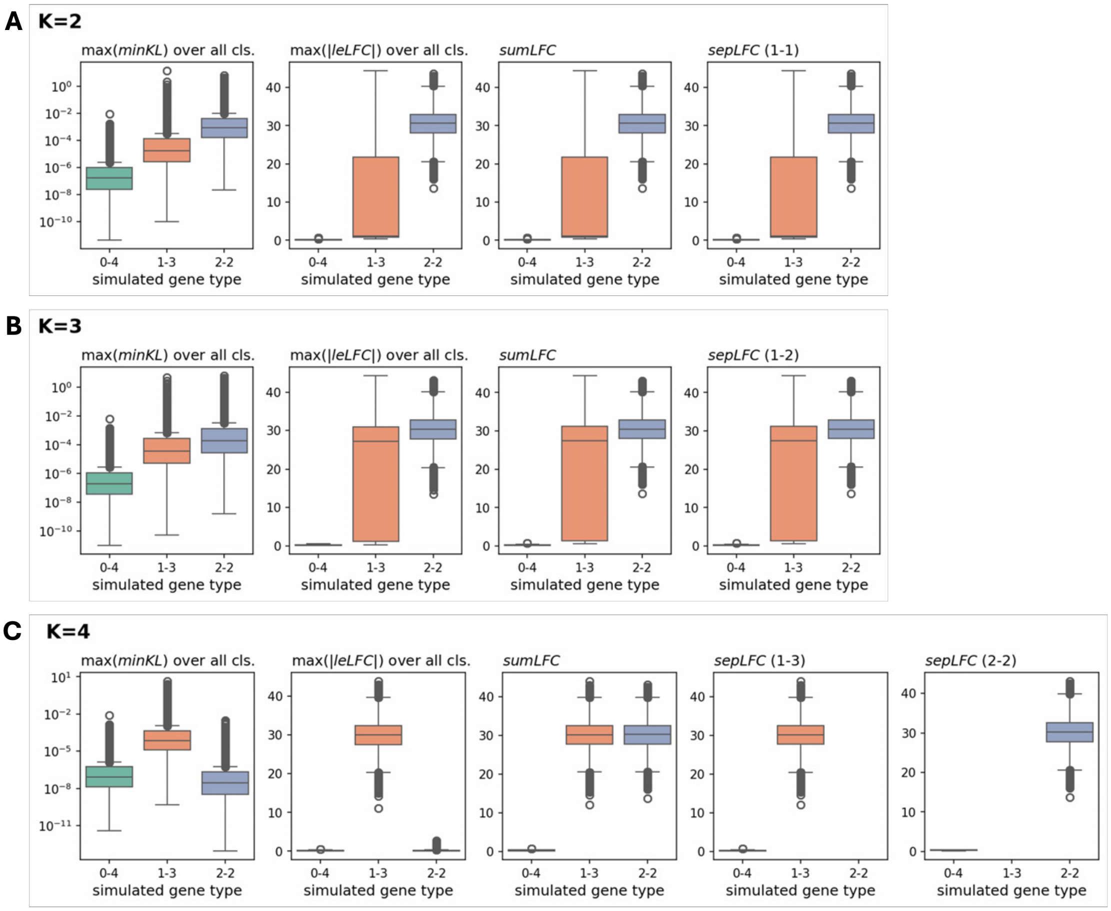


Figure S4: *sepLFC* **more accurately identifies clustering-informative genes than distinctiveness scores across simulation scenarios.** We simulated 3,000 genes under three expression patterns across $K = 4$ clusters ("0-4", "1-3", and "2-2"; the "simulated gene type" detailed in Supplementary Materials S1.6) and computed four scores for each gene: maximum value of $minKL$ (Eq. 3), maximum absolute value of $leLFC$ (Eq. 4), $sumLFC$ (Eq. 7), and $sepLFC$ (Eq. 8). $sepLFC$ is further split based on the separation pattern, that is, how many clusters it separates to the smaller-sized side (a size of 1 up to $\lceil K/2 \rceil$) versus the rest, as labeled in the parentheses following the score name in the subpanel titles. We ran mixed-membership clustering with `FastTopics` (Carbonetto, Sarkar, et al., 2021; Carbonetto, Luo, et al., 2023) at **(A)** $K = 2$, **(B)** $K = 3$, and **(C)** $K = 4$ (10 runs per $K$ on each of 20 simulated matrices; 200 clustering runs per $K$). For clustering with $K = 2$ and $K = 3$, a single $s_{\text{sepLFC}}$ metric is defined For clustering with $K = 4$, there are two $s_{\text{sepLFC}}$ metrics, corresponding to separating one cluster from the other three (1-3), and two clusters from the other two (2-2). Different metrics highlight different aspects of genes' roles in clustering, and patterns are clearest at $K = 4$, where the specified $K$ in clustering matches the true number of clusters in simulation. At $K = 4$, the two distinctiveness scores $s_{\text{minKL}}$ and $s_{\text{leLFC}}$ primarily highlight genes distinctive to a single cluster (1–3) (those with simulated gene type 1-3), whereas $s_{\text{sumLFC}}$ captures genes with strong differences involving one or more clusters (1–3 and 2–2). In contrast, the $s_{\text{sepLFC}}$ metrics identifies both the distinctive genes (simulated gene type 1-3) and genes that are not distinctive for any cluster but contribute to the separation of multiple clusters (simulated gene type 2-2).

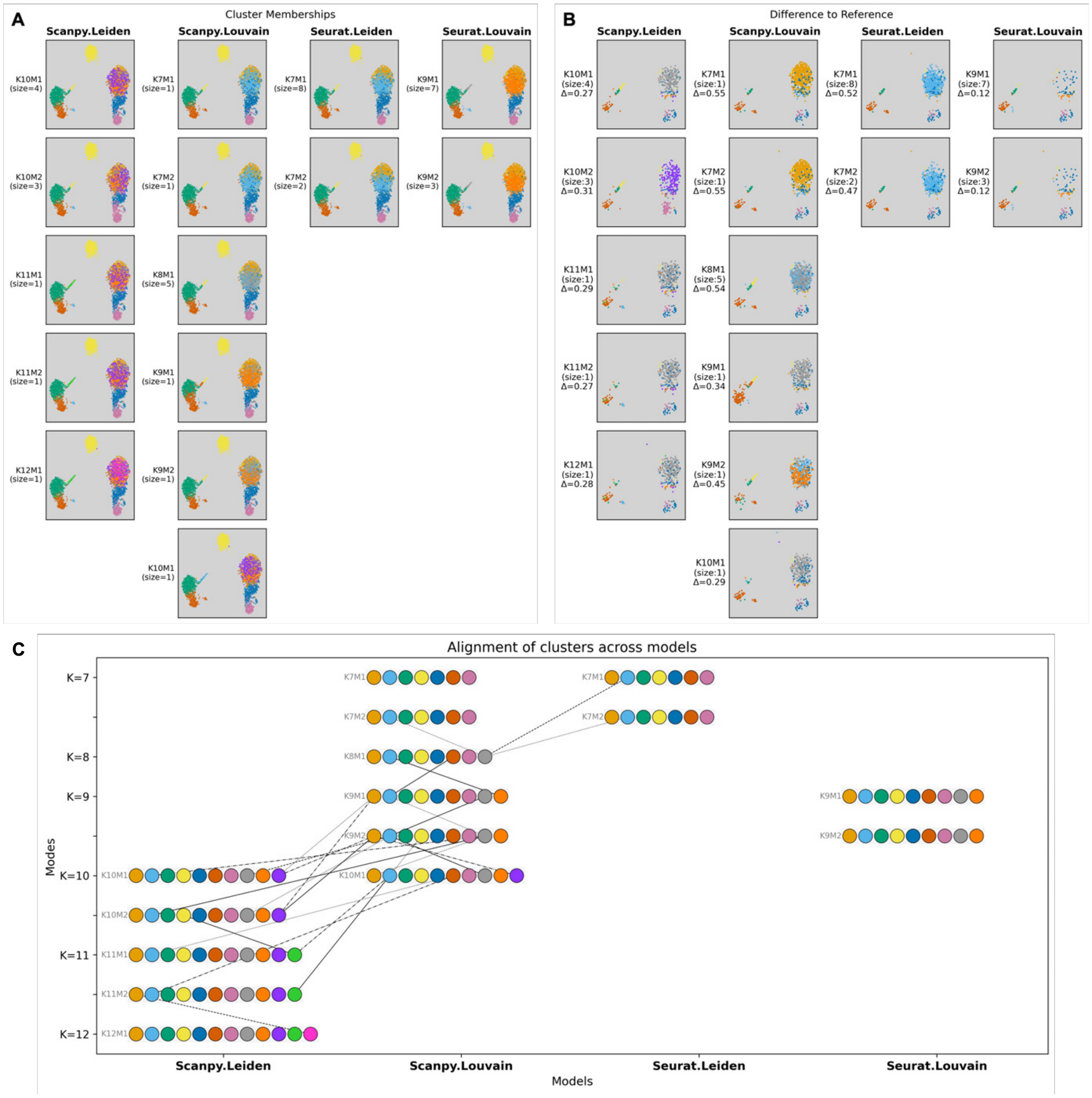


Figure S5: **UMAP plots of all aligned hard-clustering modes for PBMC3k scRNA-seq and their differences to the reference.** We show all modes detected from 10 runs of four models (`Seurat` Leiden, `Seurat` Louvain, `Scanpy` Leiden, and `Scanpy` Louvain), including the selected modes shown in Fig. 1B. Cells are embedded in UMAP space following the `Scanpy` tutorial (Scanpy development team, 2025). **(A)** Cluster memberships for each mode. Modes are labeled by model, number of clusters $K$, and mode size (in parentheses). **(B)** Differences from the `Scanpy`-tutorial reference clustering (Fig. 1A); the average total membership difference $\Delta$ is reported for each mode. **(C)** Cluster-alignment graph in which each node represents a cluster and lines connect aligned clusters across modes (solid and dashed styles alternate for readability). Modes from the same model are arranged in the same column, and modes with the same number of clusters are arranged in the same row. Note that because all modes are aligned jointly by iteratively aligning adjacent $K$ values (e.g., `Seurat` Louvain K9M1 is aligned to $K = 10$ modes from `Scanpy` and then to $K > 10$ ones), modes in non-adjacent columns are not necessarily directly connected (e.g., `Seurat` Leiden vs. `Seurat` Louvain). For clarity, we display only connections between modes in adjacent columns; consequently, some modes appear disconnected.

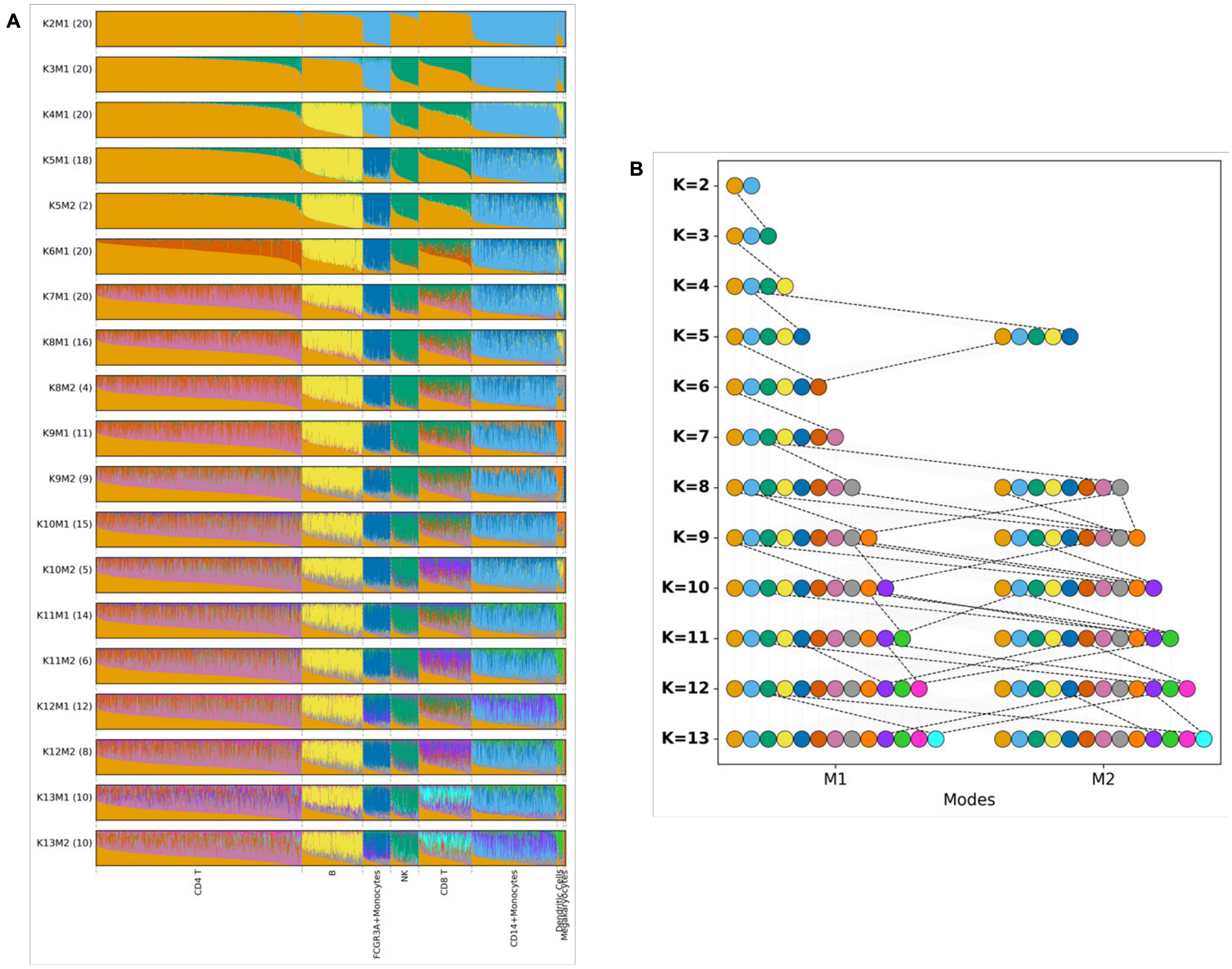


Figure S6: **Structure plots and alignment graph for mixed-membership clustering of PBMC3k scRNA-seq.** We show all modes detected from 20 runs of `FastTopics` (Carbonetto, Sarkar, et al., 2021; Carbonetto, Luo, et al., 2023) for each $K = 2$ to 13. ; labels such as K2M1 (20) denote mode 1 at $K = 2$, with mode size 20 (i.e., 20 runs in this mode). **(A)** Structure plots of cell memberships, shown as stacked bars colored by cluster and ordered by the reference annotations from the tutorial (Fig. 1A). **(B)** Cluster-alignment graph across modes, where nodes denote clusters and edges connect aligned clusters, allowing clusters to be traced to their emergence as $K$ increases.

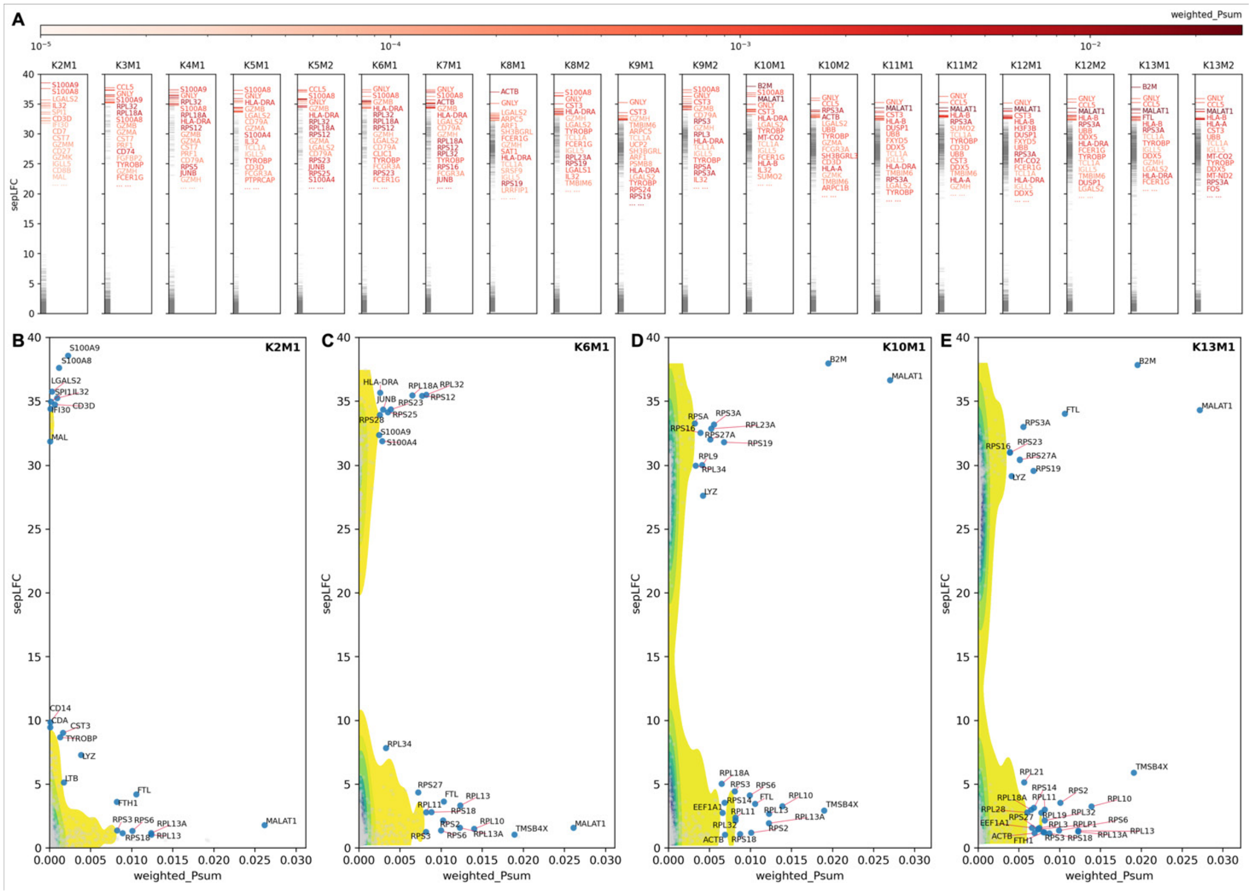

Figure S7: **Joint distributions of feature-level metrics from mixed-membership clustering of PBMC3k scRNA-seq.** We analyze 20 modes detected from 240 `FastTopics` runs (20 runs for each $K$ from 2 to 13); aligned results are shown in Supplementary Fig. S6. This figure follows the layout of Figs. 3A–C. **(A)** Top genes ranked by largest separation gap (*sepLFC* in Eq. 8) for each mode (modes ordered left to right). **(B)-(E)** The joint distribution of *sepLFC* versus weighted $P$ sum (Eq. 9) for all genes in modes K2M1, K6M1, K10M1, and K13M1, shown as contour densities; selected outlier genes are highlighted and labeled.

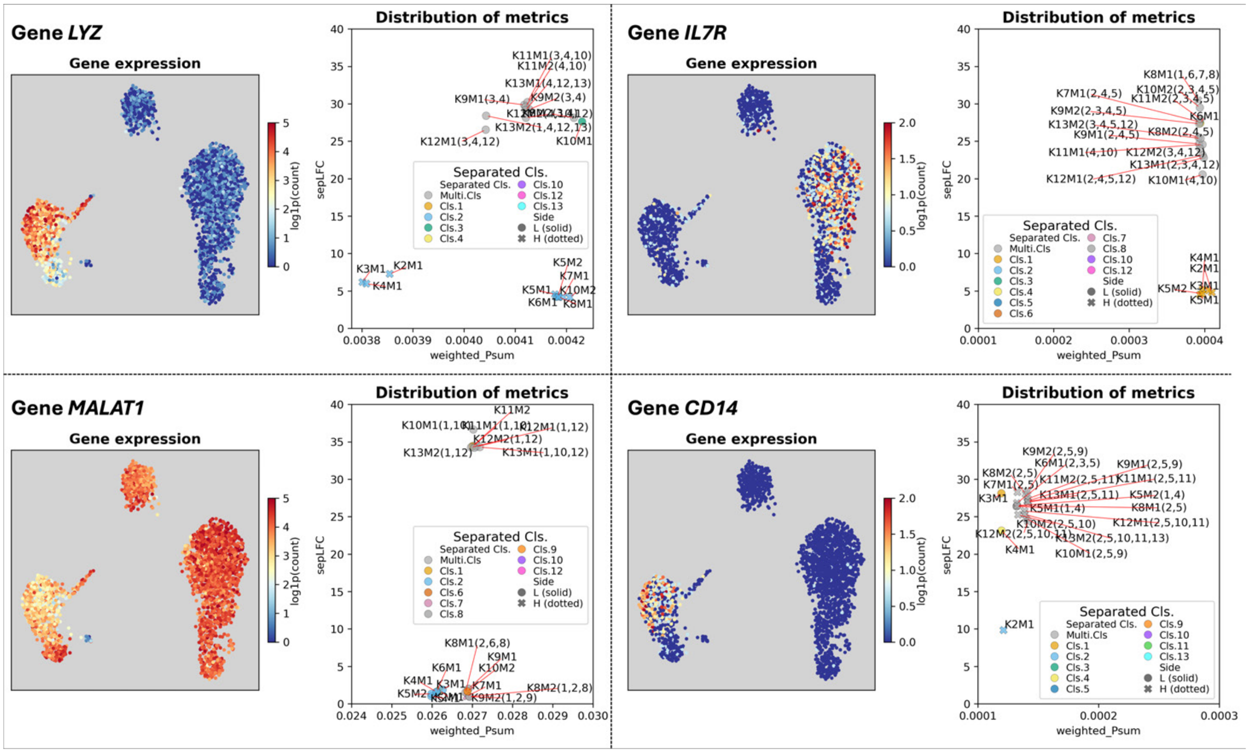


Figure S8: **Expression profiles and feature-metric distributions for candidate clustering-informative genes in PBMC.** This figure follows the layout of Figs. 3D,E. For each gene, the left panel shows log expression counts overlaid on the UMAP, and the right panel shows total relative expression (weighted $P$ sum; $\tilde{p}$ in Eq. 9) versus largest separation gap ($sepLFC$; Eq. 8). Four genes are shown: Lysozyme ($LYZ$), Interleukin-7 Receptor ($IL7R$), metastasis-associated lung adenocarcinoma transcript 1 ($MALAT1$), and cluster of Differentiation 14 ($CD14$).

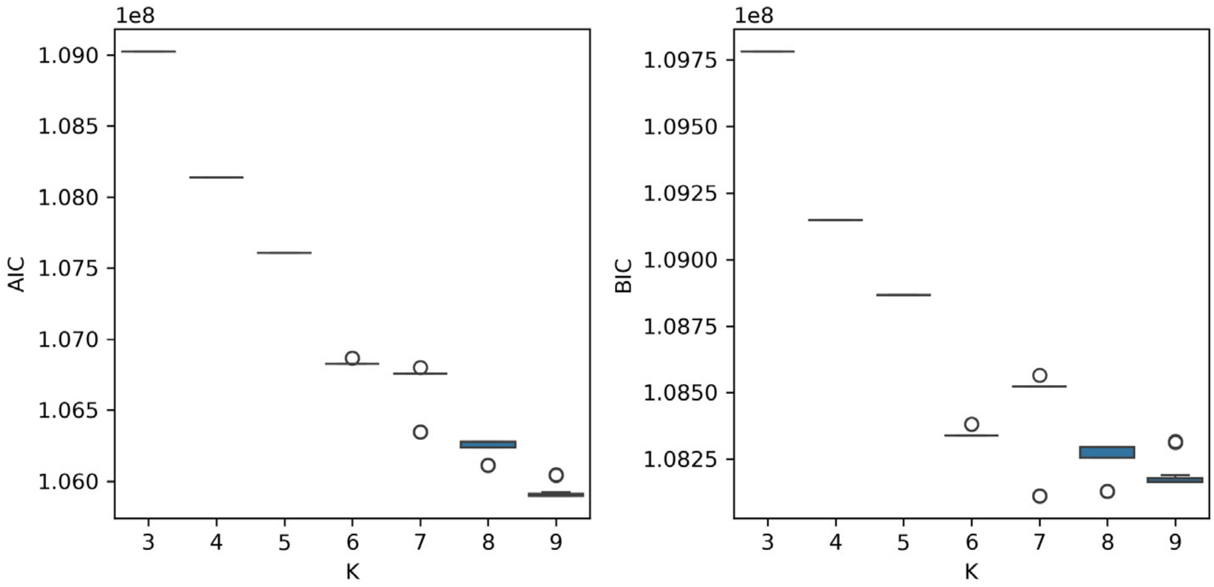


Figure S9: **Information criteria for mixed-membership clustering of human breast cancer ST data ($K$=3–9).** We computed Akaike (AIC; left) and Bayesian (BIC; right) information criteria for each `FastTopics` run (Carbonetto, Sarkar, et al., 2021; Carbonetto, Luo, et al., 2023) using the log-likelihood reported by the model. The values for each $K$ are summarized across 20 runs as box plots. Based on the decreasing AIC/BIC trends with increasing $K$, we choose to focus on mixed-membership clustering results from $K = 3$ to 7 in the main text.

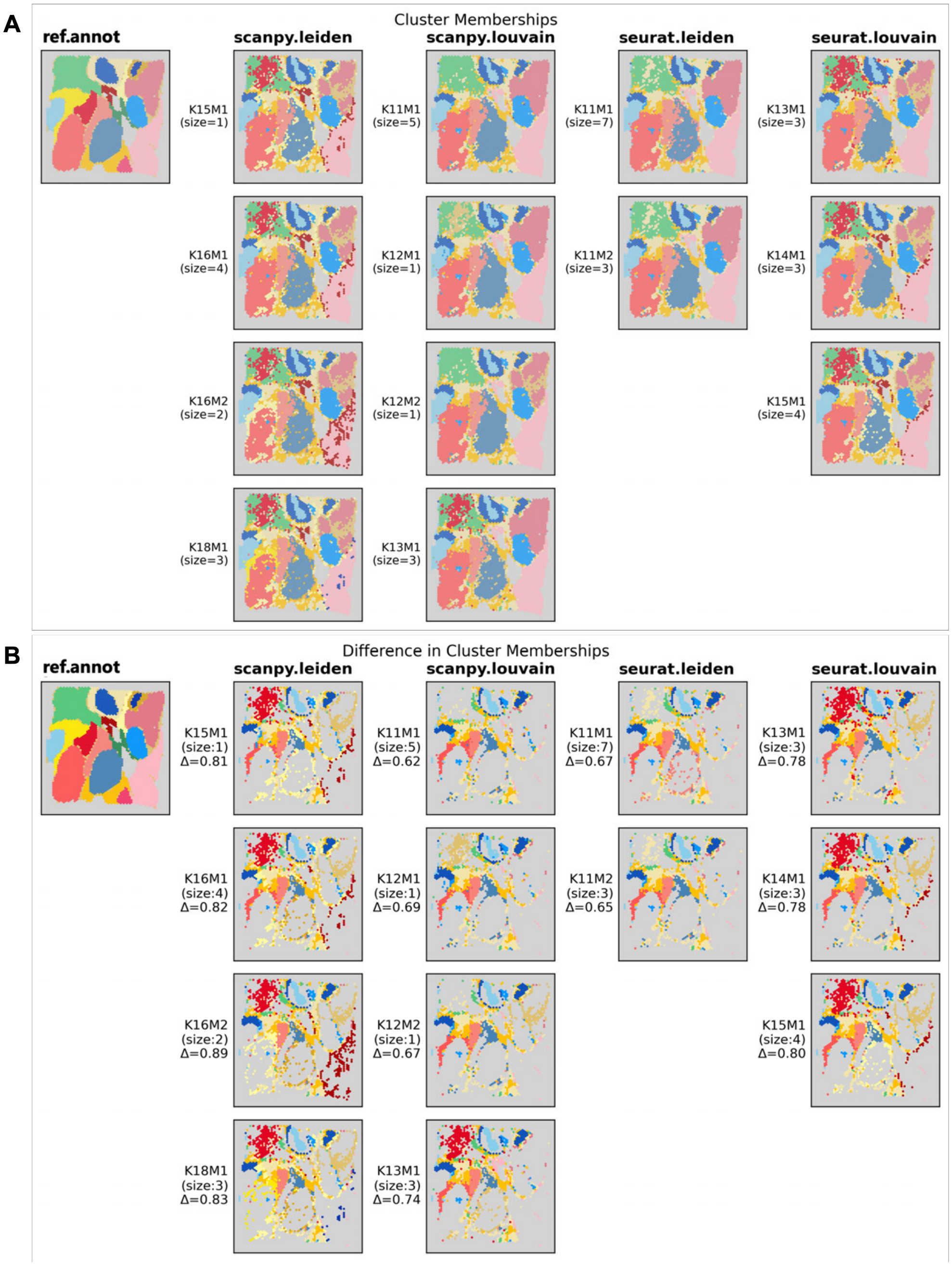


Figure S10: **Full aligned hard-clustering results for human breast cancer ST data.** This figure extends Figs. 2B,C. **(A)** All modes from 10 runs of each hard-clustering model (`Seurat` Leiden/Louvain; `Scanpy` Leiden/Louvain), aligned together with the ground truth using `Clumppling` (X. Liu et al., 2024). Spots are shown in tissue coordinates and colored by aligned cluster labels; mode sizes are reported in each label (Fig. 2B shows only the major modes). **(B)** Spots whose aligned labels disagree with the ground truth, colored by the ground-truth cluster; the average total membership difference $\Delta$ is reported for each mode.

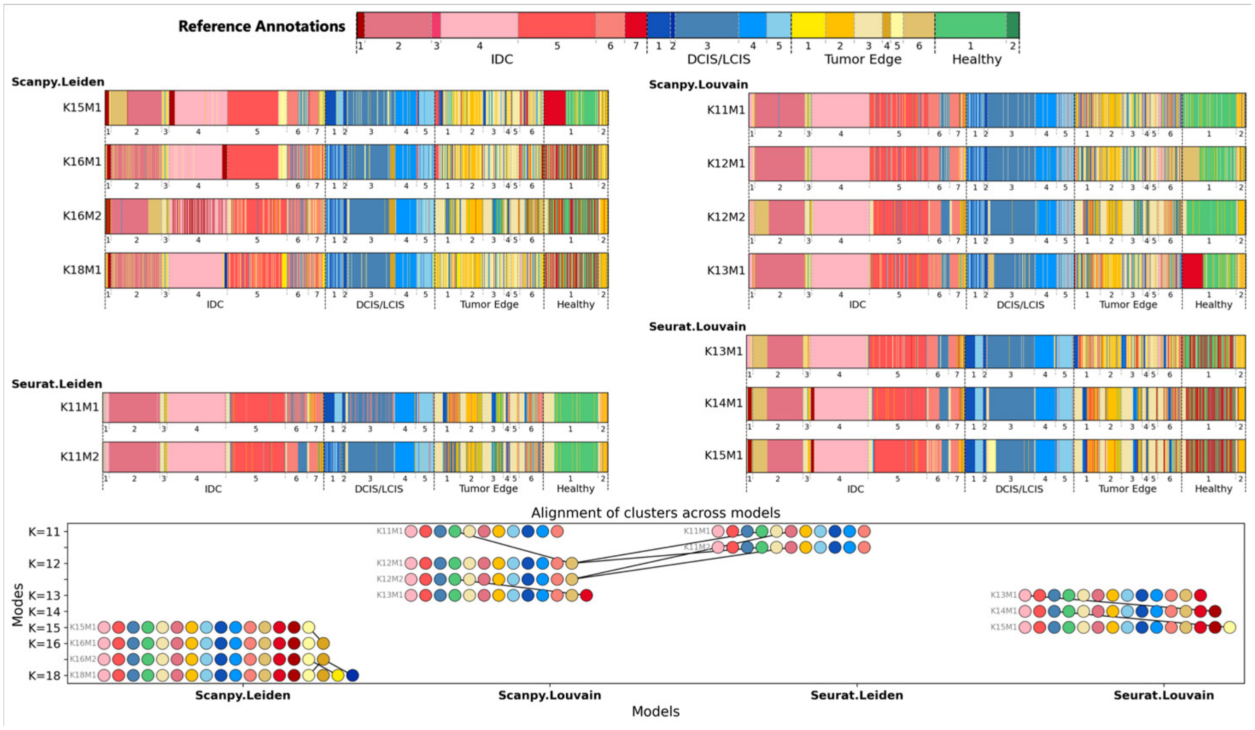


Figure S11: **Structure plots and alignment graph for the full aligned hard-clustering results of human breast cancer ST data.** This figure presents the same modes as Supplementary Fig. S10. **(A)** Structure plots of spot-level cluster assignments (one bar per spot). **(B)** Cluster-alignment graph across modes: nodes denote clusters and edges connect aligned clusters when their colors differ. The layout follows Supplementary Fig. S5C.

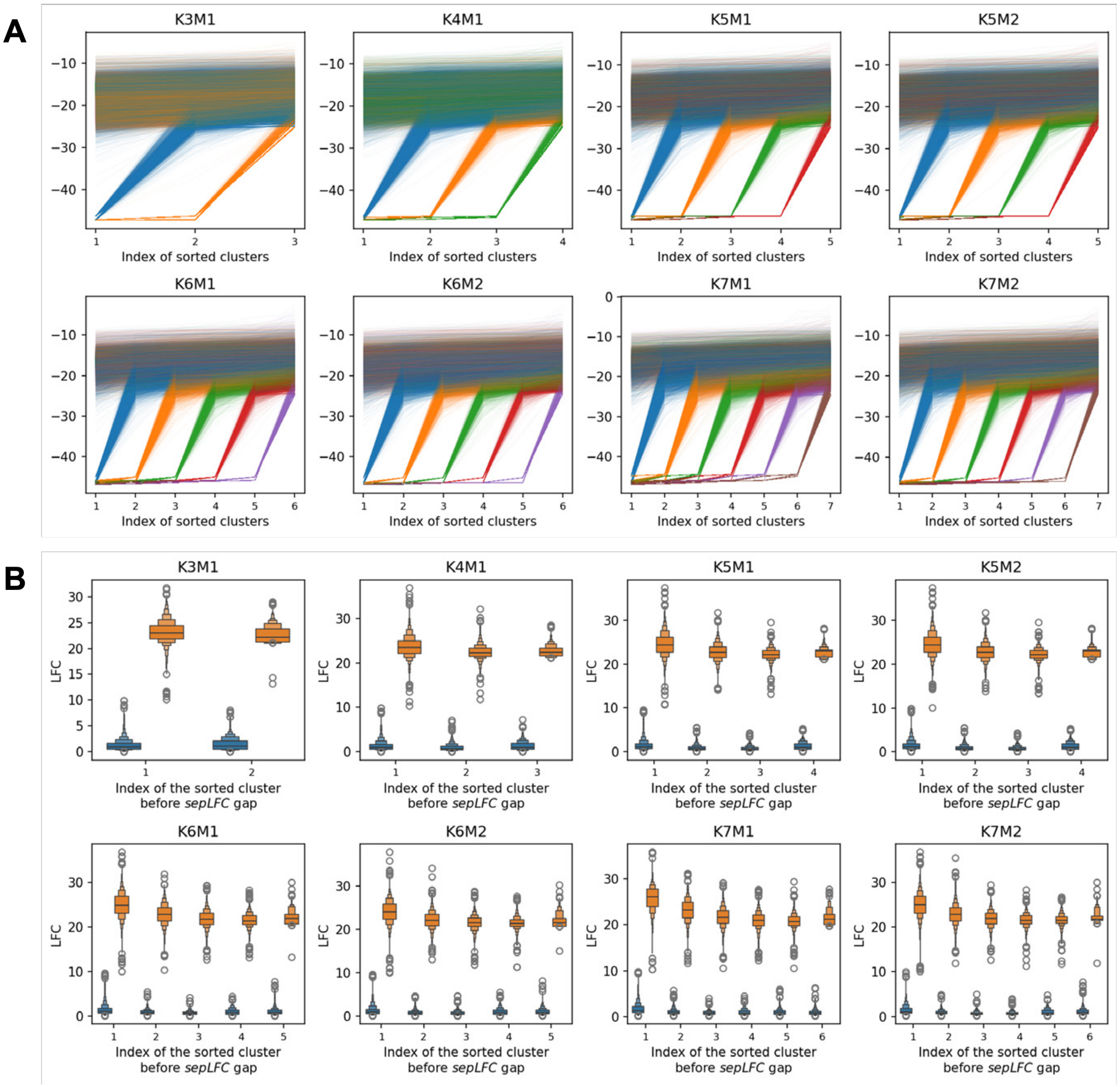


Figure S12: **Distribution of log-$P$ entries and $sepLFC$ or genes in aligned mixed-membership clustering of human breast cancer ST data.** We show selected modes K3M1–K7M2 (Fig. 2E). **(A)** Each line represents a gene, connecting its sorted log-$P$ values across clusters (as in Fig. S3B) and colored by the position of its largest separation gap. Genes either show near-flat profiles (little differentiation) or a sharp jump, yielding large $sepLFC$ (Eq. 8) and strongly separating two cluster sets. **(B)** Using a threshold of 12, we classify genes as high-$sepLFC$ (orange) or low-$sepLFC$ (blue) and group them by the cluster index immediately preceding the largest gap. For each mode, we summarize $sepLFC$ with a letter-value plot (a box-plot variant with finer quantiles), showing clear separation between high- and low-$sepLFC$ genes. Among high-$sepLFC$ genes, gaps that partition highly imbalanced cluster sets (indices near the ends) tend to have slightly larger $sepLFC$ than gaps near the middle of the sorted clusters.

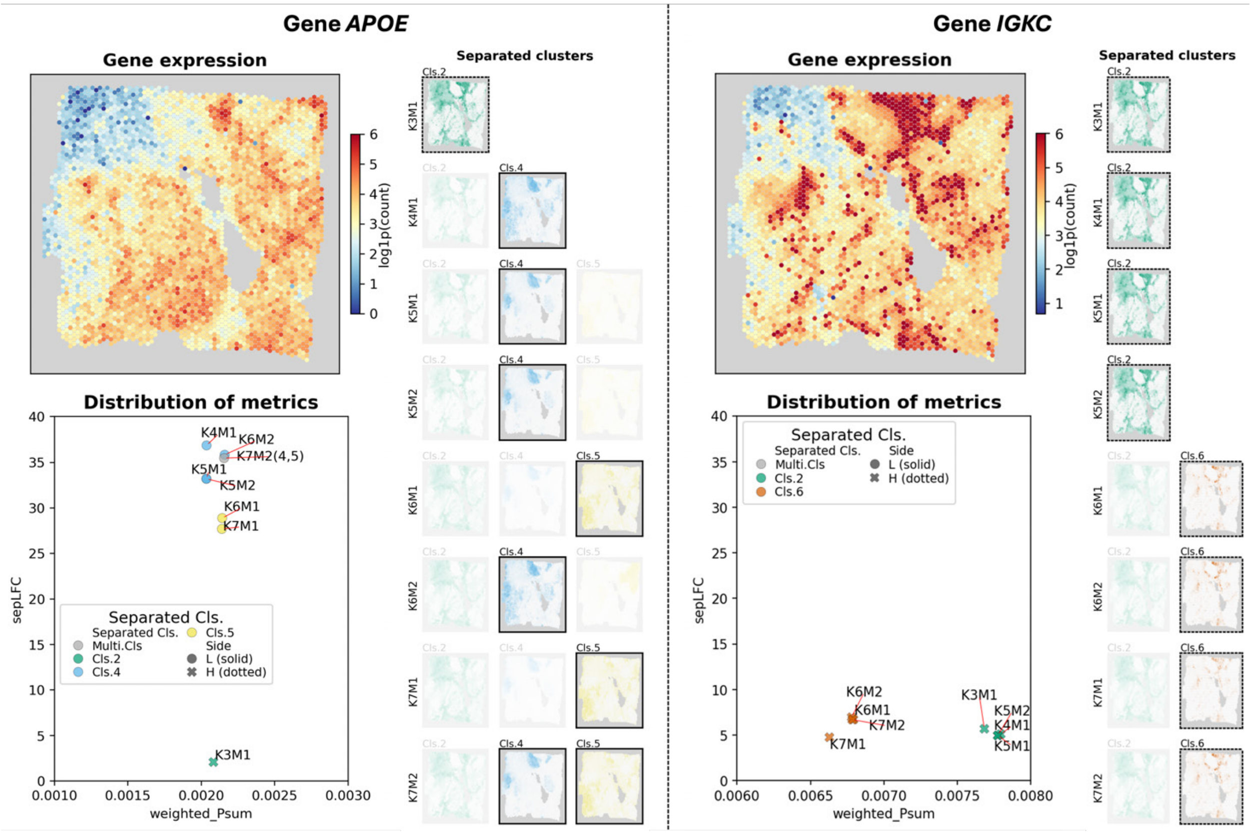


Figure S13: **Clustering profiles for two additional candidate clustering-informative genes in human breast cancer.** We show results for *APOE* and *IGKC*, both of which appear as outliers in the feature-metric distributions (Figs. 3A-C). The layout follows Figs. 3D-F.

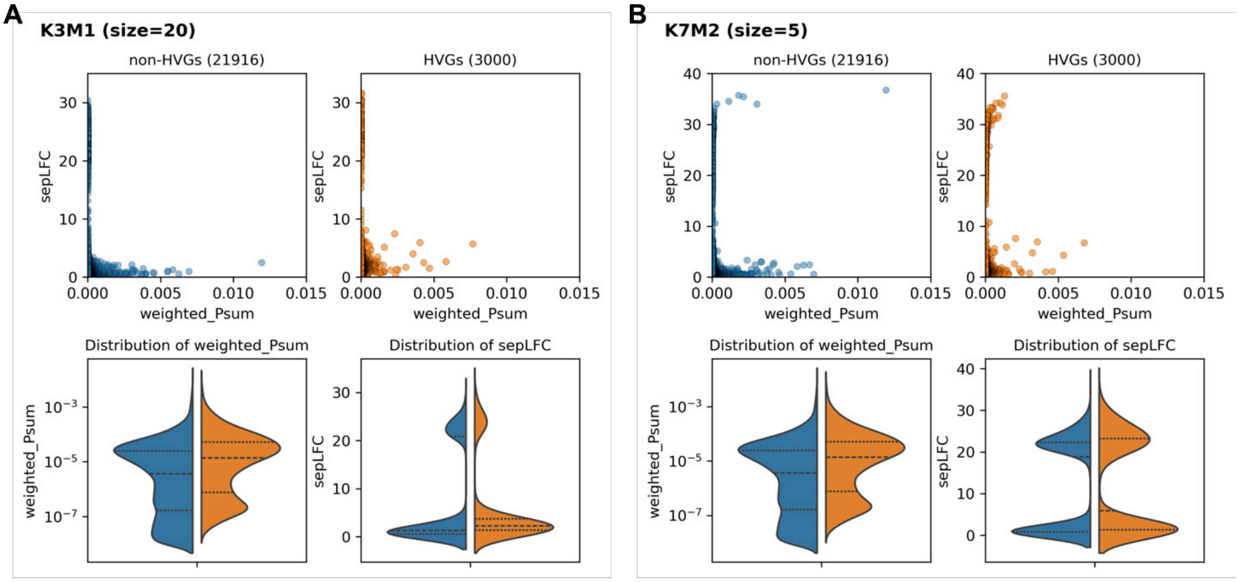


Figure S14: **HVGs versus non-HVGs in mixed-membership clustering of human breast cancer ST data.** We selected 3,000 HVGs from 24,916 genes with nonzero variation using `Scanpy`'s HVG procedure. For modes **(A)** K3M1 and **(B)** K7M2 (mode sizes in parentheses), we plot weighted $P$ sum (Eq. 9) versus $sepLFC$ (Eq. 8) (top) and their marginal distributions (bottom; dashed lines mark quartiles), with HVGs in orange and non-HVGs in blue. Metric distributions are similar between HVGs and non-HVGs, indicating that many non-HVGs are comparably informative for clustering.

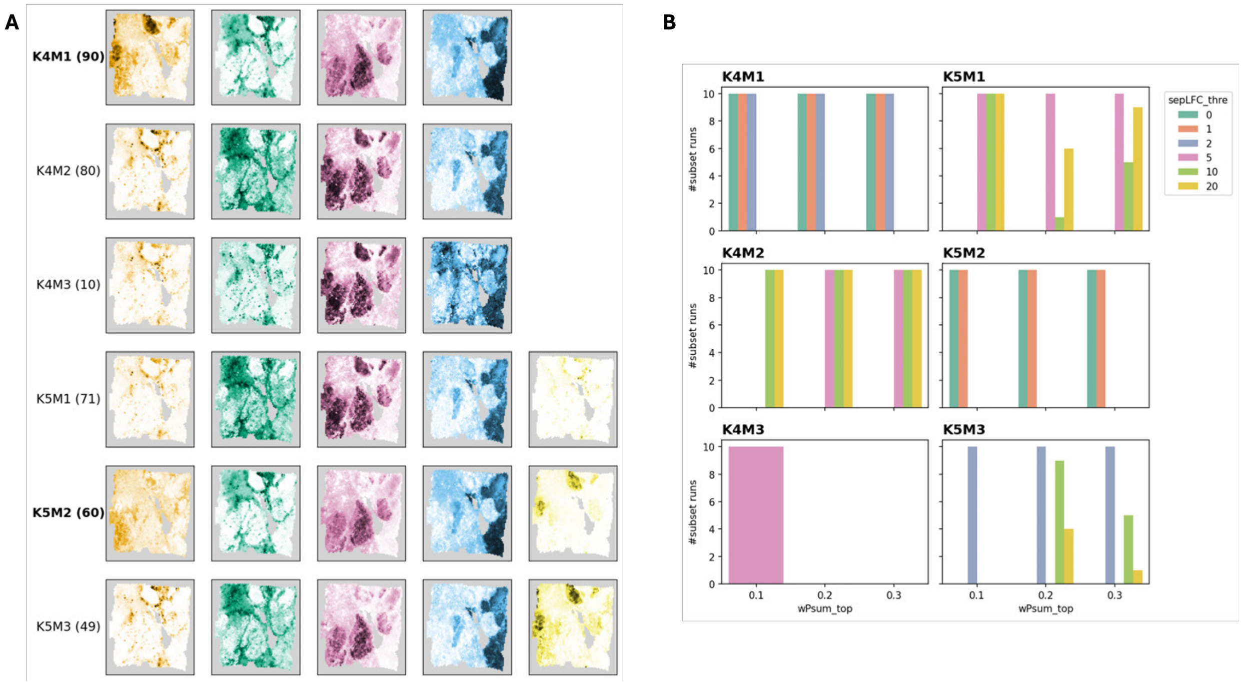


Figure S15: **Mixed-membership clustering of human breast cancer ST data using gene subsets selected by feature metrics.** We constructed 18 gene subsets by combining weighted $P$ sum percentiles (top 10%, 20%, 30%; Eq. 9) with $sepLFC$ thresholds ($> 0, 1, 2, 5, 10, 20$; Eq. 8). For each subset, we ran clustering at $K = 4$ and $K = 5$ (10 runs each; 360 runs total) and aligned all results together with full-gene modes K4M1 and K5M1 (Fig. 2E) using `Clumppling` (X. Liu et al., 2024). **(A)** Spatial membership maps for all modes identified from subset clustering; modes matching full-gene K4M1/K5M1 are bolded. Many runs align to the full-gene modes, while others form distinct modes with clear spatial differentiation. **(B)** Mode composition by subset choice (x-axis: weighted $P$ sum percentile; color: $sepLFC$ threshold). Weighted $P$ sum percentile has little effect, whereas increasing the $sepLFC$ threshold shifts runs into different modes (often for $sepLFC \geq 2$); excluding only low-$sepLFC$ genes leaves clustering largely unchanged, suggesting that genes with very small $sepLFC$ values are typically not clustering-informative even when their weighted $P$ sum is high.

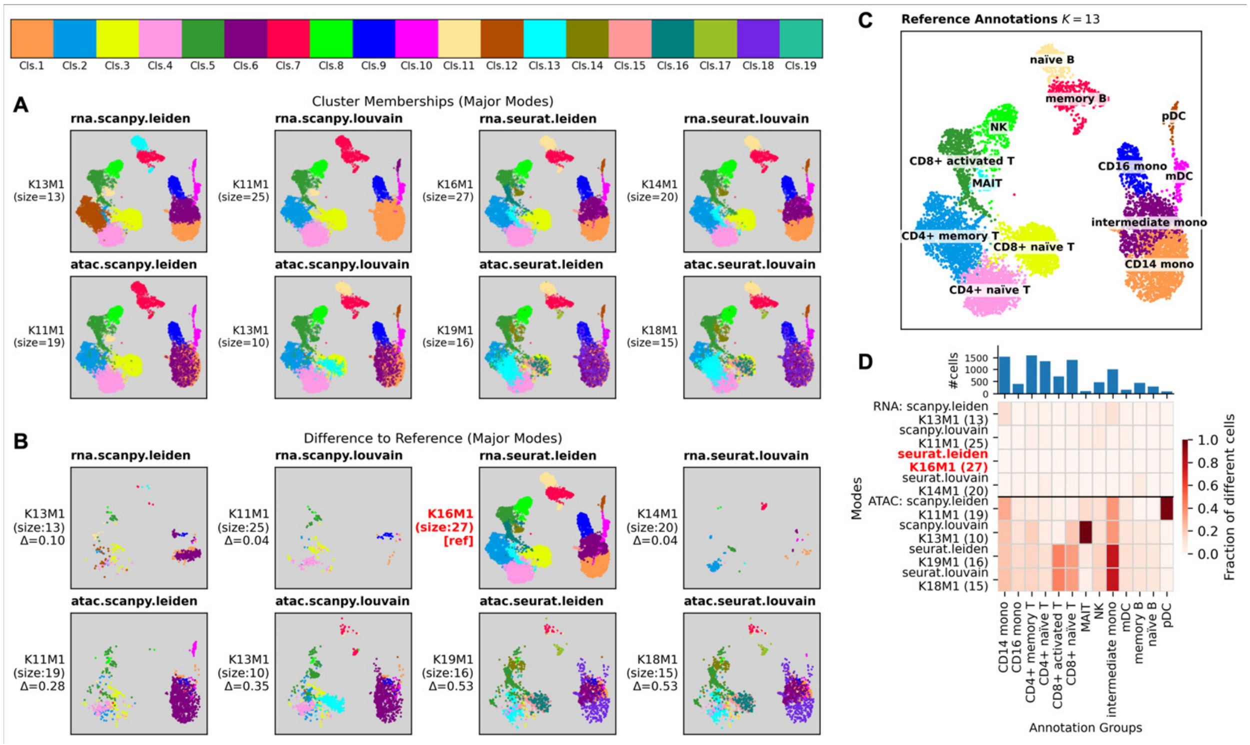


Figure S16: **Clustering alignment reveals disagreement within and across RNA-seq and ATAC-seq clusterings in PBMC10k.** For each modality, we ran 50 replicates of four models (`Scanpy` Louvain/Leiden; `Seurat` Louvain/Leiden) and aligned results across all eight model–modality settings. Major modes span inferred $K$ up to 19; cluster colors are shown in the top colorbar. **(A)** UMAP overlays of cluster memberships for each major mode (model name at top; mode label/size and average total membership difference $\Delta$ at side). UMAPs are generated separately per modality following the `muon` tutorial (Bredikhin, 2025). **(B)** Differences from the RNA-seq `Seurat` Leiden reference mode K10M1 (red); only mismatched cells are colored by their mode-specific labels. **(C)** Reference cell-type annotations from the `muon` tutorial ($K = 13$ cell types) (Bredikhin, 2025). **(D)** Per–cell-type heatmap of the fraction of cells that differ from the reference mode (RNA-seq `Seurat` Leiden K16M1; red), with cell-type sizes shown above. RNA-seq modes are more consistent with the reference than ATAC-seq modes; rare types (e.g., "MAIT", "pDC") show higher fractions but contain fewer cells, and "intermediate mono" shows large discrepancies across all ATAC-seq modes.

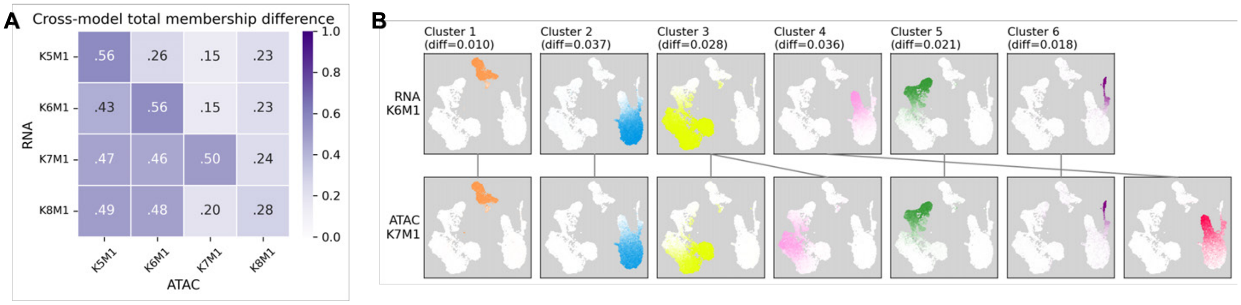


Figure S17: **Membership differences for a cross-omic mode pair.** **(A)** Pairwise average total membership differences between RNA-seq and ATAC-seq modes; RNA-seq K6M1 and ATAC-seq K7M1 have the smallest difference (0.15). **(B)** Aligned memberships for this mode pair, with per-cluster average differences shown at the top; these values sum to the average total membership difference.

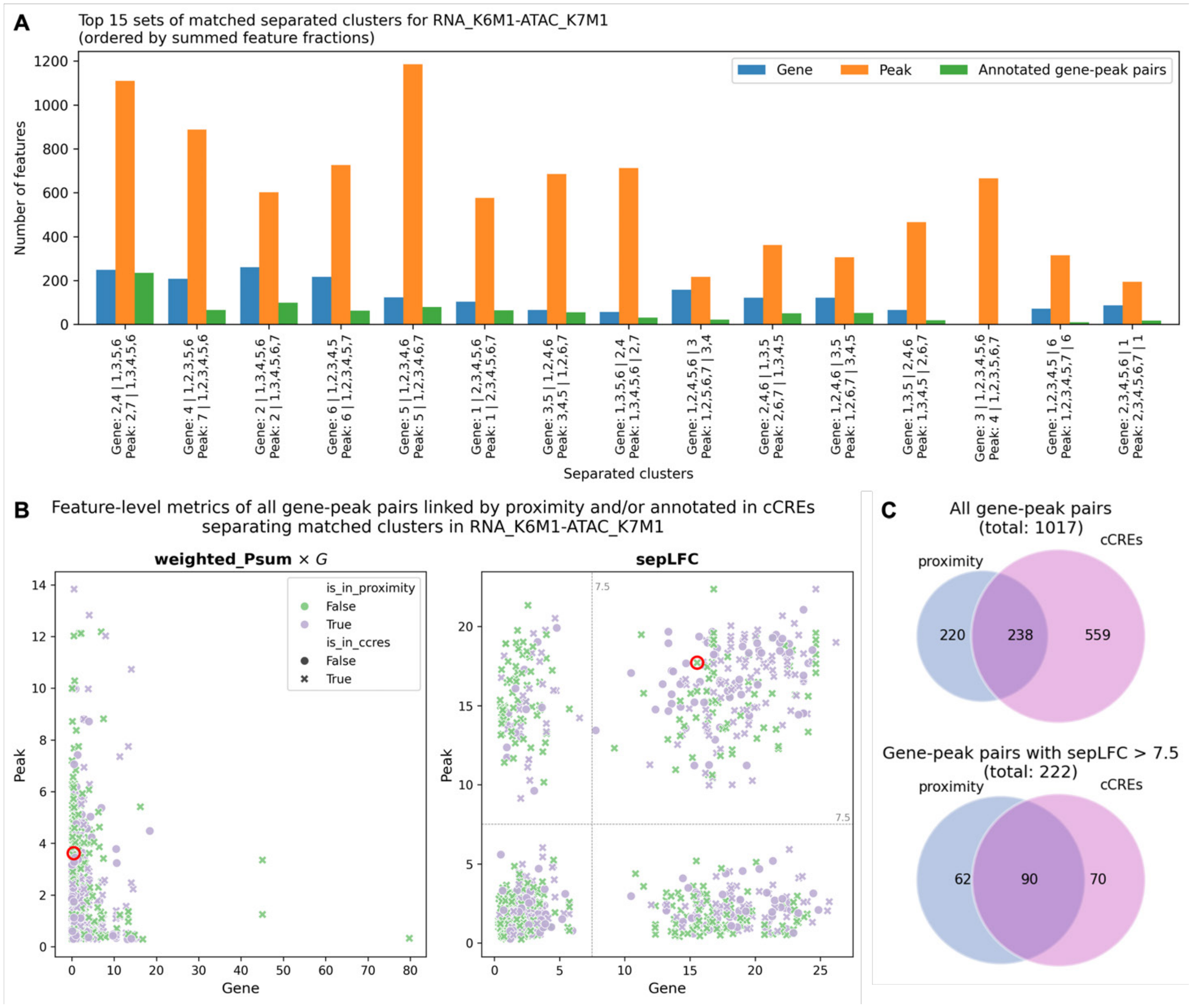


Figure S18: **Gene-peak pairs separating matched clusters in the selected mode pair.** Genes and peaks in the selected cross-omic mode pair, RNA-seq K6M1 and ATAC-seq K7M1 (Supplementary Fig. S17), separate different subsets of clusters, as reflected by their clustering profiles; the separation gap is summarized by the $sepLFC$. **(A)** Top 15 cross-omic cluster-separation patterns shared across omics, ranked by the summed feature fraction, defined as $\frac{\text{number of genes with the separation}}{\text{total number of genes}} + \frac{\text{number of peaks with the separation}}{\text{total number of peaks}}$. Each matched cluster-separation pattern on the x-axis is labeled as “cluster indices (unordered) left of the separation gap | cluster indices (unordered) right of the separation gap” for both genes and peaks. Bars show the number of genes (blue) and peaks (orange) exhibiting each pattern, and the number of annotated gene-peak pairs (green) for which both features share the pattern and the pair is supported by either cCREs or proximity. **(B)** Distribution of weighted $P$ sum and $sepLFC$ for genes versus peaks across all annotated gene-peak pairs. Points are colored/styled according to whether the pair is supported by cCREs, proximity, or both. The weighted $P$ sum is scaled by the number of features $G$. The example gene-peak pair shown in Figs. 4C,D is circled in red. **(C)** Overlap between proximity-linked and cCRE-supported gene-peak pairs shown as a Venn diagram. The top subpanel includes all annotated pairs; the bottom subpanel is restricted to pairs in which both the gene and the peak have $sepLFC > 7.5$ (corresponding to the top-right quadrant of the right subpanel in panel B).

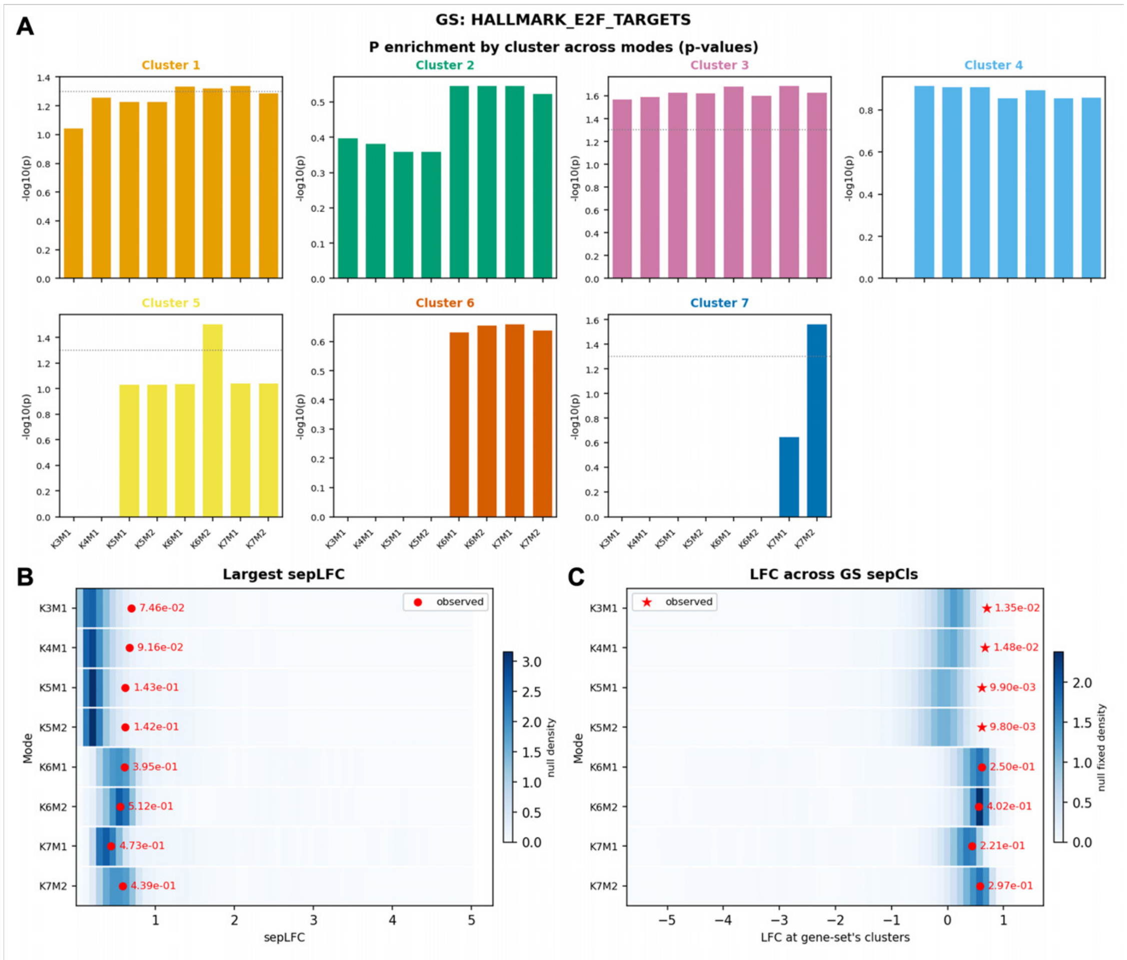


Figure S19: **Gene set enrichment analysis of *Hallmark E2F Targets* on HBC clustering results highlights cluster separations across different values of $K$.** This hallmark gene set "encodes cell cycle-related targets of E2F transcription factors" (Liberzon et al., 2015). Clusters are numbered and colored as in Fig. 2E. Cluster 3, which largely corresponds to two IDC regions, shows consistently enriched relative expression for this gene set. **(A)** Relative-expression ($P$) enrichment, shown as empirical $p$-values for the gene set in each cluster across all modes using bar charts. **(B)** Enrichment of $sepLFC$ in the gene set. The null distribution is shown as a heatmap, and the observed value is marked by a red dot or an asterisk if $p < 0.05$; empirical $p$-values are labeled. For modes with fewer clusters (smaller $K$), the observed value lies away from the null distribution, although not significantly ($p > 0.05$), suggesting that this gene set is enriched for coarse-scale cluster separation. This pattern is not observed in finer-scale clustering modes. **(C)** Enrichment of the log fold change across the two cluster sets in $sepCls$, computed as the LFC between the cluster in $sep\mathcal{H}$ with the lowest relative expression and the cluster in $sep\mathcal{L}$ with the highest relative expression (Eq. 14). The null distribution and observed value are visualized as in panel B. Similarly, the largest separation gap for this gene set is significantly different from the corresponding LFCs in the null gene sets only for the first few modes, which capture coarse-scale cluster differentiation with few clusters.
GS: gene set. $sepCls = (sep\mathcal{L}, sep\mathcal{H})$: two sets of clusters separated by $sepLFC$ for a given mode.

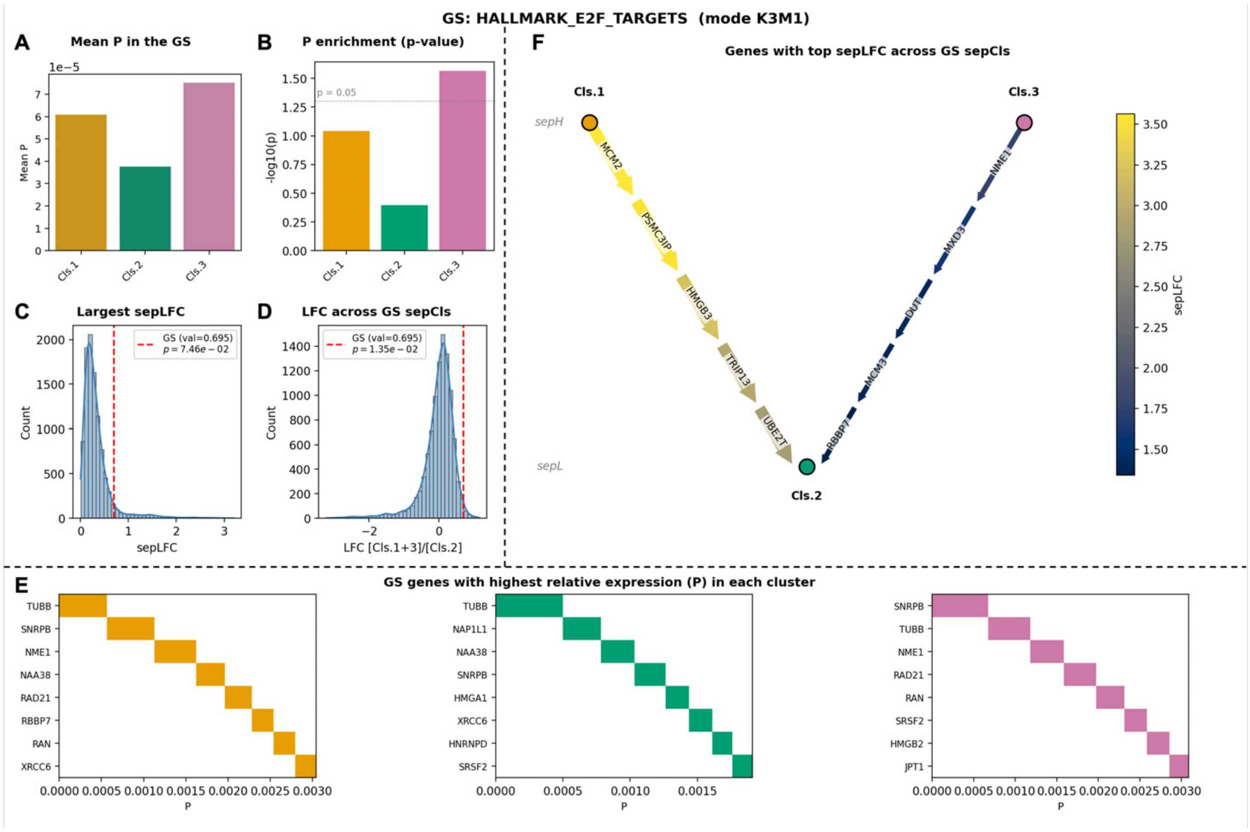


Figure S20: **Gene set analysis of *Hallmark E2F Targets* on HBC clustering mode $K3M1$ (Fig. 2) characterizes gene-level contributions to cluster separation.** Genes in this gene set that drive the separation of Clusters 1 and 3 from Cluster 2 (panel F) differ from those with the highest relative expression in each cluster (panel E). **(A)** Mean relative-expression ($P$) of genes in the GS in each cluster. **(B)** Relative-expression enrichment of the GS in each cluster. The enrichment strength across clusters is consistent with their relative expression levels, with Cluster 3 showing the strongest enrichment and Cluster 2 the weakest. **(C)** Enrichment of the largest separation gap ($sepLFC$) for the GS, assessed against the $sepLFC$ values from null gene sets. This corresponds to the $K3M1$ row in Supplementary Fig. S19B. **(D)** Enrichment of the largest separation gap ($sepLFC$) for the GS, assessed against the LFC across the GS $sepCls$ (Eq. 14). This corresponds to the $K3M1$ row in Supplementary Fig. S19C. **(E)** Top $n = 8$ genes in the GS ranked by their relative expression in each cluster decreasingly. Genes like *TUBB* and *SNRPB* rank high in all clusters. **(F)** Top $n = 5$ genes in the GS ranked by their $sepLFC$ for each each pair of clusters in the separating bipartition $sepCls = (sep\mathcal{L}, sep\mathcal{H})$, denoted in a bipartite graph with clusters in $sep\mathcal{H}$ as top nodes and clusters in $sep\mathcal{L}$ as bottom nodes. An edge connecting two clusters has its width proportional to the sum of the gene-specific $sepLFC$ for that cluster pair. Each edge is further divided into gene-specific segments, with segment lengths proportional to each gene's contribution relative to the total across all top-$n$ genes, and segment color reflecting the gene-specific $sepLFC$ value. Cluster 2 mostly corresponds to tumor-edge and healthy regions, whereas the other two clusters correspond primarily to IDC and DCIS/LCIS. These top genes are therefore the main contributors to this cluster separation. For example, *MCM2* has been shown to be a strong independent prognostic marker in breast cancer (Gonzalez et al., 2003).
GS: gene set. $sepCls = (sep\mathcal{L}, sep\mathcal{H})$: two sets of clusters separated by $sepLFC$ for a given mode.

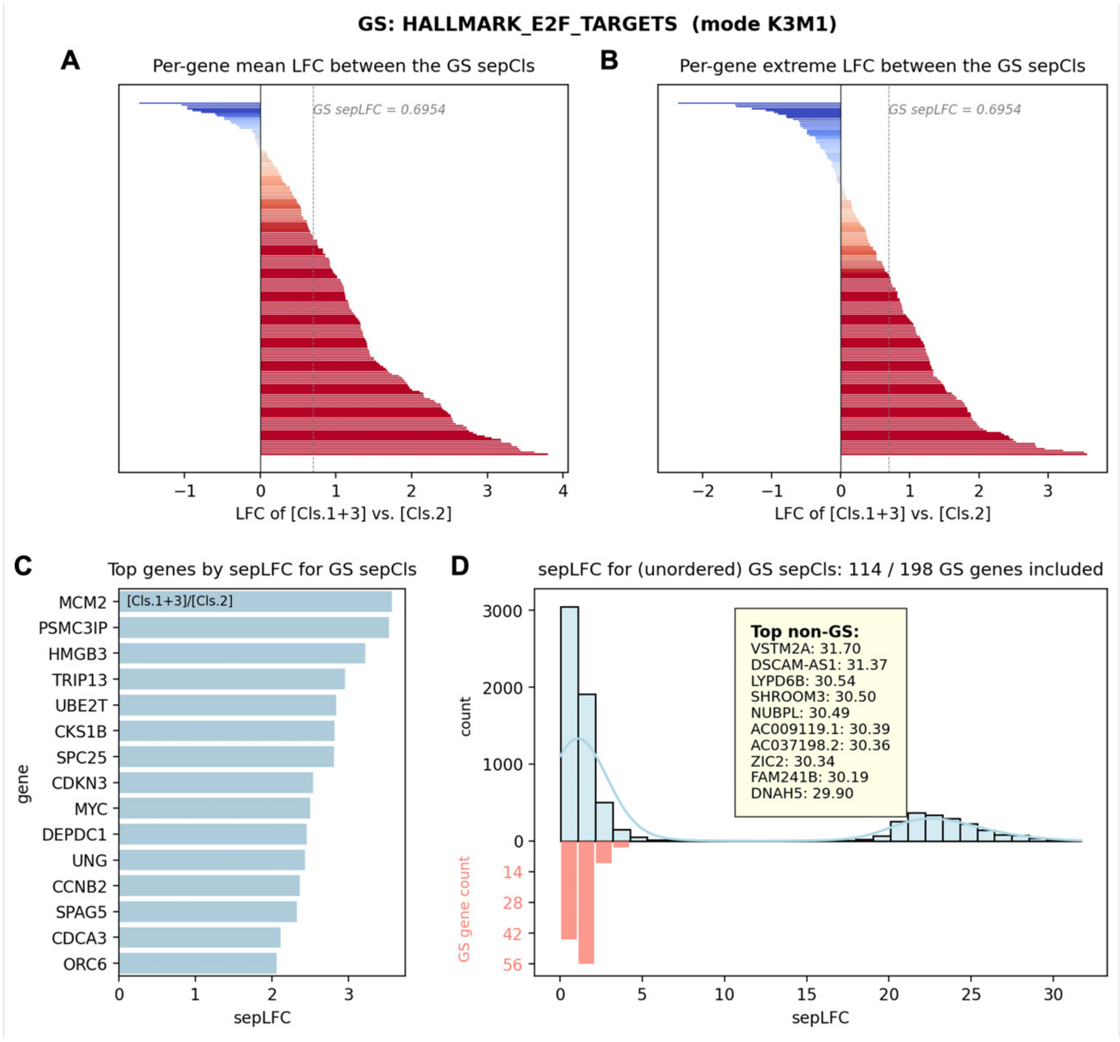


Figure S21: **Contribution of the *Hallmark E2F Targets* gene set to the separation of** $sepCls$ **clusters in HBC clustering mode** $K3M1$ **(Fig. 2) reveals heterogeneous gene-level roles within the gene set.** This visualization provides a more detailed view of the overall separation pattern and the top gene-level contributions to the GS $sepLFC$ summarized in Supplementary Fig. S20F. **(A)** Per-gene mean log fold change across the two cluster sets defined by $sepCls$, computed as the LFC between the mean relative expression over clusters in $sep\mathcal{H}$ and that over clusters in $sep\mathcal{L}$ (Eq. 15). **(B)** Per-gene extreme log fold change across the two cluster sets defined by $sepCls$, computed using the most conservative pair: the lowest relative expression among clusters in $sep\mathcal{H}$ and the highest relative expression among clusters in $sep\mathcal{L}$ (Eq. 16). **(C)** Top genes in the GS, ranked by their gene-specific $sepLFC$ values for the gene set's $sepCls$. **(D)** Distribution of gene-specific $sepLFC$ values for the gene set's $sepCls$ across all genes (top), compared with the subset of genes in the GS (bottom), with the top-ranked non-GS genes labeled. These non-GS genes are not part of the defined biological program and may not be known breast-cancer markers, but they may capture tissue-context variation in the HBC clustering.

GS: gene set. $sepCls = (sep\mathcal{L}, sep\mathcal{H})$: two sets of clusters separated by $sepLFC$ for a given mode.

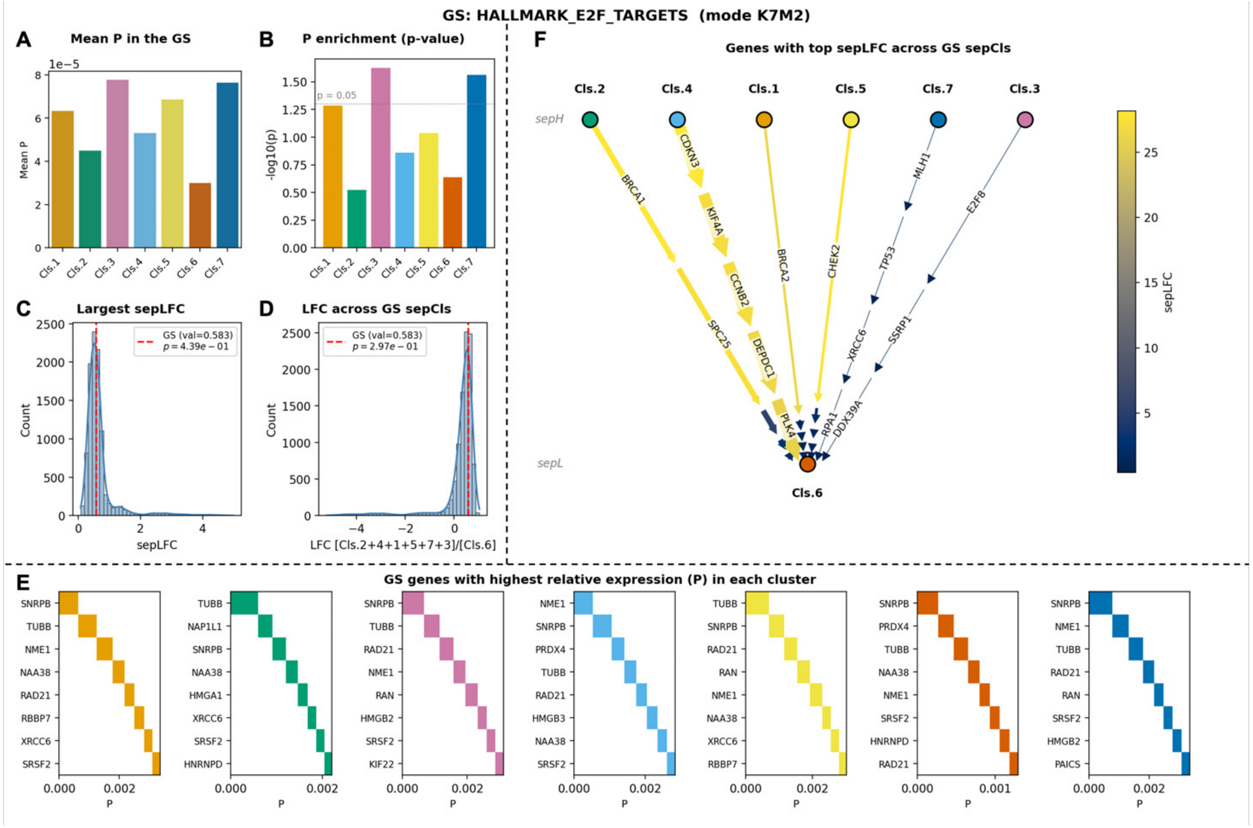


Figure S22: **Gene set analysis of *Hallmark E2F Targets* on HBC clustering mode $K7M2$ (Fig. 2) characterizes gene-level contributions to cluster separation.** **(A)** Mean relative-expression ($P$) of genes in the GS in each cluster. **(B)** Relative-expression enrichment of the GS in each cluster. The enrichment strength across clusters is largely consistent with their relative expression levels, except for Clusters 2 and 6: Cluster 6 has the lowest expression, whereas Cluster 2 shows the weakest enrichment. **(C)** Enrichment of the largest separation gap ($sepLFC$) for the GS, assessed against the $sepLFC$ values from null gene sets. This corresponds to the $K7M2$ row in Supplementary Fig. S19B. **(D)** Enrichment of the largest separation gap ($sepLFC$) for the GS, assessed against the LFC across the GS $sepCls$ (Eq. 14). This corresponds to the $K7M2$ row in Supplementary Fig. S19C. **(E)** Top $n = 8$ genes in the GS ranked by their relative expression in each cluster decreasingly. Genes like *TUBB* and *SNRPB* again rank high in multiple clusters. **(F)** Top $n = 5$ genes in the GS ranked by their $sepLFC$ for each each pair of clusters in the separating bipartition $sepCls = (sep\mathcal{L}, sep\mathcal{H})$, denoted in a bipartite graph with clusters in $sep\mathcal{H}$ as top nodes and clusters in $sep\mathcal{L}$ as bottom nodes. An edge connecting two clusters has its width proportional to the sum of the gene-specific $sepLFC$ for that cluster pair. Each edge is further divided into gene-specific segments, with segment lengths proportional to each gene's contribution relative to the total across all top-$n$ genes, and segment color reflecting the gene-specific $sepLFC$ value. The isolated Cluster 6 primarily corresponds to a subset of tumor-edge spots. Top separating contributors include *BRCA1* and *BRCA2*, which are well-established breast cancer markers.

GS: gene set. $sepCls = (sep\mathcal{L}, sep\mathcal{H})$: two sets of clusters separated by $sepLFC$ for a given mode.

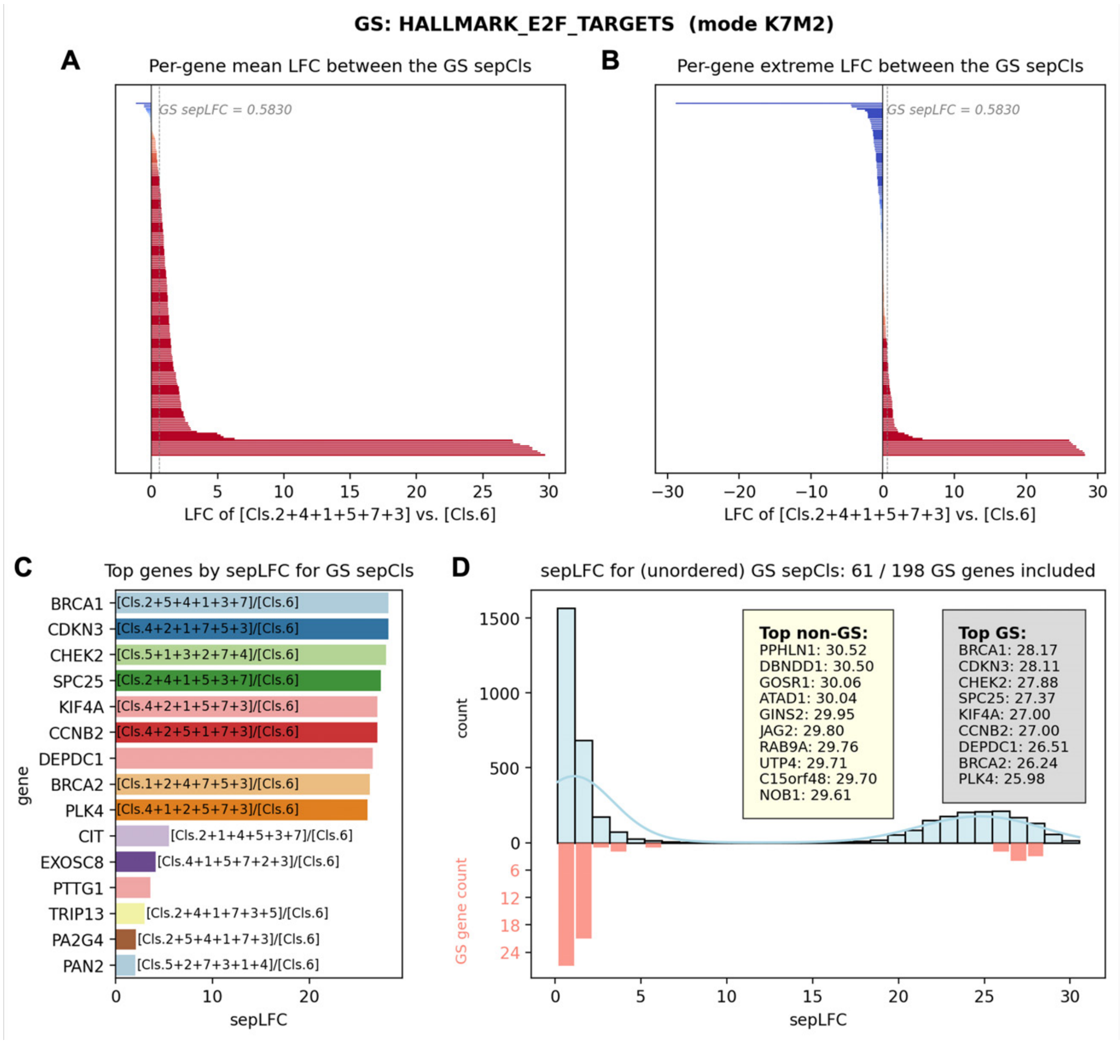


Figure S23: **Contribution of *Hallmark E2F Targets* gene set to the separation of $sepCls$ clusters in HBC clustering mode $K7M2$ (Fig. 2) reveals heterogeneous gene-level roles within the set.** This visualization provides a more detailed view of the overall separation pattern and the top gene-level contributions to the GS $sepLFC$ summarized in Supplementary Fig. S22F. **(A)** Per-gene mean log fold change across the two cluster sets defined by $sepCls$, computed as the LFC between the mean relative expression over clusters in $sep\mathcal{H}$ and that over clusters in $sep\mathcal{L}$ (Eq. 15). **(B)** Per-gene extreme log fold change across the two cluster sets defined by $sepCls$, computed using the most conservative pair: the lowest relative expression among clusters in $sep\mathcal{H}$ and the highest relative expression among clusters in $sep\mathcal{L}$ (Eq. 16). The relatively balanced distribution of positive and negative values explains the small, non-significant deviation of $sepLFC$ for this gene set from the null distribution in the clustering mode (Supplementary Figs. S19B-C and S22C-D). **(C)** Top genes in the GS, ranked by their gene-specific $sepLFC$ values for the gene set's $sepCls$. Distinct gene-specific *ordered* separation patterns are shown in different colors and labeled at their first occurrence. **(D)** Distribution of gene-specific $sepLFC$ values for the gene set's $sepCls$ across all genes (top half), compared with the subset of genes in the GS (bottom half), with the top-ranked non-GS genes labeled. Similar to clustering mode $K3M1$, some non-GS genes show breast-cancer-specific evidence, whereas some likely reflect context-specific transcriptional variation in the HBC clustering.
GS: gene set. $sepCls = (sep\mathcal{L}, sep\mathcal{H})$: two sets of clusters separated by $sepLFC$ for a given mode.
